# Supergene origin and maintenance in Atlantic cod

**DOI:** 10.1101/2021.02.28.433253

**Authors:** Michael Matschiner, Julia Maria Isis Barth, Ole Kristian Tørresen, Bastiaan Star, Helle Tessand Baalsrud, Marine Servane Ono Brieuc, Christophe Pampoulie, Ian Bradbury, Kjetill Sigurd Jakobsen, Sissel Jentoft

## Abstract

Supergenes are sets of genes that are inherited as a single marker and encode complex phenotypes through their joint action. They are identified in an increasing number of organisms, yet their origins and evolution remain enigmatic. In Atlantic cod, four megabase-scale supergenes have been identified and linked to migratory lifestyle and environmental adaptations. Here, we investigate the origin and maintenance of these four supergenes through analysis of whole-genome-sequencing data, including a new long-read-based genome assembly for a non-migratory Atlantic cod individual. We corroborate that chromosomal inversions underlie all four supergenes, and show that they originated at different times between 0.40 and 1.66 million years ago. While we found no evidence for a role of introgression in the origin of the four supergenes, we reveal gene flux between supergene haplotypes with derived and ancestral arrangements, occurring both through gene conversion and double crossover. Our results suggest that supergenes can be maintained over long timescales in the same way as hybridizing species, through the purging of introduced genetic variation.

## Introduction

Many spectacular examples of phenotypical variation within species, such as mimicry patterns in butterflies[1], social organization in ants[2], plumage morphs in birds[3, 4], and floral types in plants[5], are encoded by supergenes — tightly linked sets of genes that control a stable polymorphism in a Mendelian manner[6–9]. Even though supergenes have been known for nearly a century[10], their origin remains a challenging question[9]. The emergence of supergenes requires beneficially interacting mutations in at least two genes and a coinciding reduction of recombination between these genes[7]. As a scenario in which these requirements can be met, recent research has pointed to chromosomal inversions arising in incompletely separated groups, such as interbreeding species or locally adapted populations that exchange migrants[11–13]. In these systems, the beneficial interaction between mutations in different genes can come from their joint adaptation to the same environment, and these mutations can become linked if they are captured by the same inversion[6–9, 14]. This linkage between mutations within inversions is the result of a loop formation that occurs when chromosomes with the derived inverted haplotype pair with chromosomes with the ancestral haplotype arrangement during meiosis. If a single crossover occurs between the two chromosomes within the loop region, the recombinant chromosomes are affected by both duplications and deletions and are therefore unbalanced. The gametes carrying these unbalanced chromosomes are usually lethal, and thus do not contribute to the next generation[15, 16]. When viewed backwards in time, the recombination rate between haplotypes with derived and ancestral arrangements therefore appears reduced. On the other hand, crossovers among two haplotypes that both have the derived arrangement should not affect the viability of gametes. Since most inversions originate just once, in a single individual, the number of individuals in which haplotypes with the derived arrangement can successfully recombine is initially very low, increasing only as the derived arrangement becomes more frequent in the species. The origin of a supergene is therefore expected to be equivalent to a severe bottleneck (down to a single sequence) that affects part of the genome (the inversion region) in a part of the species (the carriers of the derived arrangement)[17].

Once established, the maintenance of supergenes depends on the interaction of selection, drift, gene flow, mutation, and recombination. The derived arrangement can be prevented from fixation — and the supergene can thus remain polymorphic — by frequency-dependent selection, by heterogeneous selection regimes in different populations, or by recessive deleterious mutations that accumulate in the inversion region[7, 11, 13, 18]. As mutations are added over time, the haplotypes with the ancestral and derived arrangements diverge from each other, due to the suppression of recombination between them[18]. Owing to the reduced opportunity for recombination, mildly deleterious mutations are more likely to be fixed inside the inversion region compared to outside and can result in the accumulation of mutational load[18]. When the mutational load becomes high within a supergene, it can lead to fitness decay for individuals carrying two copies of the same arrangement (i.e. homokaryotypes)[19]. The accumulation of deleterious mutations, however, can be counteracted by two processes that allow genetic exchange between the two arrangements[2, 9, 19, 20]: gene conversion and double crossover. Gene conversion is a process in which a homologous sequence is used as template during the repair of a double-strand break, without requiring crossover with that homologous sequence[21, 22]. The fragments copied through gene conversion are short, with lengths on the order of 50–1,000 bp[23, 24]. In vertebrates including fishes, gene conversion is known to increase the GC-content of the involved sequences due to biased repair of A-C and G-T mismatches[22, 25]. Double crossovers occurring within the loop formed by the two chromosomes can lead to the exchange of longer fragments. Either alone or in tandem, the two processes could have the potential to erode differences between supergene haplotypes if their per-site rates are high relative to the mutation rate[26]. However, outside of model systems like *Drosophila* that allow the genetic analysis of crosses produced in the laboratory, the rates of gene conversion and double crossovers are largely unknown.

In Atlantic cod (*Gadus morhua*), genomic regions with tight linkage over 4–17 Megabasepairs (Mbp) and strong differentiation between alternative haplotypes have been identified on linkage groups (LGs) 1, 2, 7, and 12 of the gadMor2 reference genome assembly[27–31]. The alternative haplotypes are associated with different life history strategies [29, 32, 33] and environments[28, 34–37]. One of the strongest of these associations is found between the alternative haplotypes on LG 1 and migratory and stationary Atlantic cod ecotypes[29, 35]. In the Northeast Atlantic, these ecotypes co-occur during the spawning season in March and April along the Norwegian coast, but are separated throughout the rest of the year, with the migratory ecotype — the Northeast Arctic cod (NEAC) — returning to the Barents Sea.[38]. With few exceptions, individuals carrying two copies of one of the haplotypes and heterozygous individuals are migratory, while individuals with two copies of the other haplotype are stationary[29, 39, 40]. Despite the suggestion of a lekking mating system[41], mating between the two ecotypes appears to occur and thus explains the presence of individuals that are heterozygous for the LG 1 haplotypes as well as the very weak separation outside of the four differentiated regions[29]. The differentiated region on LG 1 thus matches the definition of a supergene[6] and is commonly referred to by this term[33–35, 42, 43]. A number of genes from within this supergene have been proposed as candidate genes under selection, including the *Ca6* gene that plays a role in pH-level reduction in the swim bladder and might thus be important for feeding at greater depths in the migratory ecotype. However, the targets of selection among the nearly 800 genes within the supergene remain difficult to identify reliably due to tight linkage between them[32]. Similar to the different frequencies of LG 1 haplotypes between migratory and stationary ecotypes, one of the two alternative haplotypes on LG 2 is far more frequent in Atlantic cod from the Baltic sea compared to the nearest North Atlantic populations and has been suggested to carry genes adapted to low salinity[28, 37]. The alternative haplotypes on LGs 7 and 12 also differ in their frequencies among Atlantic cod populations, with one of the two haplotypes in each case being nearly absent in the Irish and Celtic Sea, which are among the southernmost populations in the Northeast Atlantic, possibly in relation to adaptation to the higher temperatures experienced by these populations[42, 44, 45]. While the ”supergene” status of the haplotypes on LGs 2, 7, and 12 will depend on further investigations of their ecological role, they, too, are commonly referred to by this term in the literature[33–35, 42], and we will do so hereafter.

Chromosomal inversions have long been suspected to be the cause of recombination suppression in the four supergenes in Atlantic cod[46], but this has been confirmed only recently, first for the supergene on LG 1 with the help of detailed linkage maps for that linkage group[32], and then for the three other supergenes through comparison of long-read-based genome assemblies[42]. The age of the supergene on LG 1 was estimated to be around 1.6 million years (Myr) based on the sequence divergence between the two alternative haplotypes and an outgroup, in combination with an assumed divergence time between Atlantic cod and that outgroup[32]. This assumed divergence time, however, may be unreliable, as it was based on a mitochondrial substitution rate[47, 48] that was originally calculated for Caribbean fishes[49]. For the supergenes on LGs 2, 7, and 12, no age estimates have yet been reported. Due to these uncertainties, conclusions about a possible joint origin of all four supergenes have necessarily remained highly speculative. The role of introgression — genetic exchange through hybridisation — in the origin of the Atlantic cod’s supergenes has so far also been uncertain. While introgression among codfishes (subfamily Gadinae) has been supported by one former study based on genome-scale sequence data[50], these results were affected by the use of an incorrectly labelled specimen, susceptibility to reference bias, and possibly incorrect outgroup choice in the application of *D*-statistics (Supplementary Note 1), and thus remain inconclusive regarding the occurrence of introgression.

Here, we investigate the origin and the maintenance of supergenes in Atlantic cod as follows: We generate a new long-read-based genome assembly for a stationary Atlantic cod individual from northern Norway as a complement to the existing genome assembly for a migratory individual belonging to the NEAC population (gadMor2[31]). Importantly, these two assemblies carry alternative haplotypes at each of the four supergenes. Through comparison of the two assemblies with each other and with an outgroup assembly, we corroborate that inversions are the cause of recombination suppression for each supergene, pinpoint the chromosomal boundaries of supergenes, and identify ancestral and derived arrangements. Using Bayesian time-calibrated phylogenetic analyses of newly generated and previously available genomic data, we identify traces of introgression among closely related codfishes and show that at least some of the supergenes originated at different times. Through demographic analyses, we find signatures of past bottlenecks associated with the origin of supergenes, and by applying *D*-statistics and sliding-window phylogenetic inference, we detect the occurrence of genetic exchange between haplotypes, both through gene conversion and double crossovers. Our results suggest that the long-term existence of supergenes may depend on genetic exchange between haplotypes to counter the accumulation of mutation load, and on selection acting on the exchanged sequences to maintain the separation of the two haplotypes.

## Results

### A genome assembly for a stationary *Gadus morhua* individual

To allow a comparison of genome architecture between migratory and stationary *Gadus morhua*, we performed PacBio and Illumina sequencing for a stationary *Gadus morhua* indivdual sampled in northern Norway, at the Lofoten islands (Fig. 1a). The resulting genome assembly (gadMor Stat) consisted of 6,961 contigs with a contig N50 length of 121,508 bp and had a size of 565,431,517 bp, corresponding to approximately 87% of the estimated size of the *Gadus morhua* genome[31]. The assembly contained 3,061 (84.1%) complete and 3,020 (83.0%) complete and single-copy genes out of 3,640 conserved genes in BUSCO’s[51] Actinopterygii gene set (see Supplementary Table 1 for further assembly statistics). When aligned to the gadMor2 reference genome assembly[31] (representing a migratory individual), the gadMor_Stat assembly was highly similar on almost all gadMor2 linkage groups, with a pairwise sequence divergence of 0.0040–0.0053 substitutions per site between the two assemblies. The exception to this were the four supergenes on LGs 1, 2, 7, and 12, that all showed an elevated sequence divergence of 0.0066–0.0129 substitutions per site between the two assemblies. This confirmed that the gadMor Stat and the gadMor2 assemblies carried alternative supergene haplotypes on all four linkage groups (Supplementary Table 2). To determine the chromosomal boundaries of the regions of tight linkage associated with the supergenes, we investigated linkage disequilibrium (LD) on LGs 1, 2, 7, and 12 with a dataset of single-nucleotide polymorphisms (SNPs) for 100 *Gadus morhua* individuals. By quantifying the strength of linkage per SNP as the sum of the distances (in bp) with which the SNP is strongly linked (*R*^2^ *>* 0.8), we identified sharp declines of linkage marking the boundaries of all four supergenes (Fig. 1b, Table 1), as expected under the assumption of large-scale chromosomal inversions[52].

**Table 1.** Tight linkage and chromosomal inversions in supergene regions in *Gadus morhua*.

| LG | High-LD region |  | Inversion region |  | Arrangement |  |
| --- | --- | --- | --- | --- | --- | --- |
|  | Beginning | End | Beginning | End | gadMor2 | gadMor_Stat |
| 1 | 9,114,741 | 26,192,489 | 9,128,372–9,130,274 | ~26,100,000<sup>1</sup> | derived | ancestral |
| 2 | 18,489,307 | 24,050,282 | ~18,490,000<sup>2</sup> | 24,054,399–24,054,406<sup>3</sup> | ancestral | derived |
| 7 | 13,606,502 | 23,016,726 | 13,651,003–13,652,432 | 23,002,424–23,043,967 | derived | ancestral |
| 12 | 589,105 | 13,631,347 | 607,782–662,878 | 13,386,293–13,614,908 | ancestral | derived |

<sup>1</sup>Comparison with the gadMor3 assembly [42] (Supplementary Figure 2) suggests that the actual end of the inversion region is misplaced in the gadMor2 assembly, between positions 18,890,477 and 18,900,044, and that the region from position ~16,800,000 and ~18,900,000 in the gadMor2 assembly is instead located after the current position ~26,100,000.
<sup>2</sup>Due to repetitive sequences at the beginning of the inversion region, contigs mapping inside and outside of the region overlap between positions 18,487,151 and 18,494,225.
<sup>3</sup>This is the end of the linkage group in the gadMor2 assembly; however, comparison with the gadMor3 assembly suggests that the region from position ~22,600,000 to ~23,700,000 in the gadMor2 assembly is incorrectly placed and instead located at the end of the linkage group.

**Fig. 1.**
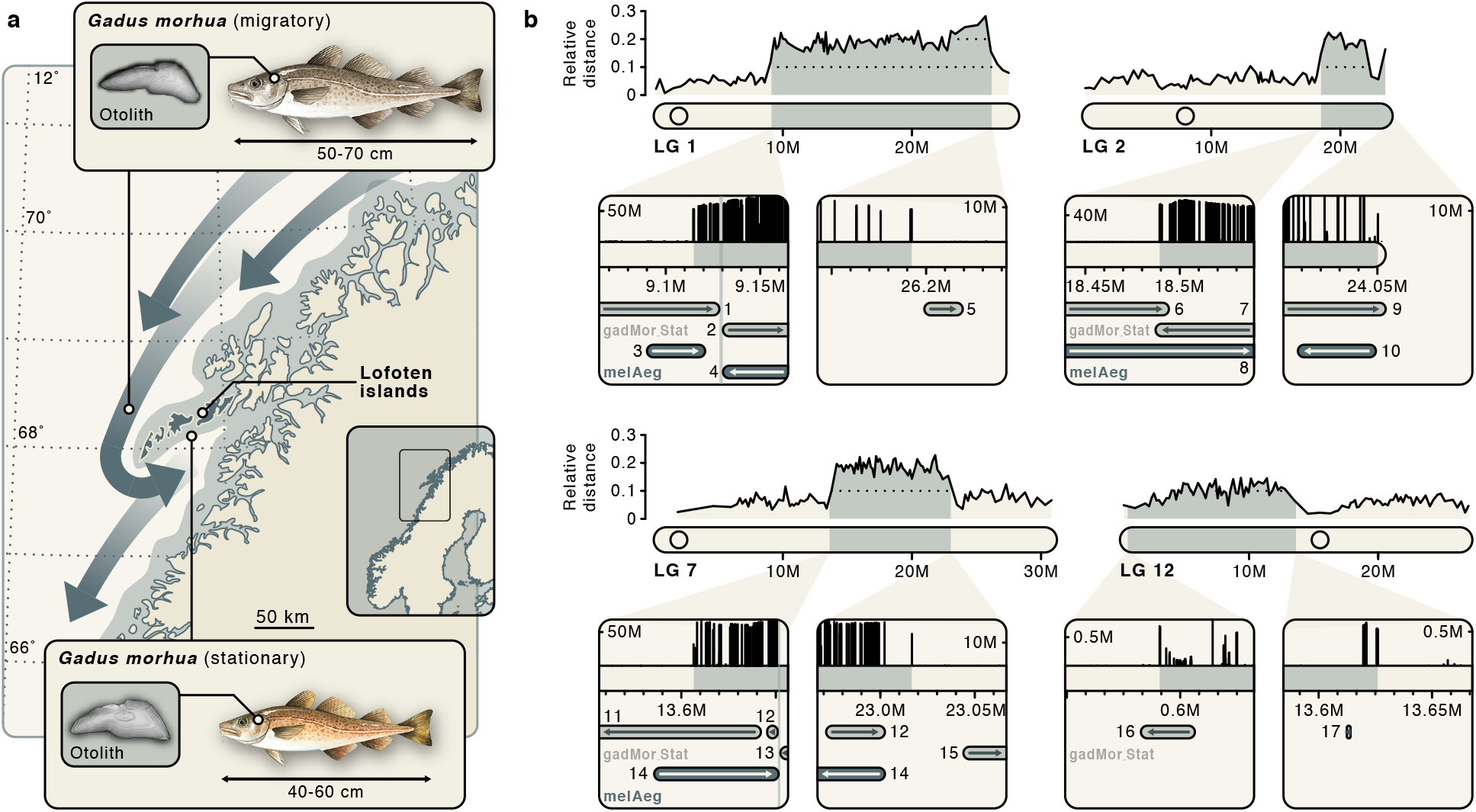
Four supergenes associated with megabase-scale chromosomal inversions in *Gadus morhua*. **a** Migratory and stationary *Gadus morhua* seasonally co-occur along the coast of northern Norway and differ in total length and otolith measurements[38, 55]. The distribution of stationary *Gadus morhua* is shaded in grey whereas the seasonal movements of migratory *Gadus morhua* are indicated with dark grey arrows. **b** Pairwise genetic distance (in substitutions per site) between the gadMor2 and gadMor Stat assemblies, relative to the genetic distance to an assembly for *Melanogrammus aeglefinus* (melAeg)[56] in a three-way whole-genome alignment. The alignment coordinates are according to the gadMor2 assembly. The four LGs 1, 2, 7, and 12 are shown as rounded horizontal bars, on which circles indicate the approximate centromere positions [42]. Supergene regions are shaded in grey, and the beginning and end of each of these regions are shown in more detail in the insets below each linkage group. Each of these insets focuses on a section of 100 kbp around a supergene’s beginning or end. Shown in black above the bar representing that section is a per-SNP measure of linkage disequilibrium (LD), calculated as the sum of the distances between SNPs in high linkage (*R*^2^ *>* 0.8). Based on this measure, the grey shading on the bar illustrates the beginning or the end of high LD. Drawn below the scale bar are contigs of the gadMor Stat and melAeg assemblies, in light grey and dark grey, respectively, that align well to the shown sections. The arrows indicate the alignment orientation of contigs (forward or reverse complement), and contigs are labelled with numbers as in Supplementary Table 3. In the first insets for LGs 1 and 7, vertical bars indicate inferred inversion breakpoints, which are found up to 45 kbp (Table 1) after the onset of high LD. Fish drawings by Alexandra Viertler; otolith images by Côme Denechaud.

The presence of megabase-scale inversions on each of the four linkage groups was further supported by alignments of contigs from the gadMor_Stat assembly to the gadMor2 assembly, as we identified several contigs with split alignments, of which one part mapped unambiguously near the beginning and another mapped near the end of a supergene (Supplementary Table 3). The positions of split contig alignments allowed us to pinpoint the inversion breakpoints on the four linkage groups with varying precision (Table 1). The most informative alignments were those near the beginnings of the supergenes on LGs 1 and 7, which in both cases placed the breakpoints within a window of approximately 2 kbp. As also reported for inversions in *Drosophila*[53], this precise placement of the inversion breakpoints revealed that they do not match the positions of LD onset exactly, but that they were located up to 45 kbp inside of the region of tight linkage (Fig. 1b, Table 1). None of the four inversion regions included centromeres[42]; thus, all of them seemed to contain paracentric inversions, which, in contrast to pericentric inversions, are not expected to decrease the fitness of heterozygotes[54].

To determine which of the two genomes carries the derived arrangement in each case, we also aligned contigs from the long-read-based genome assembly of *Melanogrammus aeglefinus* (melAeg)[56], a closely related outgroup within the subfamily Gadinae, to the gadMor2 assembly. We again identified split contig alignments mapping near the boundaries of the supergenes on LGs 1 and 7, indicating that for these supergenes, it is the gadMor2 genome that carries the derived arrangement (Fig. 1b, Table 1, Supplementary Table 3). In contrast, a single contig of the melAeg assembly was clearly colinear to the gadMor2 assembly in a region that extended about 150 kbp in both directions from one of the ends of the supergene on LG 2, indicating that the derived arrangement on LG 2 is carried not by the gadMor2 genome but by the gadMor Stat genome instead. For the supergene on LG 12, on the other hand, no informative alignments between the melAeg assembly and the gadMor2 assembly were found; thus, our contig-mapping approach did not allow us to determine which of the two *Gadus morhua* genomes carries the derived arrangement on this linkage group (however, our subsequent demographic analyses suggested that it is the gadMor Stat genome that carries the derived arrangement on LG 12; see below). Repeat content and mutation load were not increased in supergene regions compared to the genome-wide background (Supplementary Figure 1).

### Rapid divergence and introgression among codfishes

To establish the phylogenetic context within which the inversions arose in *Gadus morhua* and to test whether introgression played a role in the origin of the four supergenes[4, 57], we performed Bayesian analysis with the isolation-with-migration model as implemented in the AIM add-on package for BEAST 2[58, 59]. The dataset used in this analysis consisted of 109 alignments of sequences for the four species of the genus *Gadus* (*G. morhua, G. chalcogrammus, G. ogac*, and *G. macrocephalus*), their most closely related relatives *Boreogadus saida* and *Arctogadus glacialis*, and three outgroups (Fig. 2a; Supplementary Table 5). The 109 alignments (total length: 383,727 bp) were sampled from across the genome, excluding the four supergene regions and selecting the regions with the most reliable orthology (i.e., those unambiguouly aligned in the three-way whole-genome alignment) and the weakest signals of recombination (i.e., those with low numbers of hemiplasies). We found strong support (Bayesian posterior probability, BPP: 0.997) for a topology in which *Arctogadus* is either placed as the sister taxon to *Gadus* (BPP: 0.763) or to *Boreogadus* (BPP: 0.234). Introgression was supported (by Bayes factors greater than 10) between *Boreogadus* and *Arctogadus* in both cases and additionally from the common ancestor of the genus *Gadus* to *Arctogadus* in the latter case (Fig. 2b, Supplementary Figure 3). Regardless of the position of *Arctogadus*, our analysis with the isolation-with-migration model supported an age of around 4 Ma for the clade comprising the three genera (mean estimates: 3.81 Ma and 3.99 Ma; 95% HPD: 4.44–3.19 Ma and 4.56–3.33 Ma; Fig. 2b).

**Fig. 2.**
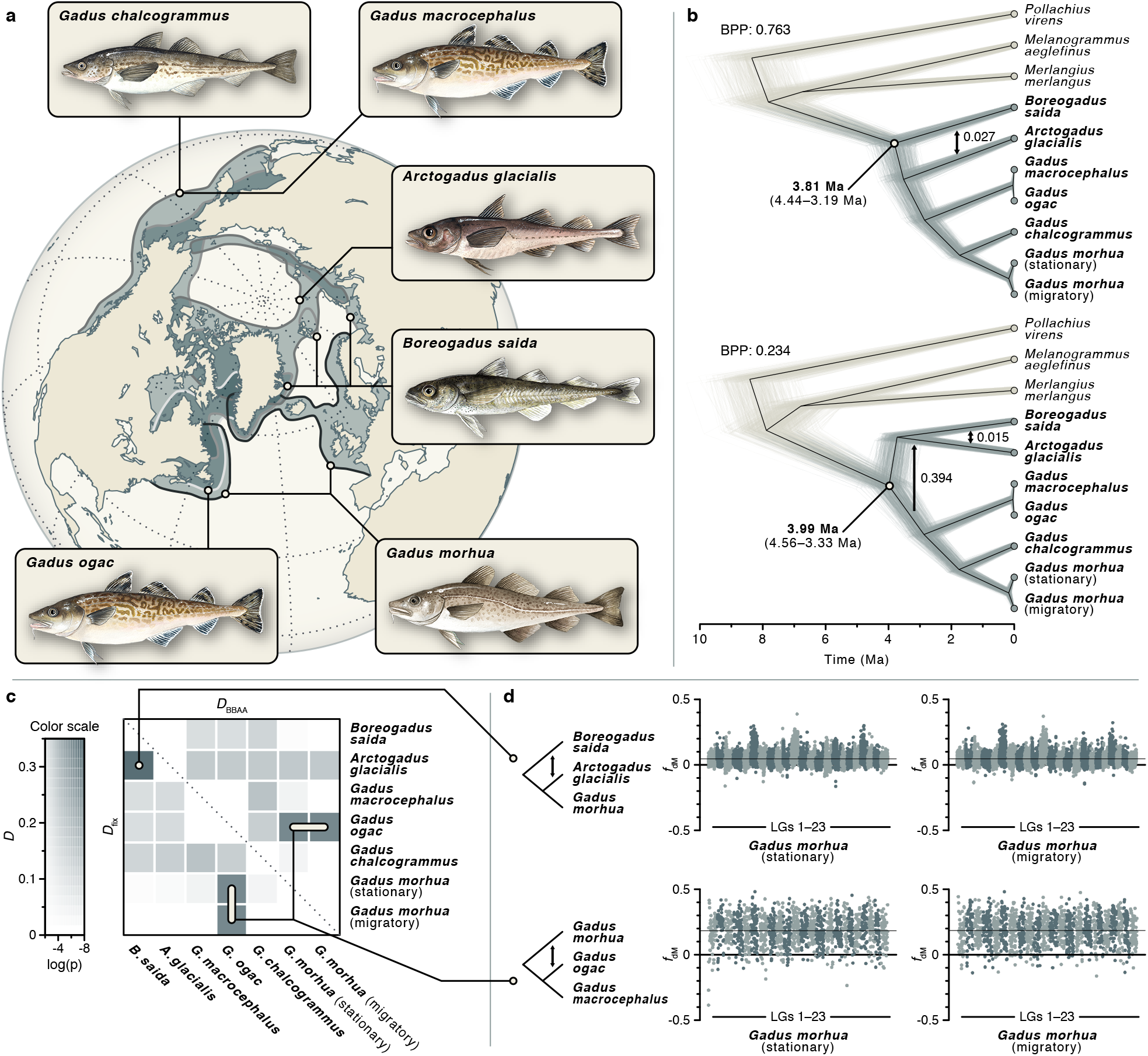
Divergence times and introgression among Gadinae. **a** Distribution ranges of sampled species of the genera *Gadus, Arctogadus*, and *Boreogadus*. Partially overlapping distribution ranges are shown in dark grey, with outline shades indicating the species (all distributions are shown separately in Supplementary Figure 4). **b** Species tree of the six species and three outgroups (*P. virens, M. aeglefinus*, and *M. merlangus*; outgroups shown in beige), estimated under the isolation-with-migration model from 109 alignments with a total length 383,727 bp. The Bayesian analysis assigned 99.7% of the posterior probability to two tree topologies that differ in the position of *Arctogadus glacialis* and were supported with Bayesian posterior probabilities (BPP) of 0.763 and 0.234, respectively. Rates of introgression estimated in the Bayesian analysis are marked with arrows. Thin grey and beige lines show individual trees sampled from the posterior distribution; the black line indicates the maximum-clade-credibility summary tree, separately calculated for each of the two topologies. Of *G. morhua*, both migratory and stationary individuals were included. **c** Pairwise introgression among species of the genera *Gadus, Arctogadus*, and *Boreogadus*. Introgression was quantified with the *D*-statistic. The heatmap shows two versions of the *D*-statistic, *D*_fix_ and *D*_BBAA_, below and above the diagonal, respectively. **d** Introgression across the genome. The *f*_dM_-statistic[60] is shown for sliding windows in comparisons of three species. The top and bottom rows show support for introgression between *B. saida* and *A. glacialis* and between *G. morhua* and *G. ogac*, respectively. Results are shown separately for the stationary and migratory *G. morhua* genomes. The mean *D*-statistic across the genome is marked with a thin solid line. Fish drawings by Alexandra Viertler.

Introgression among the three genera was further supported by Patterson’s *D*-statistic[61, 62]. Using Dsuite[63], we calculated for all possible species trios two versions of this statistic from a set of 19,035,318 SNPs: *D*_fix_, for which taxa in the trio are arranged according to a provided input tree, and *D*_BBAA_, for which taxa are arranged so that the number of sites with the “BBAA” pattern is maximised. The strongest signals of introgression were found once again between *Boreogadus* and *Arctogadus*, in a trio together with *Gadus morhua* (*D*_fix_ = 0.360, *p <* 10^−10^; Fig. 2c, Supplementary Tables 6 and 7). Within the genus *Gadus*, the *D*-statistic additionally provided strong support for introgression between the geographically co-occurring *G. ogac* and *G. morhua* in a trio with *G. macrocephalus* (*D*_fix_ = *D*_BBAA_ = 0.283, *p <* 10^−10^; Supplementary Tables 6 and 7). In both cases, signals of introgression quantified with the *f*_dM_-statistic[60] (an alternative to the *D*-statistic that is unbiased in small genomic regions[64] and symmetrically distributed around zero in the absence of introgression[63]) were found on all chromosomes, and were not affected by the choice of genome representing *Gadus morhua* (Fig. 2d). The introgression signal was not elevated in supergene regions, suggesting that they did not arise in *G. morhua* due to introgression from *G. ogac*. The occurrence of introgression between *Boreogadus* and *Arctogadus* was corroborated by a tree-based equivalent to Patterson’s *D*-statistic that does not rely on the molecular-clock assumption (the *D*_tree_-statistic of Ronco et al.[65]) and by genealogy interrogation[66, 67], but introgression between *G. ogac* and *G. morhua* did not receive this additional support (Supplementary Figure 5).

### Recent divergence among *Gadus morhua* populations

We performed phylogenomic analyses for individuals from eight *Gadus morhua* populations covering the species’ distribution in the North Atlantic (Fig. 3a; Supplementary Table 8). In addition to the individuals used for the gad-Mor2 and gadMor Stat assemblies, we selected from these populations 22 individuals for which preliminary analyses had shown that each of them carried, at each of the four supergene regions, two copies of the same haplotypes (i.e., they were homokaryotypic). For the four sampling localities Newfoundland, Iceland, Lofoten, and Møre, we discriminated between “migratory” and “stationary” individuals based on whether they carried the same supergene haplotype on LG 1 as the gadMor2 genome or the same as the gadMor Stat genome. At other localities, all individuals were considered stationary based on the well-known migration patterns of *Gadus morhua*[68]. For the individuals from Lofoten and Møre, this classification could be confirmed by an analysis of their otoliths[55], but otolith data was not available for the individuals from the other sampling localities. Based on a dataset of 20,402,423 genome-wide biallelic SNPs, we estimated relationships and divergence times among *Gadus morhua* populations under the multi-species coalescent model with SNAPP[69, 70], first only with data from outside of the supergene regions. In line with previous studies based on SNP arrays[27, 35, 71], we found the primary divergence within *Gadus morhua* to separate the populations of the Northwest Atlantic from those of the Northeast Atlantic (including Iceland). We estimated these groups to have diverged around 65.4 ka (95% HPD: 84.3–46.2 ka) but acknowledge that these results may underestimate the true divergence time because the applied model does not account for possible gene flow after divergence (Fig. 3b, Supplementary Figure 6).

**Fig. 3.**
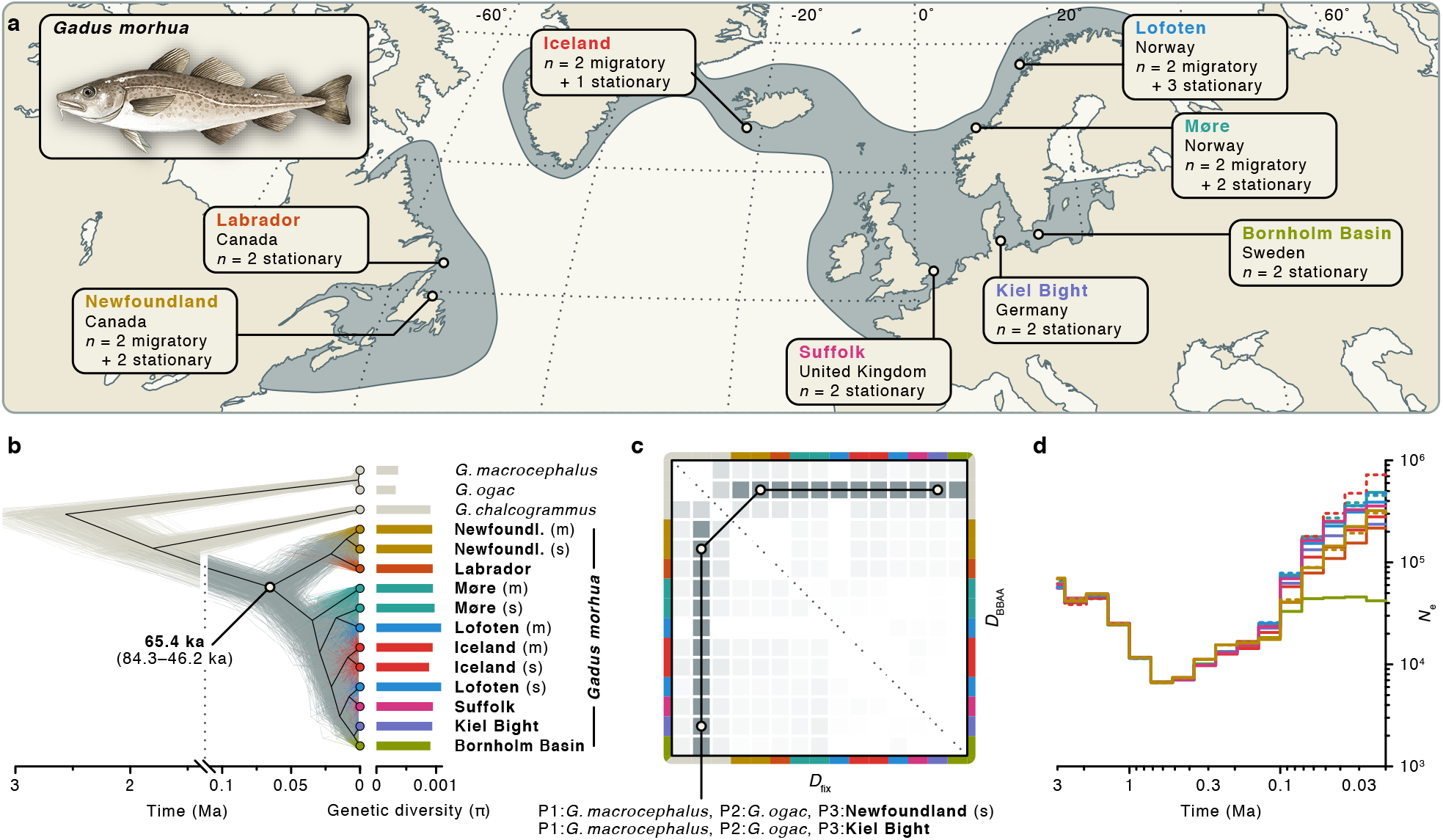
Divergence times, demography, and introgression among *Gadus morhua* populations. **a** Geographic distribution of *Gadus morhua* in the North Atlantic and sampling locations for analyses of population divergence times, demography, and introgression. **b**, Tree of *Gadus morhua* populations and three outgroups (in beige; *G. macrocephalus, G. ogac*, and *G. chalcogrammus*), inferred under the multi-species coalescent model from 1,000 SNPs sampled across the genome (excluding inversion regions). Thin grey and beige lines show individual trees sampled from the posterior distribution; the black line indicates the maximum-clade-credibility summary tree. Estimates of the genetic diversity (*π*) per population are indicated by bars to the right of the tips of the tree. **c** Pairwise introgression among *Gadus morhua* populations and outgroup species. Two versions of the *D*-statistic, *D*_fix_ and *D*_BBAA_, are shown above and below the diagonal, respectively. Colour codes on the axes indicate populations, and heatmap colouring is as in Fig. 2c. The two trios with the strongest signals, supporting introgression between *G. ogac* and both the Kiel Bight and the stationary Newfoundland *G. morhua* population with *D*_BBAA_ = *D*_fix_ = 0.250 are marked. **d** Population sizes (*N*_e_) over time in *Gadus morhua* populations, estimated with Relate. For the Newfoundland, Møre, Iceland, and Lofoten populations, migratory (m) and stationary (s) individuals were analysed separately; dashed lines are used for migratory populations.

The genetic diversity, quantified by *π*[72], was comparable among the populations of both groups, ranging from 8.82 × 10^−4^ to 1.084 × 10^−3^ (Supplementary Table 9). Applied to the set of 20,402,423 SNPs, Patterson’s *D*-statistic corroborated the occurrence of introgression between *Gadus ogac* and *Gadus morhua* and showed that this signal of introgression is similar in all populations (*D*_fix_ = *D*_BBAA_ = 0.249, *p <* 10^−10^; Fig. 3c, Supplementary Tables 10 and 11). Estimating changes in the population size (*N*_e_) over time for *Gadus morhua* with Relate[73] revealed a Pleistocene bottleneck, lasting from around 0.7 Ma to 0.3 Ma, during which the population size of the common ancestor of all populations decreased from around 50,000 to 7,000 diploid individuals. The subsequent increase in population sizes occurred in parallel with the diversification of *Gadus morhua* populations and was experienced by all of them to a similar degree but less so by the Bornholm Basin population of the Baltic Sea (Fig. 3d).

### Different age estimates for supergenes in *Gadus morhua*

To infer the ages of the supergenes on LGs 1, 2, 7, and 12, we applied SNAPP analyses to SNPs from each supergene separately, extracted from the dataset of 20,402,423 biallelic SNPs. For each of the four supergenes, we recovered a deep divergence separating the haplotypes with ancestral and derived arrangements; however, the age estimates for this divergence differed widely among the four supergenes, with mean age estimates of 0.61 Ma (95% HPD: 0.77–0.46 Ma) for the supergene on LG 1 (Fig. 4a), 0.88 Ma (95% HPD: 1.10–0.67 Ma) for the supergene on LG 2 (Fig. 4d), 1.66 Ma (95% HPD: 2.05–1.29 Ma) for the supergene on LG 7 (Fig. 4g), and 0.40 Ma (95% HPD: 0.50–0.30 Ma) for the supergene on LG 12 (Fig. 4j; Supplementary Figure 7). The migratory individuals from Newfoundland, Iceland, Lofoten, and Møre shared the same arrangement on all four linkage groups, and so did the stationary individuals from Lofoten, Suffolk, and Kiel Bight. The genetic diversity of the haplotype with the derived arrangement was on average lower on LGs 1 and 12, but higher on LGs 2 and 7 (Figs. 4a,d,g,j, Supplementary Table 9).

**Fig. 4.**
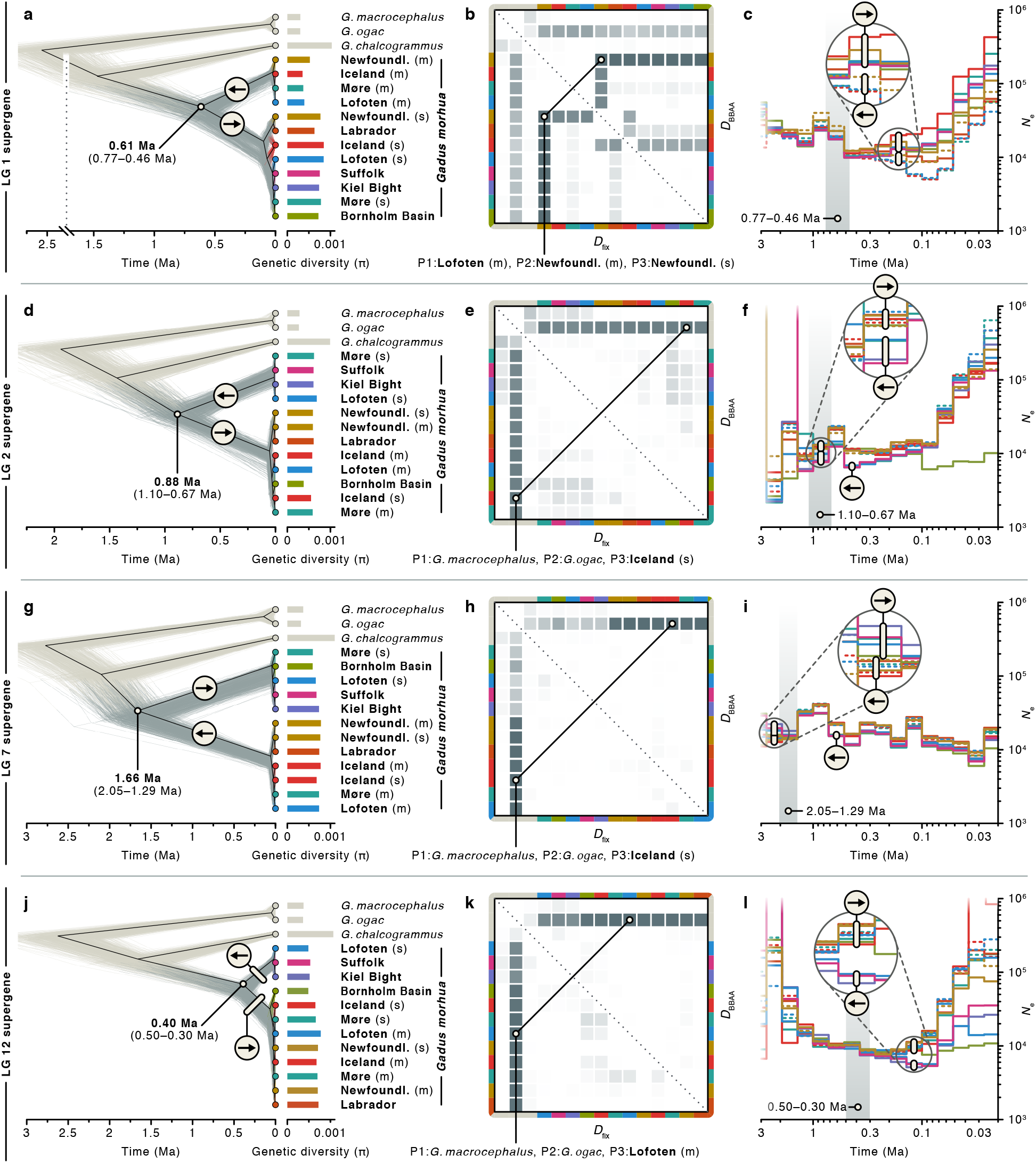
Divergence times and demography of *Gadus morhua* populations and gene flux within supergene regions. **a**,**d**,**g**,**j** Trees of *Gadus morhua* populations and three outgroups (in beige; *G. macrocephalus, G. ogac*, and *G. chalcogrammus*), inferred under the multi-species coalescent model from 1,000 SNPs sampled from within the supergene regions on LGs 1 (**a**), 2 (**d**), 7 (**g**), and 12 (**j**). Thin grey and beige lines show individual trees sampled from the posterior distribution; the black line indicates the maximum-clade-credibility summary tree. Within *G. morhua*, derived and ancestral arrangements are marked with forward and reverse arrows, respectively. Estimates of the genetic diversity (*π*) per population within supergene regions are indicated by bars to the right of the tips of the tree. **b**,**e**,**h**,**k** Pairwise signals of past gene flow among *Gadus morhua* populations and outgroup species within the supergene regions on LGs 1 (**b**), 2 (**e**), 7 (**h**), and 12 (**k**). Two versions of the *D*-statistic, *D*_fix_ and *D*_BBAA_, are shown above and below the diagonal, respectively. Colour codes on the axes indicate populations, ordered as in **a**,**d**,**g**,**j**. The trios with the strongest signals of gene flow are labelled. **c**,**f**,**i**,**l** Population sizes (*N*e) over time in *Gadus morhua* populations for the supergene regions on LGs 1 (**c**), 2 (**f**), 7 (**i**), and 12 (**l**). For the Newfoundland, Møre, Iceland, and Lofoten populations, migratory (m) and stationary (s) individuals were analysed separately; dashed lines are used for migratory populations. The grey regions indicate the confidence intervals for the inferred age of the split between the two haplotypes (from **a**,**d**,**g**,**j**).

### Gene flux between haplotypes via gene conversion

Just like gene flow between populations can lead to underestimates of their divergence time, the ages of the separation between the two alternative arrangements per supergene could be underestimated if gene flux (defined as exchange of alleles during meiosis) has occurred after their divergence, through gene conversion or double crossover[74]. To assess the frequency of these two processes within the four supergene regions, we calculated the *D*-statistic for the sets of supergene-specific SNPs. The *D*-statistic supported gene flux (and thus the occurrence of either gene conversion or double crossover) between haplotypes with derived and ancestral arrangements particularly for the supergene on LG 1 and the geographically co-occurring migratory or stationary Newfoundland populations (*D*_fix_ = *D*_BBAA_ = 0.383, *p <* 10^−10^; Fig. 4b, Supplementary Tables 12 and 13). To test whether gene conversion could be the cause of this gene flux occurring between the Newfoundland populations, we tested for the GC bias expected from gene conversion[24, 25], comparing GC-content of sites shared between the two Newfoundland populations (“ABBA” sites) to that of sites shared between the migratory Newfoundland population and other migratory populations (“BBAA” sites). The mean GC-content of the former, 0.482, is indeed significantly higher than that of the latter, 0.472 (one-sided *t*-test; *t* = *−*4.74, df = 27833, *p <* 10^−5^), supporting gene conversion as an agent of gene flux between haplotypes with derived and ancestral arrangements in the Newfoundland populations. For the supergenes on LGs 2, 7, and 12, the *D*-statistic indicated only comparatively weak signals of gene flux between derived and ancestral arrangements (*D ≤* 0.2, *p ≥* 10^−4^; Fig. 4e,h,k, Supplementary Tables 14-19).

### Demographic analyses recover signatures of bottlenecks following inversions

It is likely that each of the inversions associated with the four supergenes in *Gadus morhua* originated just once and that all of the current carriers of the inversion descended from the individual in which it originated. The supergene origin should therefore have been equivalent to an extreme bottleneck event during which the population size was reduced to a single sequence, but which affected only the inversion region, and only the carriers of the derived arrangement. We might thus expect to see signatures of this extreme bottleneck event in analyses of population size over time, in the form of differences in ancestral population sizes between the haplotypes with derived and ancestral arrangements that date to the time of supergene origin.

To verify that signatures of extreme bottlenecks are detectable in descending genomes even after long periods of time, we first performed a series of power analyses based on coalescent simulations (Supplementary Note 2). After confirming that the program Relate is in principle able to pick up such signals, we tested for the presence of bottleneck signatures associated with inversion events by performing demographic analyses with Relate separately for sets of SNPs from each of the four supergene regions. As expected due to the smaller amount of input data (1–3% compared to the genome-wide SNP data), these analyses produced estimates that were less clear (Fig. 4c,f,i,l) than those obtained with genome-wide SNP data (Fig. 3d). Nevertheless, the supergene-specific demographic analyses supported a difference between the population sizes of haplotypes with derived and ancestral arrangements, with a temporary reduction of population sizes for the haplotype with the derived arrangement that coincided with the estimated ages of the supergenes on LGs 2 and 7 (Fig. 4f,i). For the supergenes on LGs 1 and 12, the difference in the population sizes did not coincide with the supergene ages. However, for the supergene on LG 1, the population size of the haplotype with the derived arrangement was reduced compared to the ancestral arrangement in several time intervals following the supergene origin (Fig. 4c). While our contig-mapping approach had not allowed us to infer which of the two haplotypes of the LG 12 supergene had the derived arrangement, the reduced inferred population size of the haplotype carried by the sampled individuals from the Suffolk, Kiel Bight, and stationary Lofoten populations, in a time interval following the supergene origin, suggested that this haplotype might have the derived arrangement on that linkage group. The ancestral population sizes estimated from the supergene regions on LGs 2 and 7 further highlighted that estimates of the current genetic diversity may not always be useful for the identification of derived arrangements (Fig. 4i), as the population-size estimates for the haplotype with the ancestral arrangement were in both cases higher than for the derived arrangement at the time of supergene origin, but lower for most of the subsequent time towards the present (Fig. 4i).

### Divergence-time profiles reveal double crossovers

To explore whether divergence times between the two arrangements per supergene are homogeneous across the supergene region, we repeated divergence-time inference with SNAPP in sliding windows of 250 kbp along all linkage groups. We expected that if any gene flux between haplotypes with derived and ancestral arrangements should proceed via double crossovers, its effect should be less pronounced near the inversion breakpoints at the boundaries of the supergenes and stronger towards their centers, which could generate U-shaped divergence profiles for supergene regions[20, 74, 75]. Contrary to this expectation, the divergence-time profiles were relatively homogeneous from beginning to end, particularly for the supergenes on LGs 1, 7, and 12 (Fig. 5a,c,d; Supplementary Figure 8), suggesting either that double crossovers are rare within these supergenes, or that sequences exchanged through double crossovers are frequently purged from the recipient haplotypes. As the supergene on LG 1 is known to include not one but two adjacent inversions of roughly similar size[32], our results also suggested a similar age and possibly a joint origin for both of these inversions. The divergence-time profile for the supergene on LG 2 appeared consistent with the expectation of a U-shaped pattern; however, comparison of the gadMor2 assembly with the recently released gadMor3 assembly[42] showed that the end of this linkage group may be misassembled in gadMor2, and the region of low divergence time appearing within the supergene (around position 22.5–23.0 Mbp) may in fact lie outside of it (Supplementary Figure 2). Additional analyses of *F*_st_ and *d*_xy_ in windows across LGs 1, 2, 7, and 12, performed with both the gadMor2 and gadMor3 assemblies as references, confirmed this assumption as well as the absence of U-shaped patterns for the four regions (Supplementary Figure 9). However, a closer look at the divergence-time profile for LG 12 revealed a single window within the supergene in which the otherwise clear separation between the groups carrying the alternative arrangements was interrupted: Unlike in all other windows within this supergene, the Bornholm Basin population grouped (BPP: 1.0) with the three populations representing the derived arrangement (Suffolk, Kiel Bight, and stationary Lofoten; see Fig. 4j) in the window for positions 7.50–7.75 Mbp (Fig. 5d). To investigate the genotypes of the two sampled Bornholm Basin individuals within this region in more detail, we identified 219 haplotype-informative sites between positions 7–8 Mbp on LG 12, and found that these individuals were both heterozygous at these sites, for a region of ∼275 kbp between positions 7,478,537 bp and 7,752,994 bp (Fig. 6). The two individuals from the Bornholm Basin population thus carried a long sequence from the haplotype with the derived arrangement even though they were otherwise clearly associated with the ancestral arrangement. As the length of this introduced sequence was far longer than the 50–1,000 bp expected to be copied per gene-conversion event[23, 24], it strongly supports double crossover between the two haplotypes of the LG 12 supergene. The region covered by the introduced sequence contains 24 predicted genes (Supplementary Table 20), including a cluster of three vitellogenin genes, out of a total of four vitellogenin genes found in the gadMor2 genome. These genes are known to influence the buoyancy of fish eggs[76–79], and could thus be a target of selection in *Gadus morhua* populations in the brackish Baltic Sea[28].

**Fig. 5.**
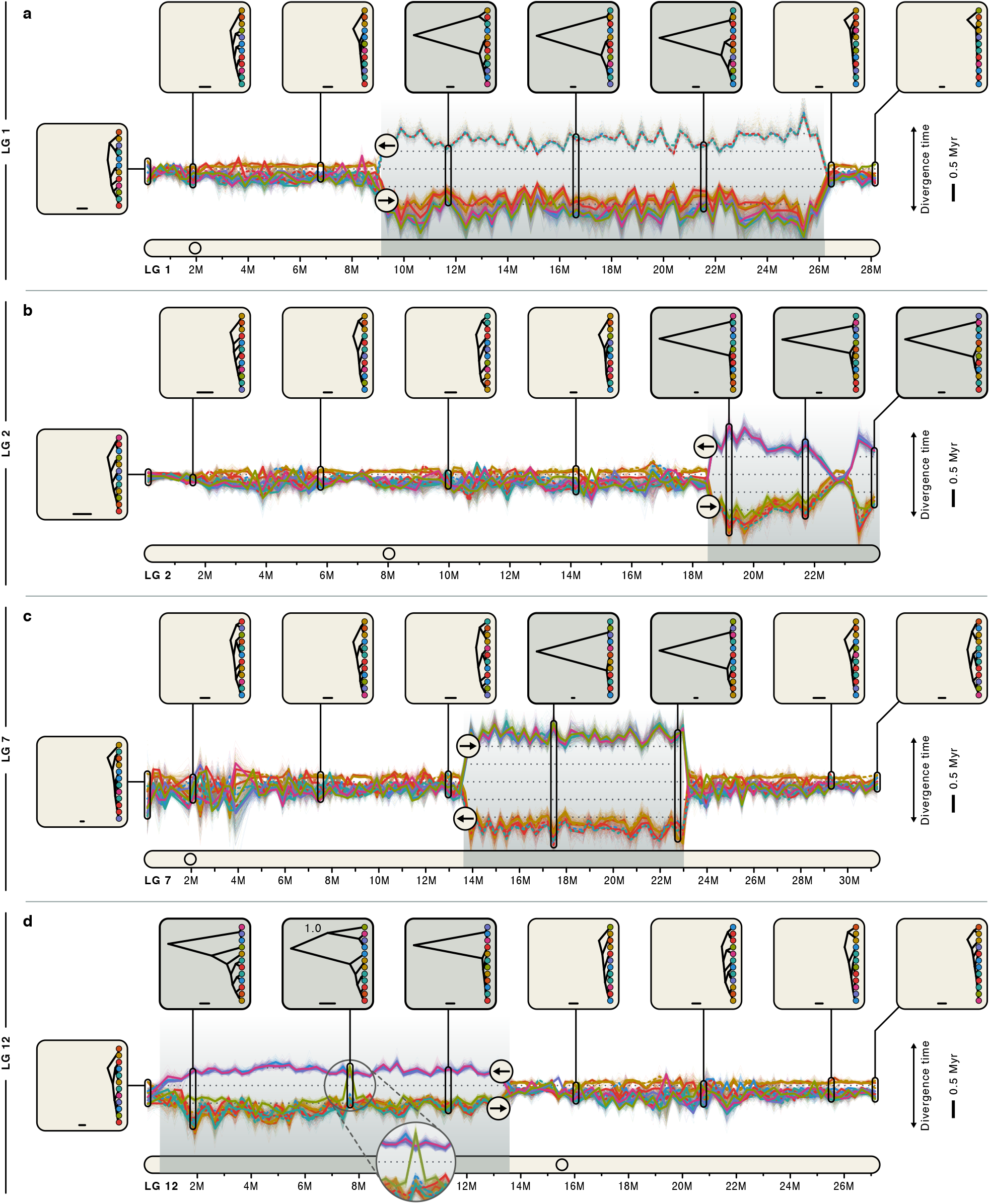
Divergence-time profiles for linkage groups with supergenes. **a**–**d** Illustration of between-population divergence times along LGs 1 (**a**), 2 (**b**), 7 (**c**), and 12 (**d**), estimated with SNAPP from SNPs in sliding windows. Supergene regions are indicated by grey backgrounds. Along the vertical axis, the distance between two adjacent lines shows the time by which the corresponding populations have been separated on the ladderised population tree for a given window; the scale bar indicates a duration of 0.5 Myr. Examples of the population tree are shown in insets for eight selected windows. The scale bar in these insets indicates the branch length equivalent to 50,000 years. The node label in one inset in (**d**) indicates the support for the grouping of the Bornholm Basin population with three populations representing the derived arrangement (BPP: 1.0).

**Fig. 6.**
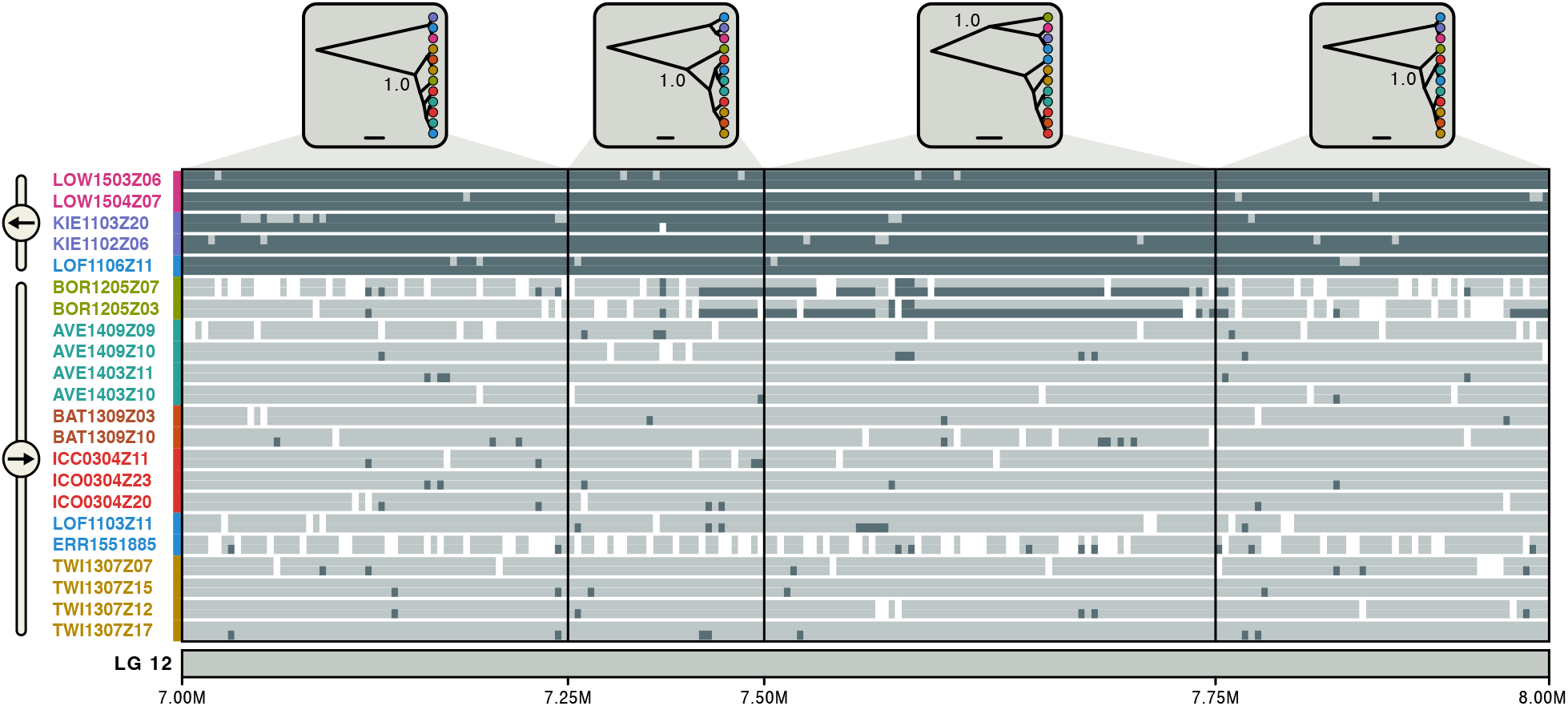
Ancestry painting for part of the supergene on LG 12. The ancestry painting[67, 80] shows genotypes at 219 haplotype-informative sites between positions 7 and 8 Mbp on LG 12, within the supergene on that linkage group. For each of 22 *Gadus morhua* individuals, homozygous genotypes are shown in dark or light grey while heterozygous genotypes are illustrated with a light grey top half and a dark grey bottom half; white colour indicates missing genotypes. As haplotype-informative sites, we selected those that have less than 10% missing data and strongly constrasting allele frequencies (≥ 0.9 in one group and ≤ 0.1 in the other) between the group carrying the derived arrangement (individuals from Suffolk, Kiel Bight, and stationary Lofoten) and the group carrying the ancestral arrangement (individuals from the Møre, Labrador, Reykjavík, migratory Lofoten, and Newfoundland populations). The four insets at the top show population trees inferred with SNAPP; the node labels in these insets indicate Bayesian support for the grouping of the Bornholm Basin population with either the derived or ancestral arrangement.

## Discussion

Through comparison of long-read-based genome assemblies for migratory and stationary Atlantic cod individuals, we corroborated earlier conclusions that chromosomal inversions underlie all of the four supergenes in Atlantic cod[42]. The inversion breakpoints do not coincide exactly with the boundaries of supergenes, but lie up to 45 kbp inside of them, in agreement with findings reported for *Drosophila* that suggested that recombination suppression can extend beyond inversion breakpoints [53]. By also comparing the genome assemblies for Atlantic cod with an assembly of the closely related haddock (*Melanogrammus aeglefinus*)[56], we were further able to identify the gadMor2 assembly — representing a migratory Atlantic cod individual — as the carrier of the derived arrangement of the supergenes on LG 1 and 7, but not of that on LG 2. In addition, our demographic analyses indicated that the gadMor2 assembly might also carry the ancestral arrangement on LG 12. The haplotypes with derived arrangements were not consistently characterised by lower genetic diversity than ancestral arrangements, contrary to assumptions that were used in previous studies to distinguish between them[29, 30]. As suggested by our demographic analyses (Fig. 4i) and simulations (Supplementary Table 21), this contrast can be explained by an ability of haplotypes with derived arrangements to recover from the initial bottleneck. While this recovery may require substantial frequencies of the derived arrangement (comparable to those of the ancestral one) within the species and sufficient time since the inversion origin to allow the accumulation of new mutations[17], both of these requirements may be met for the Atlantic cod supergenes. Additionally, haplotypes with the derived arrangement can have higher genetic diversity than those with the ancestral arrangement if the latter were subject to a selective sweep after the divergence.

According to our time-calibrated phylogenomic analyses, the four supergenes in Atlantic cod originated between ∼0.40 and ∼1.66 Ma. These dates could potentially be underestimates due to gene flux between the haplotypes with derived and ancestral arrangement that could not be accounted for in our age estimation. Nevertheless, we consider these age estimates as evidence that at least some of the four supergenes had separate origins, as the age estimate for the supergene on LG 7 is more than four times as old as that for the supergene on LG 12 and their confidence intervals are clearly non-overlapping. This conclusion is further supported by the homogeneity in sliding-window age estimates from beginning to end of each supergene and the support for demographic bottlenecks coinciding with the inferred age estimates (Figs. 4,5). On the other hand, a joint origin can not be excluded for the supergenes on linkage groups 1, 2, and 12. Our age estimates thus indicate that some, but not necessarily all of the four supergenes evolved separately, after the Atlantic cod’s divergence from the walleye pollock (*Gadus chalcogrammus*), and before the divergence of all extant Atlantic cod populations.

Introgression from other codfish species does not seem to have played a role in the origin of the four supergenes. However, we found strong signals of introgression between Atlantic cod and Greenland cod (*Gadus ogac*), as Atlantic cod unambigously shares more alleles with Greenland cod than with the Greenland cod’s sister species, the Pacific cod (*Gadus macrocephalus*), despite the very recent divergence between Greenland and Pacific cod (∼40 ka). This genetic similarity between Atlantic cod and Greenland cod does not appear to be an artifact resulting from reference bias and is best explained by gene flow into Greenland cod, from an Atlantic cod population cooccurring in the Northwest Atlantic (Supplementary Note 3). Besides the signals for introgression into Greenland cod, our results strongly support gene flux between the two haplotypes of the supergene on LG 1 (Fig. 4b): between all carriers of the ancestral arrangement and the migratory individuals from Newfoundland (which carry the derived arrangement) and between all carriers of the derived arrangement and the stationary individuals from Newfoundland (which carry the ancestral arrangement). Due to elevated GC-content of sites shared between the haplotypes, we interpret these signals of gene flux as evidence for gene conversion occurring at Newfoundland, but note that double crossovers could potentially also influence GC-content through crossover-associated gene conversion[81]. It remains unclear why gene flux through gene conversion does not seem to occur at other locations where the two arrangements co-occur (e.g. in northern Norway).

As a second mechanism allowing gene flux between haplotypes with derived and ancestral arrangements, our results demonstrated the occurrence of double crossover, between the two haplotypes of the LG 12 supergene. The sequence introduced through double crossover included the vitellogenin gene cluster[82], which is assumed to contribute to the proper hydration of fish eggs and thus to the maintenance of neutral buoyancy[76–79]. However, in contrast to the fully marine environments of the open North Atlantic, the almost land-locked Baltic Sea has a severely reduced salinity, and thus requires adaptations in the hydration of eggs, so that they remain neutrally buoyant at a salinity of ∼12.5 ppt[83] (compared to ∼35 ppt in the North Atlantic and ∼18 ppt at Kiel Bight[84]) and do not sink to anoxic layers[85–87]. Thus, the three vitellogenin genes may be under selection in *Gadus morhua* from the Baltic Sea[28], increasing the frequency of the introduced sequence within the Bornholm Basin population.

The presence of four long (4–17 Mbp) and old (0.40–1.66 Ma) inversion-based supergenes in Atlantic cod adds to recent findings of inversions of similar size and/or age in butterflies[12], ants[2], birds[4], lampreys[88], and *Drosophila*[75]. For non-model organisms, these findings are largely owed to improvements in sequencing technology within the last decade, including long-read sequencing and chromosome conformation capture techniques, and may become more common as these techniques are applied to an increasing number of species. These findings are, however, in contrast to earlier expectations based on theoretical work and empirical studies on selected model organisms: Only about 20 years ago, the available evidence indicated that inversions (and thus inversion-based supergenes) would be “generally not ancient”[20], with maximum ages on the order of *N*_e_ generations (which would correspond to ∼100 ka in Atlantic cod), because they would either degenerate through the accumulation of mutation load, or erode if gene flux occurs through gene conversion and double crossover[20, 74, 89]. However, we observed neither mutation-load accumulation nor erosion of supergenes in Atlantic cod: Mutation load was not increased within the four supergenes compared to the genome-wide background. And supergene erosion — at least when resulting from double crossovers — would be expected to produce U-shaped divergence profiles[20, 74, 75], but no such profiles were found for the Atlantic cod supergenes (Fig. 5). These observations mirror those made recently by Yan et al.[2] for a supergene in fire ants, and we thus concur with their conclusion that “low levels of recombination and/or gene conversion may play an underappreciated role in preventing rapid degeneration of supergenes”. But, since our results also indicated that selection may have acted on sequences exchanged between supergene haplotypes, we further suggest that — just like in interbreeding species that maintain stable species boundaries despite frequent hybridisation[67] — selective purging of introduced sequences may also be important for the maintenance of supergenes, by maintaining the rate of gene flux between haplotypes at exactly the right balance, between too little of it and the consequential mutation-load accumulation, and too much of it and the resulting supergene erosion.

## Methods

### Construction of the gadMor Stat genome assembly

We performed high-coverage genome sequencing for a stationary *Gadus morhua* individual (LOF1106Z11) sampled at the Lofoten islands in northern Norway. The specimen used for this was selected based on a preliminary investigation that had suggested that it carried, homozygously on each of the four LGs 1, 2, 7, and 12, a supergene haplotype that was complementary to the one in the gadMor2 genome[31], which represents a migratory individual from northern Norway. We used the Pacific Biosciences RS II platform, operated by the Norwegian Sequencing Centre (NSC; www.sequencing.uio.no), to generate 2.4 million PacBio SMRT reads with a total volume of 12.5 Gbp. This is approximately equivalent to an 19*×* coverage of the *Gadus morhua* genome, the size of which has been estimated at 650 Mbp[31]. The PacBio SMRT reads were assembled with Celera Assembler v.8.3rc2[90], adjusting the following settings according to the nature of the PacBio reads (all others were left at their defaults): merSize =16, merThreshold=0, merDistinct=0.9995, merTotal=0.995, ovlErrorRate=0.40, ovlMinLen=500, utgGraphErrorRate=0.300, utgGraphErrorLimit=32.5, utgMergeErrorRate=0.35, utgMergeErrorLimit=40, utgBubblePopping=1, utgErrorRate=0.40, utgErrorLimit=25, cgwErrorRate=0.40, cnsErrorRate=0.40. The consensus sequence of the assembly was polished with Quiver v.0.9.0[91] and refined with Illumina reads sequenced for the same individual (see below). A total volume of 6.2 Gbp of Illumina reads were mapped to the assembly with BWA MEM v.0.7.12-r1039[92] and sorted and indexed with SAMtools v.1.10[93, 94]. Subsequently, Pilon v.1.16[95] was applied to recall consensus and assembly completeness was assessed with BUSCO v.5.0[51], using the Actinopterygii dataset of conserved gene sequences.

### Whole-genome sequencing and population-level variant calling

Twenty-two migratory and stationary *Gadus morhua* individuals were sampled in Canada, Iceland, the United Kingdom, Germany, Sweden, and Norway using longline fishing, hand fishing, gillnets, and trawling, and subjected to medium-coverage (8.1–17.0*×*) whole-genome Illumina sequencing (Fig. 3; Supplementary Table 8). DNA extraction, library preparation, and sequencing were performed at the NSC using the Illumina Truseq DNA PCR-free kit for DNA extraction and an Illumina HiSeq 2500 instrument with V4 chemistry for paired-end (2 *×* 125 bp) sequencing. Reads from these 22 individuals were mapped to the gadMor2 assembly for *Gadus morhua*[31], together with Illumina reads from the two individuals used for the gadMor2 and gadMor Stat assemblies. Mapping was done with BWA MEM v.0.7.17, followed by sorting and indexing with SAMtools v.1.9. Read duplicates were marked and read groups were added with Picard tools v.2.18.27 (http://broadinstitute.github.io/picard). Variant calling was performed with GATK’s v.4.1.2.0[96, 97] HaplotypeCaller and GenotypeGVCFs tools, followed by indexing with BCFtools v.1.9[94].

### Delimiting high-LD regions associated with inversions

As chromosomal inversions locally suppress recombination between individuals carrying the inversion and those that do not, we used patterns of linkage disequilibrium (LD) to guide the delimitation of inversion regions for each of the four supergenes[28, 32, 42]. To maximize the signal of LD generated by the inversions, we selected 100 *Gadus morhua* individuals from a separate dataset (Supplementary Table 22) so that for each of the four supergenes, 50 individuals carried two copies of one of the two alternative supergene haplotypes, and the other 50 individuals carried two copies of the other. Variant calls of the 100 individuals were filtered with BCFtools, excluding all indels and multi-nucleotide polymorphisms and setting all genotypes with a Phred-scaled quality below 20, a read depth below 3, or a read depth above 80 to missing. Sites with more than 80% missing data or a minor allele count below 20 were then removed from the dataset with VCFtools v.0.1.14[98]. Linkage among single-nucleotide polymorphisms (SNPs) spaced less than 250,000 bp from each other was calculated with PLINK v.1.90b3b[99]. The strength of short- to mid-range linkage for each SNP was then quantified as the sum of the distances (in bp) between that SNP and all other SNPs with which it was found to be linked with *R*^2^ *>* 0.8. We found this measure to illustrate well the sharp decline of linkage at the boundaries of the four supergenes (Fig. 1).

### Contig mapping

To confirm the presence of chromosomal inversions within the four supergenes on LGs 1, 2, 7, and 12 of the *Gadus morhua* genome, we aligned contigs of the gadMor_Stat assembly to the gadMor2 assembly by using BLASTN v.2.2.29[100] searches with an e-value threshold of 10^−10^, a match reward of 1, and mismatch, gap opening, and gap extension penalties of 2, 2, and 1, respectively. Matches were plotted and visually analysed for contigs of the gadMor_Stat assembly that either span the boundaries of the four supergene regions or map partially close to both boundaries of one such region. We considered the latter to support the presence of a chromosomal inversion if one of two parts of a contig mapped just inside of one boundary and the other part mapped just outside of the other boundary, and if the two parts had opposite orientation; in contrast, an observation of contigs clearly spanning one of the boundaries would reject the assumption of an inversion. To further assess which of the two *Gadus morhua* genomes (gadMor2 or gadMor Stat) carries the haplotype with the derived arrangement at each of the four regions, we also aligned contigs of the genome assembly for *Melanogrammus aeglefinus* (melAeg)[56] to the gadMor2 assembly.

### Threeway whole-genome alignment

To identify the regions that are most reliably orthologous among the gadMor2, gadMor Stat, and melAeg assemblies, we generated whole-genome alignments using three different approaches. First, we visually inspected the plots of BLASTN matches (see above), determined the order and orientation of all gadMor Stat and melAeg contigs unambiguously mapping to the gadMor2 assembly, and then combined these contigs into a single FASTA file per species and gadMor2 linkage group. For each linkage group, pairwise alignments were then produced with the program MASA-CUDAlign v.3.9.1.1024[101]. Second, we used the program LASTZ v.1.0.4[102] to align both the gadMor_Stat assembly and the melAeg assembly to the gadMor2 assembly, after masking repetitive regions in all three assemblies with RepeatMasker v.1.0.8 (http://www.repeatmasker.org). Third, Illumina sequencing reads of the individuals used for the three assemblies were mapped to the gadMor2 assembly with BWA MEM, followed by sorting and indexing with SAMtools and conversion of the result files to FASTA format. Finally, we generated a conservative threeway whole-genome alignment by comparing the three different types of alignments and setting all sites to missing at which one or more of the three alignment types differed. Alignment sites that opened gaps in the gadMor2 sequence were deleted so that the resulting strict consensus alignment retained the coordinate system of the gadMor2 assembly.

Based on the threeway whole-genome alignment, we calculated the sequence divergence between the gadMor Stat and gadMor2 assemblies, relative to the sequence divergence between the melAeg and gadMor2 assemblies, in sliding windows of 100,000 bp. Sequence divergence was calculated in pairwise sequence comparisons as uncorrected p-distances. We also used the threeway wholegenome alignment to generate a mask of unreliable alignment sites, including all sites that had been set to missing in the alignment.

### Estimating divergence times of Gadinae

To estimate the ages of supergene origins in *Gadus morhua* based on the carefully calibrated timeline of Musilova et al.[103], we performed two nested phylogenetic analyses. The first one used constraints specified according to the results of Musilova et al.[103] to estimate the divergence times of species within the subfamily Gadinae. The second analysis was constrained according to the results of the first one to refine the divergence-time estimates among species of the genera *Gadus, Arctogadus*, and *Boreogadus* with a larger dataset and while co-estimating introgression (see below).

The phylogenomic dataset used for the first phylogenetic analysis comprised genome assemblies for eight Gadinae species published by Malmstrøm et al.[104], a genome assembly for the most closely related outgroup *Brosme brosme*[104], the gadMor2 assembly for *Gadus morhua*, and sets of unassembled Illumina reads for *Gadus macrocephalus* and *Gadus ogac*[50] (Supplementary Table 4). Aiming to identify sequences orthologous to 3,061 exon markers used in a recent phylogenomic analysis of teleost relationships by Roth et al.[105], we first performed targeted assembly of these markers from the sets of Illumina reads for *Gadus macrocephalus* and *Gadus ogac*. Targeted assembly was conducted with Kollector v.1.0.1[106], using marker sequences of *Gadus morhua* from Roth et al.[105] as queries. From the set of whole-genome and targeted assemblies, candidate orthologs to the 3,061 exon markers used by Roth et al.[105] were then identified through TBLASTN searches, using sequences of *Danio rerio* as queries as in the earlier study. The identified sequences were aligned with MAFFT and filtered to exclude potentially remaining paralogous sequences and misaligned regions: We removed all sequences with TBLASTN bitscore values below 0.9*×* the highest bitscore value and all sequences that had dN/dS values greater than 0.3 in comparison to the *Danio rerio* queries, we removed codons from the alignment for which BMGE v.1.1[107] determined a gap rate greater than 0.2 or an entropy-like score greater than 0.5, and we excluded exon alignments with a length shorter than 150 bp, more than two missing sequences, or a GC-content standard deviation greater than 0.04. We then grouped exon alignments by gene and excluded all genes that 1) were represented by less than three exons, 2) had one or more completely missing sequences, 3) were supported by a mean RAxML v.8.2.4[108] bootstrap value lower than 0.65, 4) were located within the four supergene regions, 5) exhibited significant exon tree discordance according to an analysis with Concaterpillar v.1.7.2[109], or 6) had a gene tree with non-clock-like evolution (mean estimate for coefficient of variation greater than 0.5 or 95% highest-posterior-density interval including 1.0) according to a relaxed-clock analysis with BEAST 2[59]. Finally, concatenated exon alignments per gene were inspected by eye, and six genes were removed due to remaining possible misalignment. The filtered dataset included alignments for 91 genes with a total alignment length of 106,566 bp and a completeness of 92.8%.

We inferred the species tree of Gadinae with StarBEAST2[59, 110] under the multi-species coalescent model, assuming a strict clock, constant population sizes, and the birth-death tree model[111], and averaging over substitution models with the bModelTest package[112] for BEAST 2. For time calibration, we placed lognormal prior distributions on the age of the divergence of the outgroup *Brosme brosme* from Gadinae (mean in real space: 32.325; standard deviation: 0.10) and on the crown age of Gadinae (mean in real space: 18.1358; standard deviation: 0.28); in both cases, the distribution parameters were chosen to approximate the age estimates for these two divergence events obtained by Musilova et al.[103]. We performed five replicate StarBEAST2 analyses, each with a length of one billion Markov-chain Monte Carlo (MCMC) iterations. After merging replicate posterior distributions, effective sample sizes (ESS) for all model parameters were greater than 1,000, indicating full stationarity and convergence of MCMC chains. We then used TreeAnnotator from the BEAST 2 package to summarize the posterior tree distribution in the form of a maximum-clade-credibility (MCC) consensus tree with Bayesian posterior probabilities as node support[113].

### Estimating divergence times and introgression among species of the genera *Gadus, Arctogadus*, and *Boreogadus*

To further investigate divergence times and introgression among species of the closely related genera *Gadus, Arctogadus*, and *Boreogadus*, we used a second phylogenomic dataset based on read mapping to the gadMor2 assembly. This dataset included Illumina read data for all four species of the genus *Gadus* (*G. morhua, G. chalcogrammus, G. macrocephalus*, and *G. ogac*)[50, 104], *Arctogadus glacialis* [104], and *Boreogadus saida*[104], as well as *Merlangius merlangius, Melanogrammus aeglefinus*, and *Pollachius virens*[56, 104], which we here considered outgroups (Supplementary Table 5). Read data from a stationary and a migratory indivual (both sampled at the Lofoten islands) were used to represent *Gadus morhua*. Mapping, read sorting, and indexing were again performed with BWA MEM and SAMtools, and variant calling was again performed with GATK’s HaplotypeCaller and GenotypeGVCFs tools as described above except that we now also exported invariant sites to the output file. To limit the dataset to the most reliably mapping genomic regions, we applied the mask of unreliable sites generated from the threeway whole-genome alignment (see above), resulting in set of 19,035,318 SNPs. We then extracted alignments from GATK’s output files for each non-overlapping window of 5,000 bp for which no more than 4,000 sites were masked, setting all genotypes with a Phred-scaled likelihood below 20 to missing. Alignments were not extracted from the four supergene regions and those windows with less than 100 variable sites were ignored. As we did not model recombination within alignments in our phylogenomic inference, the most suitable alignments for the inference were those with weak signals of recombination. Therefore, we calculated the number of hemiplasies per alignment by comparing the number of variable sites with the parsimony score, estimated with PAUP*[114], and excluded all alignments that had more than ten hemiplasies. Finally, we again removed all alignment sites for which BMGE determined a gap rate greater than 0.2 or an entropy-like score greater than 0.5. The resulting filtered dataset was composed of 109 alignments with a total length of 383,727 bp and a completeness of 91.0%.

We estimated the species tree and introgression among *Gadus, Arctogadus*, and *Boreogadus* under the isolation-with-migration model implemented in the AIM package[58] for BEAST 2. The inference assumed a strict clock, constant population sizes, the pure-birth tree model[115], and the HKY[116] substitution model with gamma-distributed rate variation among sites[117]. We time-calibrated the species tree with a single lognormal prior distribution on the divergence of *Pollachius virens* from all other taxa of the dataset (mean in real space: 8.56; standard deviation: 0.08), constraining the age of this divergence event according to the results of the analysis of divergence times of Gadinae (see above; Supplementary Figure 10). We performed ten replicate analyses that each had a length of five billion MCMC iterations, resulting in ESS values greater than 400 for all model parameters. The posterior tree distribution was subdivided according to tree topology and inferred gene flow, and we produced separate MCC consensus trees for each of the tree subsets.

To further test for introgression among *Gadus, Arctogadus*, and *Boreogadus*, we calculated Patterson’s *D*-statistic from the masked dataset for all possible species trios (with *Pollachius virens* fixed as outgroup) using the “Dtrios” function of Dsuite v.0.1.r3[63]. For the calculation of the *D*-statistic, species trios were sorted in two ways; with a topology fixed according to the species tree inferred under the isolation-with-migration model (*D*_fix_), and so that the number of “BBAA” patterns was greater than those of “ABBA” and “BABA” patterns (*D*_BBAA_). The significance of the statistic was assessed through block-jackknifing with 20 blocks of equal size. For the trios with the most significant signals of introgression, we further used the “Dinvestigate” function of Dsuite to calculate the *f*_dM_-statistic[60] within sliding windows of 50 SNPs, overlapping by 25 SNPs.

To corroborate the introgression patterns inferred with Dsuite, we performed two analyses based on comparisons of the frequencies of trio topologies in maximum-likelihood phylogenies. Alignments for these analyses were selected as for the species-tree inference under the isolation-with-migration model, except that up to 20 hemiplasies were allowed per alignment. The resulting set of 851 alignments had a total length of 3,052,697 bp and a completeness of 91.0%. From each of these alignments, a maximum-likelihood phylogeny was inferred with IQ-TREE v.1.6.8[118] with a substitution model selected through IQ-TREE’s standard model selection. Branches with a length below 0.001 were collapsed into polytomies. Based on the inferred maximum-likelihood trees, we calculated, for all possible species trios, the *D*_tree_-statistic of Ronco et al.[65], a tree-based equivalent to Patterson’s *D*-statistic in which the frequencies of pairs of sister taxa are counted in a set of trees instead of the frequencies of shared sites in a genome (a related measure was proposed by Huson et al.[119]): *D*_tree_ = (*f*_2nd_ *− f*_3rd_)*/*(*f*_2nd_ + *f*_3rd_), where for a given trio, *f*_2nd_ is the frequency of the second-most frequent pair of sisters and *f*_3rd_ is the frequency of the third-most frequent (thus, the least frequent) pair of sisters. As a second tree-based analysis of introgression, we applied genealogy interrogation[66], comparing the likelihoods of trees with alternative topological constraints for the same alignment, as in Barth et al.[67]. We tested two hypotheses of introgression with this method: 1) Introgression between *Arctogadus glacialis* and either *Boreogadus saida* or the group of the four species of the genus *Gadus*; and 2) introgression between *Gadus ogac* and the sister species *Gadus chalcogrammus* and *Gadus morhua*.

### Estimating divergence times, demography, and gene flow among *Gadus morhua* populations

To investigate divergence times among *Gadus morhua* populations, we applied phylogenetic analyses to the dataset based on whole-genome sequencing and variant calling for 24 *Gadus morhua* individuals (Supplementary Table 8). This dataset included, now considered as outgroups, the same representatives of *Gadus chalcogrammus, G. macrocephalus, G. ogac, Arctogadus glacialis*, and *Boreogadus saida* as our analyses of divergence times and introgression among *Gadus, Arctogadus*, and *Boreogadus* (see above). “Migratory” and “stationary” *Gadus morhua* individuals from Newfoundland, Iceland, Lofoten, and Møre were used as separate groups in these analyses. Subsequent to mapping with BWA MEM and variant calling with GATK’s HaplotypeCaller and GenotypeGVCFs tools, we filtered the called variants with BCFtools to include only sites for which the Phred-scaled *p* value for Fisher’s exact test was smaller than 20, the quality score normalised by read depth was greater than 2, the root-mean-square mapping quality was greater than 20, the overall read depth across all individuals was between the 10 and 90% quantiles, and the inbreeding coefficient was greater than -0.5. We further excluded sites if their Mann-Whitney-Wilcoxon rank-sum test statistic was smaller than -0.5 either for site position bias within reads or for mapping quality bias between reference and alternative alleles. After normalizing indels with BCFtools, SNPs in proximity to indels were discarded with a filter that took into account the length of the indel: SNPs were removed within 10 bp of indels that were 5 bp or longer, but only within 5, 3, or 2 bp if the indel was 3–4, 2, or 1 bp long, respectively. After applying this filter, all indels were removed from the dataset. For the remaining SNPs, genotypes with a read depth below 4 or a genotype quality below 20 were set to missing. Finally, we excluded all sites that were no longer variable or had more than two different alleles; the filtered dataset then contained 20,402,423 biallelic SNPs.

We inferred the divergence times among *Gadus morhua* populations from the SNP data under the multi-species coalescent model with the SNAPP add-on package for BEAST 2. Due to the high computational demand of SNAPP, we performed this analysis only with a further reduced set of 1,000 SNPs, randomly selected from all biallelic SNPs that were without missing genotypes and located outside of the supergene regions. The input files for SNAPP were prepared with the script snapp prep.rb[70], implementing a strict-clock model and a pure-birth tree model. The tree of *Gadus morhua* populations and outgroup species was time-calibrated with a single lognormal prior distribution (mean in real space: 3.83; standard deviation: 0.093) that constrained the root age of the tree according to the results of the analysis of divergence times and introgression among *Gadus, Arctogadus*, and *Boreogadus* (see above; Fig. 2b, Supplementary Figure 3). We performed three replicate SNAPP analyses, each with a length of 400,000 MCMC iterations, resulting in ESS values that were all greater than 400. The posterior tree distribution was again summarised as a MCC consensus tree.

Gene flow among *Gadus morhua* populations and outgroup species was investigated with Dsuite from all biallelic SNPs that were without missing genotypes and located outside of the four supergene regions; there were 408,574 of these. The gene flow analyses were performed with Dsuite’s “Dtrios” function as described above.

Population sizes over time were estimated for all sampled *Gadus morhua* populations with Relate v.1.1.2[73]. To maximize the number of suitable SNPs for this analysis, we excluded all outgroups except the sister species, *Gadus chalcogrammus*, and repeated variant calling and filtering with the same settings as before. After applying a mask to exclude all variants from repetitive regions in the gadMor2 assembly (784,488 bp in total)[31], 10,872,496 biallelic SNPs remained and were phased with BEAGLE v.5.1[120], setting the population size assumed by BEAGLE to 10,000. We excluded all sites that were heterozygous in the *Gadus chalcogrammus* individual and then reconstructed an “ancestral” genome sequence from the gadMor2 assembly and the called variants for *G. chalcogrammus*. Following this reconstruction, we removed *G. chalcogrammus* from the set of SNPs and excluded all sites that had become monomorphic after this removal, leaving 7,101,144 SNPs that were biallelic among the sampled *Gadus morhua* individuals. In addition to the “ancestral” genome sequence and the set of biallelic SNPs, we prepared a mask for the Relate analysis, covering all sites that were also included in the mask for repetitive regions, all sites that would have been excluded from variant calling due to proximity to indels (see above), and all sites that were ignored in the reconstruction of the “ancestral” sequence due to heterozygous genotype calls for the *G. chalcogrammus* individual.

As Relate further requires an estimate of the mutation rate, we calculated this rate for the filtered set of SNPs as the mean number of substitutions between *Gadus morhua* individuals from the Northwest Atlantic (thus, from the populations Newfoundland and Labrador) and those from the Northeast Atlantic (thus, from all other populations), divided by 2 times the expected coalescence time between the two groups and the genome size. We excluded the four linkage groups carrying supergenes from this calculation. The expected coalescence time was calculated as the divergence time between the two groups, which was estimated in the analysis with SNAPP as 65,400 years (Fig. 3), plus the expected time to coalescence within the common ancestor, which is the product of the generation time and the diploid population size under the assumption of a panmictic ancestral population. With an assumed generation time of 10 years[121] and a population size of 57,400, as estimated in the SNAPP analysis, the expected time to coalescence within the common ancestor is 574,000 years, and the total expected coalescence time was thus set to 65, 400 + 574, 000 = 639, 400 years. As the mean number of substitutions between the individuals of the two groups was 878,704.31 and the size of the gadMor2 assembly without LGs 1, 2, 7, and 12, and excluding masked sites, is 419,183,531 bp, the calculated mutation rate was *µ* = 878, 704.31*/*(2 × 639, 400 × 419, 183, 531) = 1.64 *×* 10^−9^ per bp and per year, or 1.64 × 10^−8^ per bp and per generation. Because the number of substitutions was calculated from the filtered set of SNPs, this rate is likely to underestimate the true mutation rate of *Gadus morhua*; however, because the same filtered set of SNPs was used as input for Relate, this rate is applicable in our inference of population sizes over time. The input file was converted from variant call format to haplotype format using RelateFileFormats with the flag “–mode ConvertFromVcf”. The script PrepareInputFiles.sh was used to flip genotypes according to the reconstructed “ancestral” genome sequence and to adjust distances between SNPs using the mask prepared for this analysis. Relate was first run to infer genome-wide genealogies and mutations assuming the above calculated mutation rate of 1.64 × 10^−8^ per bp and per generation and a diploid effective population size of 50,000. This was followed by an estimation of population-size changes over time by running the script EstimatePopulationSize.sh for five iterations, applying the same mutation rate and setting the threshold to remove uninformative trees to 0.5. The tools and scripts RelateFileFormats, PrepareInputFiles.sh, and EstimatePopulationSize.sh are all distributed with Relate.

### Estimating divergence times, demography, and gene flow specific to supergenes

The analyses of divergence times, demography, and gene flow among *Gadus morhua* populations were repeated separately with SNPs from each of the four supergene regions on LGs 1, 2, 7, and 12. While the SNAPP analyses for these regions were again performed with reduced subsets of 1,000 SNPs per region, the data subsets used in analyses of gene flow with Dsuite comprised 11,474, 3,123, 10,4121, and 10,339 biallelic SNPs, and those used in the analyses of demography with Relate comprised 211,057, 71,046, 130,918, and 130,620 biallelic SNPs, respectively. The mutation rate used as input for these Relate analyses was identical to the one used for the analysis with genome-wide SNPs.

### Estimating population divergence times across the genome

In addition to the genome-wide and supergene-specific SNAPP analyses that used biallelic SNPs from the entire genome or the entire length of supergene regions, we also performed sliding-window SNAPP analyses across all linkage groups to quantify differences in population divergence times across the genome. Our motivation for these analyses was primarily to assess whether or not divergence times were homogeneous over the lengths of supergenes, as differences in these divergence times within supergenes could be informative both about the presence of separate inversion within these regions and about their erosion processes. Additionally, we expected that these analyses could reveal further putative inversions elsewhere in the genome if they should exist.

From the set of 20,402,423 biallelic SNPs, we extracted subsets of SNPs for each non-overlapping window of a length of 250,000 bp, with a minimum distance between SNPs of 50 bp. We discarded windows with less than 500 remaining biallelic SNPs and used a maximum of 1,000 biallelic SNPs per window; these were selected at random if more biallelic SNPs were available per window. Input files for SNAPP were then prepared as for the genome-wide and supergene-specific SNAPP analyses. Per window, we performed two replicate SNAPP analyses with an initial length of 100,000 MCMC iterations, and these analyses were resumed up to a maximum of 500,000 MCMC iterations as long as the lowest ESS value was below 100. Windows with less than 300 sufficiently complete SNPs for SNAPP analyses, with an ESS value below 100 after the maximum number of MCMC iterations, or with a mean BPP node support value below 0.5 were discarded after the analysis. Per remaining window, posterior tree distributions from the two replicate analyses were combined and summarised in the form of MCC consensus trees. Additionally, a random sample of 100 trees was drawn from each combined posterior distribution.

Instead of showing all resulting trees, we developed a type of plot that shows, without loss of phylogenetic information, the divergence times stacked upon each other on a single axis, which allowed us to illustrate these divergence times efficiently across linkage groups. For this plot, all trees were first ladderised, outgroups were pruned, and the divergence times between each pair of populations adjacent to each other on the ladderised trees were extracted. Per window, the order of populations on the ladderised tree, together with the extracted divergence times between them, was used to define the positions of points on the vertical axis of the plot, so that each point represents a population and their vertical distances indicate the divergence times between populations that are next to each other on the ladderised tree. The positions of window on the linkage group were used to place these dots on the horizontal axis of the plot, and all dots representing the same population were connected by lines to produce the complete plot of divergence times across linkage groups.

## Code availability

Code for computational analyses is available from Github (http://github.com/mmatschiner/super-genes).

## Data availability

The gadMor_Stat assembly (ENA accession GCA_905250895) and read data for all *Gadus morhua* specimens listed in Supplementary Table 8 are deposited on ENA with project number PRJEB43149. Alignment files, SNP datasets in PED and VCF format, and input and output of phylogenetic analyses are available from Zenodo (doi: 10.5281/zenodo.4560275).

## Acknowledgements

We thank M. Malmstrøm, P. Berg, and D. Righton for help with fieldwork, and M. Skage, S. Kollias, M. S. Hansen, and A. Tooming-Klunderud from the Norwegian Sequencing Centre (NSC; https://www.sequencing.uio.no) for sequencing and processing of samples. Aliaksandr Hubin and Geir Storvik helped with initial analyses. PacBio and Illumina library creation and high-throughput sequencing were carried out at NSC, University of Oslo, Norway. We also thank Alexandra Viertler for drawings of codfishes, Côme Denechaud and the Institute for Marine Research, Norway, for providing otolith images, and Ethan Schoonover for developing the Solarized color palette (https://ethanschoonover.com/solarized/). All computational analyses were performed on the Abel and Saga supercomputing clusters (Norwegian metacenter for High Performance Computing and the University of Oslo) operated by the Research Computing Services group at USIT, the University of Oslo IT-department, and by UNINETT Sigma2, the National Infrastructure for High Performance Computing and Data Storage in Norway. This work was funded by the Research Council of Norway (RCN) through “The Aqua Genome Project” (221734/O30) to K.S.J.

## Author contributions

M.M., K.S.J., and S.J. conceived this study. M.M. performed most analyses. J.M.I.B. contributed demographic analyses, O.K.T. produced the gadMor_Stat assembly, and B.S. performed variant calling. H.T.B. and M.S.O.B. contributed to the organization of the study, and K.S.J. and S.J. arranged whole-genome sequencing. C.P. and I.B. provided samples for sequencing. M.M. wrote the manuscript, with individual sections contributed by J.M.I.B. and O.K.T. All authors provided feedback and approved the final version of the manuscript.

## Competing interests

The authors declare no competing interests.

## Notes

https://zenodo.org/record/4560275#.YDvriy2ZN24

https://github.com/mmatschiner/supergenes

### Competing Interest Statement

The authors have declared no competing interest.

## References

1. Joron, M. et al. Chromosomal rearrangements maintain a polymorphic supergene controlling butterfly mimicry. Nature 477, 203–206 (2011).

2. Yan, Z. et al. Evolution of a supergene that regulates a trans-species social polymorphism. Nat. Ecol. Evol. 4, 210–249 (2020).

3. Lamichhaney, S. et al. Structural genomic changes underlie alternative reproductive strategies in the ruff (Philomachus pugnax). Nat. Genet. 48, 84–88 (2016).

4. Tuttle, E. M. et al. Divergence and functional degradation of a sex chromosome-like supergene. Curr. Biol. 26, 344–350 (2016).

5. Li, J. et al. Genetic architecture and evolution of the S locus supergene in Primula vulgaris. Nat. Plants 2, 16188 (2016).

6. Thompson, M. J. & Jiggins, C.D. Supergenes and their role in evolution. Heredity 113, 1–8 (2014).

7. Schwander, T., Libbrecht, R. & Keller, L. Supergenes and complex phenotypes. Curr. Biol. 24, R288–R294 (2014).

8. Tigano, A. & Friesen, V. L. Genomics of local adaptation with gene flow. Mol. Ecol. 25, 2144–2164 (2016).

9. Gutiérrez-Valencia, J., Hughes, P. W., Berdan, E. L. & Slotte, T. The genomic architecture and evolutionary fates of supergenes. Genome Biol. Evol. 13 (2021).

10. Fisher, R. A. The genetical theory of natural selection (Clarendon Press, Oxford, UK, 1930).

11. Kirkpatrick, M. Chromosome inversions, local adaptation and speciation. Genetics 173, 419–434 (2006).

12. Jay, P. et al. Supergene evolution triggered by the introgression of a chromosomal inversion. Curr. Biol. 28, 1839–1845.e3 (2018).

13. Jay, P., Aubier, T. G. & Joron, M. Admixture can readily lead to the formation of supergenes. bioRxiv. doi:10.1101/2020.11.19.389577 (2020).

14. Dobzhansky, T. & Epling, C. The suppression of crossing over in inversion heterozygotes of Drosophila pseudoobscura. Proc. Natl. Acad. Sci. U.S.A. 34, 137–141 (1948).

15. Sturtevant, A. H. & Beadle, G. W. The relations of inversions in the X chromosome of Drosophila melanogaster to crossing over and disjunction. Genetics 21, 554–604 (1936).

16. Anton, E., Blanco, J., Egozcue, J. & Vidal, F. Sperm studies in heterozygote inversion carriers: a review. Cytogenet. Genome Res. 111, 297–304 (2005).

17. Navarro, A., Barbadilla, A. & Ruiz, A. Effect of inversion polymorphism on the neutral nucleotide variability of linked chromosomal regions in Drosophila. Genetics 155, 685–698 (2000).

18. Faria, R., Johannesson, K., Butlin, R. K. & Westram, A. M. Evolving inversions. Trends Ecol. Evol. 34, 239–248 (2019).

19. Berdan, E. L., Blanckaert, A., Butlin, R. K. & Bank, C. Deleterious mutation accumulation and the long-term fate of chromosomal inversions. PLOS Genet. 17, e1009411 (2021).

20. Andolfatto, P., Depaulis, F. & Navarro, A. Inversion polymorphisms and nucleotide variability in Drosophila. Genet. Res. 77, 1–8 (2001).

21. Chovnick, A. Gene conversion and transfer of genetic information within the inverted region of inversion heterozygotes. Genetics 75, 123–131 (1973).

22. Chen, J.-M., Cooper, D. N., Chuzhanova, N., Férec, C. & Patrinos, G. P. Gene conversion: mechanisms, evolution and human disease. Nat. Rev. Genet. 8, 762–775 (2007).

23. Jeffreys, A. J. & May, C. A. Intense and highly localized gene conversion activity in human meiotic crossover hot spots. Nat. Genet. 36, 151–156 (2004).

24. Williams, A. L. et al. Non-crossover gene conversions show strong GC bias and unexpected clustering in humans. eLIFE 4, e04637 (2015).

25. Figuet, E., Ballenghien, M., Romiguier, J. & Galtier, N. Biased gene conversion and GCcontent evolution in the coding sequences of reptiles and vertebrates. Genome Biol. Evol. 7, 240–250 (2014).

26. Korunes, K. L. & Noor, M. A. F. Pervasive gene conversion in chromosomal inversion heterozygotes. Mol. Ecol. 28, 1302–1315 (2019).

27. Bradbury, I. R. et al. Long distance linkage disequilibrium and limited hybridization suggest cryptic speciation in Atlantic Cod. PLOS ONE 9, e106380 (2014).

28. Berg, P. R. et al. Adaptation to low salinity promotes genomic divergence in Atlantic cod (Gadus morhua L.) Genome Biol. Evol. 7, 1644–1663 (2015).

29. Berg, P. R. et al. Three chromosomal rearrangements promote genomic divergence between migratory and stationary ecotypes of Atlantic cod. Sci. Rep. 6, 23246 (2016).

30. Sodeland, M. et al. “Islands of divergence” in the Atlantic cod genome represent polymorphic chromosomal rearrangements. Genome Biol. Evol. 8, 1012–1022 (2016).

31. Tørresen, O. K. et al. An improved genome assembly uncovers prolific tandem repeats in Atlantic cod. BMC Genomics 18, 95 (2017).

32. Kirubakaran, T. G. et al. Two adjacent inversions maintain genomic differentiation between migratory and stationary ecotypes of Atlantic cod. Mol. Ecol. 25, 2130–2143 (2016).

33. Kess, T. et al. A migration-associated supergene reveals loss of biocomplexity in Atlantic cod. Sci. Adv. 5, eaav2461 (2019).

34. Barney, B. T., Munkholm, C., Walt, D. R. & Palumbi, S. R. Highly localized divergence within supergenes in Atlantic cod (Gadus morhua) within the Gulf of Maine. BMC Genomics 18, 271 (2017).

35. Berg, P. R. et al. Trans-oceanic genomic divergence of Atlantic cod ecotypes is associated with large inversions. Heredity 119, 418–428 (2017).

36. Barth, J. M. I. et al. Genome architecture enables local adaptation of Atlantic cod despite high connectivity. Mol. Ecol. 26, 4452–4466 (2017).

37. Barth, J. M. I. et al. Disentangling structural genomic and behavioural barriers in a sea of connectivity. Mol. Ecol. 87, 449 (2019).

38. Berg, E. & Albert, O. T. Cod in fjords and coastal waters of North Norway: distribution and variation in length and maturity at age. ICES J. Mar. Sci. 60, 787–797 (2003).

39. Case, R., Hutchinson, W. F., Hauser, L., Van Oosterhout, C. & Carvalho, G. R. Macro- and micro-geographic variation in pantophysin (PanI) allele frequencies in NE Atlantic cod Gadus morhua. Mar. Ecol. Prog. Ser. 301, 267–278 (2005).

40. Star, B. et al. Ancient DNA reveals the Arctic origin of Viking Age cod from Haithabu, Germany. Proc. Natl. Acad. Sci. U.S.A. 114, 9152–9157 (2017).

41. Nordeide, J. T. & Folstad, I. Is cod lekking or a promiscuous group spawnerã Fish Fish. 1, 90–93 (2000).

42. Kirubakaran, T. G. et al. A nanopore based chromosome-level assembly representing Atlantic cod from the Celtic Sea. G3, g3.401423.2020 (2020).

43. Puncher, G. N. et al. Life-stage-dependent supergene haplotype frequencies and metapopulation neutral genetic patterns of Atlantic cod, Gadus morhua, from Canada’s Northern cod stock region and adjacent areas. J. Fish Biol. 98, 817–828 (2021).

44. Johansen, T. et al. Genomic analysis reveals neutral and adaptive patterns that challenge the current management regime for East Atlantic cod Gadus morhua L. Evol. Appl. 13, 2673–2688 (2020).

45. Kess, T. et al. Modular chromosome rearrangements reveal parallel and nonparallel adaptation in a marine fish. Ecol. Evol. 10, 638–653 (2020).

46. Hemmer-Hansen, J. et al. A genomic island linked to ecotype divergence in Atlantic cod. Mol. Ecol. 22, 2653–2667 (2013).

47. Carr, S. M., Kivlichan, D. S., Pepin, P. & Crutcher, D. C. Molecular systematics of gadid fishes: implications for the biogeographic origins of Pacific species. Can. J. Zool. 77, 19–26 (1999).

48. Coulson, M. W., Marshall, H. D., Pepin, P. & Carr, S. M. Mitochondrial genomics of gadine fishes: implications for taxonomy and biogeographic origins from whole-genome data sets. Genome 49, 1115–1130 (2006).

49. Bermingham, E., McCafferty, S. S. & Martin, A. P. in Molecular Systematics of Fishes (eds Kocher, T.D. & Stepien, C.A.) 113–128 (Academic Press, San Diego, USA, 1997).

50. Árnason, E. & Halldórsdóttir, K. Codweb: Whole-genome sequencing uncovers extensive reticulations fueling adaptation among Atlantic, Arctic, and Pacific gadids. Sci. Adv. 5, eaat8788 (2019).

51. Simão, F. A., Waterhouse, R. M., Ioannidis, P., Kriventseva, E. V. & Zdobnov, E. M. BUSCO: assessing genome assembly and annotation completeness with single-copy orthologs. Bioinformatics 31, 3210–3212 (2015).

52. Sturtevant, A. H. A case of rearrangement of genes in Drosophila. Proc. Natl. Acad. Sci. U.S.A. 7, 235–237 (1921).

53. Stevison, L. S., Hoehn, K. B. & Noor, M. A. F. Effects of inversions on within- and between-species recombination and divergence. Genome Biol. Evol. 3, 830–841 (2011).

54. Kirkpatrick, M. How and why chromosome inversions evolve. PLOS Biol. 8, e1000501 (2010).

55. Stransky, C. et al. Separation of Norwegian coastal cod and Northeast Arctic cod by outer otolith shape analysis. Fish. Res. 90, 26–35 (2008).

56. Tørresen, O. K. et al. Genomic architecture of haddock (Melanogrammus aeglefinus) shows expansions of innate immune genes and short tandem repeats. BMC Genomics 19, 240 (2018).

57. Edelman, N. B. et al. Genomic architecture and introgression shape a butterfly radiation. Science 366, 594–599 (2019).

58. Müller, N. F., Ogilvie, H. A., Zhang, C., Drummond, A. & Stadler, T. Inference of species histories in the presence of gene flow. bioRxiv. doi:10.1101/348391 (2018).

59. Bouckaert, R. R. et al. BEAST 2.5: An advanced software platform for Bayesian evolutionary analysis. PLOS Comput. Biol. 15, e1006650 (2019).

60. Malinsky, M. et al. Genomic islands of speciation separate cichlid ecomorphs in an East African crater lake. Science 350, 1493–1498 (2015).

61. Green, R. E. et al. A draft sequence of the Neandertal genome. Science 328, 710–722 (2010).

62. Durand, E. Y., Patterson, N., Reich, D. & Slatkin, M. Testing for ancient admixture between closely related populations. Mol. Biol. Evol. 28, 2239–2252 (2011).

63. Malinsky, M., Matschiner, M. & Svardal, H. Dsuite - Fast D-statistics and related admixture evidence from VCF files. Mol. Ecol. Resour. 19, 1655 (2020).

64. Martin, S. H., Davey, J. W. & Jiggins, C. D. Evaluating the use of ABBA-BABA statistics to locate introgressed loci. Mol. Biol. Evol. 32, 244–257 (2015).

65. Ronco, F. et al. Drivers and dynamics of a massive adaptive radiation in cichlid fishes. Nature 589, 76–81 (2021).

66. Arcila, D. et al. Genome-wide interrogation advances resolution of recalcitrant groups in the tree of life. Nat. Ecol. Evol. 1, 1–10 (2017).

67. Barth, J. M. I. et al. Stable species boundaries despite ten million years of hybridization in tropical eels. Nat. Commun. 11, 1433 (2020).

68. Robichaud, D. & Rose, G. A. Migratory behaviour and range in Atlantic cod: inference from a century of tagging. Fish Fish. 5, 185–214 (2004).

69. Bryant, D., Bouckaert, R. R., Felsenstein, J., Rosenberg, N. A. & RoyChoudhury, A. Inferring species trees directly from biallelic genetic markers: bypassing gene trees in a full coalescent analysis. Mol. Biol. Evol. 29, 1917–1932 (2012).

70. Stange, M., Sánchez-Villagra, M. R., Salzburger, W. & Matschiner, M. Bayesian divergence-time estimation with genome-wide SNP data of sea catfishes (Ariidae) supports Miocene closure of the Panamanian Isthmus. Syst. Biol. 67, 681–699 (2018).

71. Bradbury, I. R. et al. Parallel adaptive evolution of Atlantic cod on both sides of the Atlantic Ocean in response to temperature. Proc. R. Soc. Lond. B 277, 3725–3734 (2010).

72. Ruegg, K., Anderson, E. C., Boone, J., Pouls, J. & Smith, T. B. A role for migration-linked genes and genomic islands in divergence of a songbird. Mol. Ecol. 23, 4757–4769 (2014).

73. Speidel, L., Forest, M., Shi, S. & Myers, S. R. A method for genome-wide genealogy estimation for thousands of samples. Nat. Genet. 1321–1329 (2019).

74. Navarro, A., Betrán, E., Barbadilla, A. & Ruiz, A. Recombination and gene flux caused by gene conversion and crossing over in inversion heterokaryotypes. Genetics 146, 695–709 (1997).

75. Reis, M., Vieira, C. P., Lata, R., Posnien, N. & Vieira, J. Origin and consequences of chromosomal inversions in the virilis group of Drosophila. Genome Biol. Evol. 10, 3152–3166 (2018).

76. Thorsen, A., Kjesbu, O. S., Fyhn, H. J. & Solemidal, P. Physiological mechanisms of buoyancy in eggs from brackish water cod. J. Fish Biol. 48, 457–477 (1996).

77. Matsubara, T. et al. Multiple vitellogenins and their unique roles in marine teleosts. Fish Physiol. Biochem. 28, 295–299 (2003).

78. Braasch, I. & Salzburger, W. In ovo omnia: diversification by duplication in fish and other vertebrates. J. Biol. 8, 1–5 (2009).

79. Finn, R. N. & Fyhn, H. J. Requirement for amino acids in ontogeny of fish. Aquac. Res. 41, 684–716 (2010).

80. Runemark, A. et al. Variation and constraints in hybrid genome formation. Nat. Ecol. Evol. 2, 549–556 (2018).

81. Arbeithuber, B., Betancourt, A. J., Ebner, T. & Tiemann-Boege, I. Crossovers are associated with mutation and biased gene conversion at recombination hotspots. Proc. Natl. Acad. Sci. USA 112, 2109–2114 (2015).

82. Finn, R. N., Kolarevic, J., Kongshaug, H. & Nilsen, F. Evolution and differential expression of a vertebrate vitellogenin gene cluster. BMC Evol. Biol. 9, 2 (2009).

83. Westin, L. & Nissling, A. Effects of salinity on spermatozoa motility, percentage of fertilized eggs and egg development of Baltic cod (Gadus morhua), and implications for cod stock fluctuations in the Baltic. Mar. Biol. 108, 5–9 (1991).

84. Hüssy, K. Review of western Baltic cod (Gadus morhua) recruitment dynamics. ICES J. Mar. Sci. 68, 1459–1471 (2011).

85. Johannesson, K., Smolarz, K., Grahn, M. & André, C. The future of Baltic Sea populations: local extinction or evolutionary rescueã AMBIO 40, 179–190 (2011).

86. Nissling, A., Kryvi, H. & Vallin, L. Variation in egg buoyancy of Baltic cod Gadus morhua and its implications for egg survival in prevailing conditions in the Baltic Sea. Mar. Ecol. Prog. Ser. 110, 67–74 (1994).

87. Nissling, A. & Westin, L. Salinity requirements for successful spawning of Baltic and Belt Sea cod and the potential for cod stock interactions in the Baltic Sea. Mar. Ecol. Prog. Ser. 152, 261–271 (1997).

88. Hess, J. E. et al. Genomic islands of divergence infer a phenotypic landscape in Pacific lamprey. Mol. Ecol. 29, 3841–3856 (2020).

89. Schaeffer, S. W. & Anderson, W. W. Mechanisms of genetic exchange within the chromosomal inversions of Drosophila pseudoobscura. Genetics 171, 1729–1739 (2005).

90. Miller, J. R. et al. Aggressive assembly of pyrosequencing reads with mates. Bioinformatics 24, 2818–2824 (2008).

91. Chin, C.-S. et al. Nonhybrid, finished microbial genome assemblies from long-read SMRT sequencing data. Nat. Methods 10, 563–569 (2013).

92. Li, H. & Durbin, R. Fast and accurate long-read alignment with Burrows-Wheeler transform. Bioinformatics 26, 589–595 (2010).

93. Li, H. et al. The Sequence Alignment/Map format and SAMtools. Bioinformatics 25, 2078–2079 (2009).

94. Li, H. A statistical framework for SNP calling, mutation discovery, association mapping and population genetical parameter estimation from sequencing data. Bioinformatics 27, 2987–2993 (2011).

95. Walker, B. J. et al. Pilon: An integrated tool for comprehensive microbial variant detection and genome assembly improvement. PLOS ONE 9, e112963 (2014).

96. McKenna, A. et al. The Genome Analysis Toolkit: A MapReduce framework for analyzing next-generation DNA sequencing data. Genome Res. 20, 1297–1303 (2010).

97. Poplin, R. et al. Scaling accurate genetic variant discovery to tens of thousands of samples. bioRxiv. doi:10.1101/201178 (2018).

98. Danecek, P. et al. The variant call format and VCFtools. Bioinformatics 27, 2156–2158 (2011).

99. Purcell, S. et al. PLINK: A tool set for whole-genome association and population-based linkage analyses. Am. J. Hum. Genet. 81, 559–575 (2007).

100. Altschul, S. F., Gish, W., Miller, W., Myers, E. W. & Lipman, D. J. Basic local alignment search tool. J. Mol. Biol. 215, 403–410 (1990).

101. Sandes, E. F. d. O., Miranda, G., Melo, A. C. M. A. d., Martorell, X. & Ayguade, E. CUDAlign 3.0: Parallel Biological Sequence Comparison in Large GPU Clusters. 2014 14th IEEE/ACM International Symposium on Cluster, Cloud and Grid Computing (CCGrid), 160–169 (2014).

102. Harris, R. S. Improved pairwise alignment of genomic DNA. PhD thesis (Pennsylvania State University, 2007).

103. Musilova, Z. et al. Vision using multiple distinct rod opsins in deep-sea fishes. Science 364, 588–592 (2019).

104. Malmstrøm, M. et al. Evolution of the immune system influences speciation rates in teleost fishes. Nat. Genet. 48, 1204–1210 (2016).

105. Roth, O. et al. Evolution of male pregnancy associated with remodeling of canonical vertebrate immunity in seahorses and pipefishes. Proc. Natl. Acad. Sci. U.S.A. 117, 9431–9439 (2020).

106. Kucuk, E. et al. Kollector: transcript-informed, targeted de novo assembly of gene loci. Bioinformatics 33, 1782–1788 (2017).

107. Criscuolo, A. & Gribaldo, S. BMGE (Block Mapping and Gathering with Entropy): a new software for selection of phylogenetic informative regions from multiple sequence alignments. BMC Evol. Biol. 10, 210 (2010).

108. Stamatakis, A. RAxML version 8: a tool for phylogenetic analysis and post-analysis of large phylogenies. Bioinformatics 30, 1312–1313 (2014).

109. Leigh, J. W., Susko, E., Baumgartner, M. & Roger, A. J. Testing congruence in phylogenomic analysis. Syst. Biol. 57, 104–115 (2008).

110. Ogilvie, H. A., Bouckaert, R. R. & Drummond, A. J. StarBEAST2 brings faster species tree inference and accurate estimates of substitution rates. Mol. Biol. Evol. 34, 2101–2114 (2017).

111. Gernhard, T. The conditioned reconstructed process. J. Theor. Biol. 253, 769–778 (2008).

112. Bouckaert, R. R. & Drummond, A. J. bModelTest: Bayesian phylogenetic site model averaging and model comparison. BMC Evol. Biol. 17, 42 (2017).

113. Heled, J. & Bouckaert, R. R. Looking for trees in the forest: summary tree from posterior samples. BMC Evol. Biol. 13, 221 (2013).

114. Swofford, D. L. PAUP*. Phylogenetic Analysis Using Parsimony (*and other methods). Version 4. (2003).

115. Yule, G. U. A mathematical theory of evolution, based on the conclusions of Dr. J. C. Willis, F.R.S. Phil. Trans. R. Soc. B 213, 21–87 (1925).

116. Hasegawa, M., Kishino, H. & Yano, T. Dating of the human-ape splitting by a molecular clock of mitochondrial DNA. J. Mol. Evol. 22, 160–174 (1985).

117. Yang, Z. Maximum likelihood phylogenetic estimation from DNA sequences with variable rates over sites: approximate methods. J. Mol. Evol. 39, 306–314 (1994).

118. Nguyen, L.-T., Schmidt, H. A., Von Haeseler, A. & Minh, B. Q. IQ-TREE: A fast and effective stochastic algorithm for estimating maximum-likelihood phylogenies. Mol. Biol. Evol. 32, 268–274 (2015).

119. Huson, D. H., Klöpper, T., Lockhart, P. J. & Steel, M. A. in Research in Computational Molecular Biology. RECOMB 2005. Lecture Notes in Computer Science (eds Miyano, S.et al.) 233–249 (Springer, Berlin, Heidelberg, 2005).

120. Browning, S. R. & Browning, B. L. Rapid and accurate haplotype phasing and missing-data inference for whole-genome association studies by use of localized haplotype clustering. Am. J. Hum. Genet. 81, 1084–1097 (2007).

121. Smedbol, R. K., Shelton, P. A., Fréchet, A. & Chouinard, G. A. Review of population structure, distribution and abundance of cod (Gadus morhua) in Atlantic Canada in a species-at-risk context. Canadian Science Advisory Secretariat Research Document 2002/082, 1–134 (2002).

