## Supplementary Information for "Supergene origin and maintenance in Atlantic cod"

**Matschiner et al.**

### Supplementary Notes

**Supplementary Note 1:** Unreliable support for introgression in the study by Árnason and Halldórsdóttir (2019).

In their study, Árnason and Halldórsdóttir[1] investigated signals of introgression among species of the genera *Gadus*, *Arctogadus*, and *Boreogadus* using a combination of low-coverage (3×) and high-coverage ($12-45\times$) whole-genome-sequencing data. The authors reported discordance between mitochondrial and nuclear phylogenies, elevated $D$-statistics, and reticulations inferred with PhyloNetworks[2], and interpreted their results as support for i) introgression between *Gadus morhua* and *Gadus ogac*, ii) strong genetic divergence between eastern and western populations of *Arctogadus glacialis*, iii) the (past) existence of ghost lineages as pathways for introgression, iv) introgression between *Gadus morhua* and *Gadus chalcogrammus* or a homoploid hybrid speciation origin of the latter. As we argue below, these conclusions are at least partially unreliable due to mislabelled sequences, reference bias, and incorrect application of statistical tests.

*Mislabelled sequences and reference bias generated tree discordance.* Árnason and Halldórsdóttir inferred phylogenies using the Neighbor-Joining algorithm[3], implemented in ape[4], from both mitochondrial sequence alignments and nuclear single-nucleotide-polymorphism (SNP) data (Fig. 2 in their study). The resulting trees were discordant with each other because the placement of *Arctogadus glacialis* differed between them. Árnason and Halldórsdóttir interpreted this discordance as evidence for introgression into *Arctogadus glacialis*, the (past) existence of ghost lineages, and — because a second added mitochondrial sequence for *Arctogadus glacialis* did not cluster with the first — strong divergence between separate populations of *Arctogadus glacialis*. Notably, however, both the mitochondrial and the nuclear phylogenies inferred by Árnason and Halldórsdóttir also differed from the consensus view of the relationships among the investigated species, which has been supported rather unambiguously, both by mitochondrial and nuclear datasets, and in a range of publications[5–9]. According to that consensus view, *Gadus morhua* is the sister species to *Gadus chalcogrammus*, *Gadus macrocephalus* is the sister species to *Gadus ogac*, the two pairs of sister species are closer to each other than either is to *Arctogadus glacialis* or *Boreogadus saida*, and the clade formed by all of these six species is monophyletic. In the mitochondrial tree inferred by Árnason and Halldórsdóttir, on the other hand, the sampled *Arctogadus glacialis* specimen clustered with an outgroup, *Micromesistius poutassou*, whereas the same specimen clustered with *Gadus morhua* in their nuclear tree.

As the *Arctogadus glacialis* specimen used by Árnason and Halldórsdóttir was the one specimen sequenced to a far lower read depth (3×) than all other specimens ($12-45\times$) used in their study, and as they had mapped all reads towards the gadMor2[10] genome assembly for *Gadus morhua*, we assumed that reference bias could have led to a placement of *Arctogadus glacialis* near *Gadus morhua* in their nuclear phylogeny, which would explain at least the discordance between their nuclear phylogeny and the consensus view[11, 12]. To test this assumption, we reanalyzed the genomic read data produced by Árnason and Halldórsdóttir together with data from two other recent genomic studies of codfishes (family Gadidae)[8, 13]. We downloaded all accessions used by Árnason and Halldórsdóttir from the European Nucleotide Archive (ENA): SRR2906188 (*Arctogadus glacialis*), SRR2906185 (*Boreogadus saida*), SRR2906193 (*Gadus ogac*), SRR2906345 (*Gadus macrocephalus*), SRR2906339 (*G. chalcogrammus*), and SRR2906208, SRR2906256, SRR2906368, SRR2906362, SRR2906197, and SRR2906248 (all *G. morhua*). In addition, we downloaded the following accessions produced by Malmstrøm et al.[8]: ERR1473878 (*Pollachius virens*), ERR1473879 (*Melanogrammus aeglefinus*), ERR1473880 and ERR1473881 (*Merlangius merlangus*), ERR1473882 and ERR1473883 (*Arctogadus glacialis*), ERR1473884 and ERR1473885 (*Boreogadus saida*), ERR1473886 (*Gadus chalcogrammus*), ERR1473887, ERR1473888, ERR1473889, and ERR1473890 (all *Gadus morhua*), ERR1473875 (*Brosme brosme*), ERR1473876 (*Trisopterus minutus*), and ERR1473877 (*Gadiculus argenteus*). Finally, we added accession ERR1278928 (*Gadus ogac*) generated by Kirubakaran et al.[13]. To exlude reference bias in our own inference of the nuclear phylogeny, we applied local assembly with Kollector v.1.0.1[14] for a set of nuclear marker sequences from Roth et al.[15], instead of mapping the reads to the gadMor2 assembly. We also identified, aligned, and filtered orthologous sequences as in Roth et al.[15], and concatenated the resulting 210 alignments with a total length of 41,811 bp. From this concatenated alignment, we inferred the nuclear phylogeny with IQ-TREE v.1.6.8[16].

The inferred nuclear phylogeny confirmed our assumption of reference bias: The *Arctogadus glacialis* specimen from Árnason and Halldórsdóttir did not appear close to *Gadus morhua* as in their nuclear phylogeny. Instead, the two included *Gadus chalcogrammus* specimens appeared as the sister clade to a monophyletic group formed by all included *Gadus morhua* specimens and all other relationships also agreed with the consensus view, including the position of the *Arctogadus glacialis* specimen used by Malmstrøm et al., which appeared as the sister to the *Boreogadus saida* specimens. The one exception to this was the specimen identified as *Arctogadus glacialis* by Árnason and Halldórsdóttir[1] (ENA accession SRR2906188): Instead of clustering with the *Arctogadus glacialis* from Malmstrøm et al., this specimen appeared as the sister to the *Trisopterus minutus* specimen (accession ERR1473876). A reanalysis of mitochondrial sequences, extracted from the same set of accessions, produced the exact same topology, in which the *Arctogadus glacialis* specimen from Árnason and Halldórsdóttir once again appeared as the sister of the *Trisopterus minutus* specimen. As the genus *Trisopterus* is likely the sister lineage to the genus *Micromesistius*[17] (which was not included in our reanalysis), this result is entirely consistent with the position inferred by Árnason and Halldórsdóttir based on mitochondrial data, under the assumption that the specimen considered by them to be an *Arctogadus glacialis* specimen is mislabelled and in fact represents the species *Micromesistius poutassou*. This assumption of taxon mislabeling appears even more plausible as the mitochondrial sequence of the "*Arctogadus glacialis*" specimen is 99% identical to the mitochondrial sequence of *Micromesistius poutassou* that Árnason and Halldórsdóttir had obtained from GenBank (accession FR751401) and the two species *Arctogadus glacialis* and *Micromesistius poutassou* are phenotypically similar, grow to similar lengths, and occur both in the region sampled by Árnason and Halldórsdóttir. Thus, we conclude that instead of introgression, ghost lineages, and population separation in *Arctogadus glacialis*, it was reference bias in the low-coverage nuclear data, and mislabeling of a *Micromesistius poutassou* specimen as "*Arctogadus glacialis*" that caused the discordance between the mitochondrial and the nuclear trees inferred by Árnason and

Halldórsdóttir, as well as the discordance between these trees and the consensus view of relations among codfishes.

*Incorrect application led to exaggerated D-statistics.* To further investigate the occurrence of introgression among codfishes, Árnason and Halldórsdóttir applied Patterson's $D$-statistic[18, 19] to their nuclear SNP dataset, using ANGSD[20]. A $D$-statistic significantly different from zero supports introgression between two taxa in a quartet of taxa P1, P2, P3, and P4, if the four taxa are labelled so that P1 and P2 are sister taxa and P4 is the outgroup to the other three. In the study by Árnason and Halldórsdóttir, this basic requirement for the application of $D$-statistics was not met for 26 out of 30 tested quartets (Fig. 3 of their study), in part because of the use of the specimen mislabelled as "*Arctogadus glacialis*". All of these 26 quartets had produced highly significant $D$-statistics, but because the test requirement was not met, none of these significant $D$-statistics actually support introgression. Of the remaining four tested quartets, one produced an insignificant $D$-statistics and the last three seemed to provide weak support ($0.04 \leq D \leq 0.06$) for introgression between the two Pacific species *Gadus chalcogrammus* and *Gadus macrocephalus*, and between the two Atlantic species *Gadus morhua* and *Gadus ogac*. These inferred occurrences of introgression may be plausible given the overlapping distributions and close relationships of the species involved.

*Reference bias and incorrect root placement affected results of PhyloNetworks analyses.* Finally, Árnason and Halldórsdóttir used the SNaQ method implemented in PhyloNetworks[2] to infer trees in which reticulation edges were supposed to indicate introgression events (Figs. 4–6 and Supplementary Figures S2 and S3 in their study). This analysis was applied separately to SNP data from each of the 23 linkage groups of the gadMor2 assembly. The results of these analyses are difficult to interpret, as — contrary to expectations — not just the inferred reticulation edges but also the inferred species tree differed strongly among the linkage groups. A common feature to most of the presented trees was a reticulation edge connecting the mislabelled "*Arctogadus glacialis*" specimen, for which only low-coverage (3×) data had been available, with *Gadus morhua* specimens, indicating that, just like the comparison of mitochondrial and nuclear phylogenies, the SNaQ analyses were affected by reference bias in this particular specimen. Additionally, the presented trees suffer from incorrect root placement, given that the most divergent species of the dataset was not *Boreogadus saida*, but *Micromesistius poutassou*, represented by the specimen mislabelled as "*Arctogadus glacialis*".

**Supplementary Note 2:** Analyses of the detectability of extreme ancient bottlenecks.

To verify that signatures of extreme bottlenecks are detectable in descending genomes even after long periods of time, we first performed a series of coalescent simulations of a panmictic population that experienced a bottleneck during which the population size was reduced to a single sequence, using msprime v.0.7.4[21]. These simulations confirmed that, depending on the age of the bottleneck and the population size following the bottleneck, substantial proportions of the genome coalesce inside the bottleneck and are thus able to carry its signature: For example, 22% of the genome coalesce inside a bottleneck that occurred 300,000 years ago if the population size outside of the bottleneck is 10,000, and 19% of the genome coalesce inside of a million-year old bottleneck if the population size outside of it is 30,000 (Supplementary Table 20). To further verify that such types of bottleneck signatures can be detected with Relate from datasets comparable to our empirical ones, we performed additional simulations with msprime in which we applied a model that mimicked diversification of *Gadus morhua* populations as inferred from the supergene region on LG 2; this model included a bottleneck that affected the ancestor of one of two groups immediately after the divergence between them, and a population size reduction to a single sequence during this bottleneck. Demographic analysis of the simulated data with Relate in fact recovered the signature of the bottleneck, in the form of a slight but clear reduction in the estimated population size of members of the affected group that coincided with the age of the bottleneck (Supplementary Figure 11a). The ability of Relate to recover the bottleneck was not affected by an erroneous assumption of homogeneous recombination when data were simulated with a high-resolution recombination map (Supplementary Figure 11b).

**Supplementary Note 3:** Interpretation of signals of introgression between *Gadus morhua* and *Gadus ogac*.

We observed strong signals of introgression between *Gadus morhua* and *Gadus ogac*, affecting all sampled populations of *Gadus morhua* (Figs. 2c, 3c, 4b,e,h,k). Given that these signals were based on datasets mapped to the gadMor2 assembly, reference bias could potentially explain the observed introgression signals if the sequence data for *Gadus ogac* would have lower quality or shorter read lengths than that for *Gadus macrocephalus*[11], the sister species of *Gadus ogac*. This, however, appears unlikely, as the samples of both species were sequenced in the same study[1], with the same protocol, and to a similar read depth (17.8–27.5×). Nevertheless, to rule out reference bias as an explanation of these introgression signals, we repeated tests for introgression with an alternative dataset that was mapped to melAeg sequences from the threeway whole-genome alignment, representing *Melanogrammus aeglefinus*, a common outgroup to all taxa included in the dataset. Just like the original dataset mapped to gadMor2, this alternative dataset mapped to melAeg supported introgression between all *Gadus morhua* populations and *Gadus ogac* with $0.063 \leq D \leq 0.084$ ($p < 10^{-4}$). The absence of reference bias was further supported by a comparison of the numbers of reads with reference and alternate alleles at sites called as heterozygous in *Gadus ogac*, showing that sites with more reads carrying the reference allele are not more frequent than those with more reads carrying the alternate allele ($p > 0.05$), regardless of whether the dataset was mapped to gadMor2 or melAeg.

On the other hand, the fact that the introgression signals with *Gadus ogac* are largely homogenous across all sampled *Gadus morhua* populations, and with both the ancestral and derived arragments of the four supergenes, appears puzzling: Any gene flow with *Gadus ogac* could only have occurred after its divergence from *Gadus macrocephalus* (~40 ka), but because that divergence time is far younger than that of Northwest and Northeast Atlantic populations of *Gadus morhua* (~65 ka) or those between the supergene haplotypes (0.40–1.66 Ma), multiple separate but similarly strong gene flow events would be required from *Gadus ogac* into the different *Gadus morhua* populations and supergene haplotypes to explain the homogeneous signal. The signals of introgression with *Gadus ogac* are therefore best explained by gene flow in the opposite direction — from *Gadus morhua* into *Gadus ogac*. In this case, gene flow between one *Gadus morhua* population that co-occurs with *Gadus ogac* in the Northwest Atlantic would be sufficient to explain the observed signals for introgression, because it would increase allele sharing with *Gadus ogac* more or less evenly among all *Gadus morhua* populations, due to the relatedness among the *Gadus morhua* populations and supergene haplotypes. Under this assumption of gene flow from a Northwest Atlantic population into *Gadus ogac*, we expect a decreasing homogeneity of the introgression signal when it is quantified for more divergent supergene haplotypes, and this expectation is in fact met as the signal appears the least homogeneous for the supergene on LG 7 — the oldest of the supergenes — where it is markedly stronger for the haplotype that is carried by the Northwest Atlantic populations (Fig. 4h). The conclusion that *Gadus ogac* genomes carry introgression from *Gadus morhua* is also consistent with recent studies suggesting that introgression is more efficiently purged in species with larger population sizes[22–24], given that *Gadus ogac* appears to have low genetic diversity compared to *Gadus morhua*.

### Supplementary Figures

**Supplementary Figure 1:** Repeat content and mutational load in *Gadus morhua*.

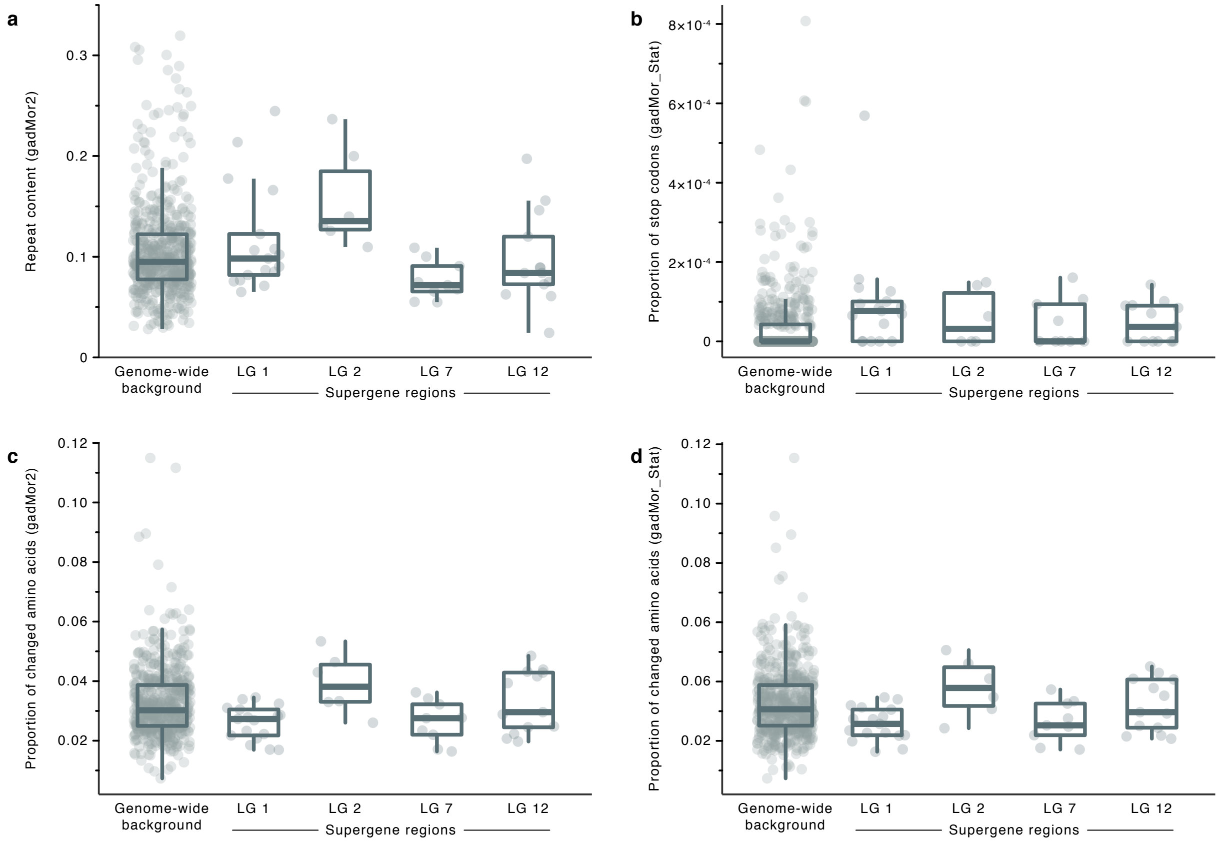


Repeat content and mutational load were quantified in sliding windows along the gadMor2 assembly[10]. Windows had a length or 1 Mbp and were grouped into supergene regions and the remainder of the genome ($n$ = 544, 17, 6, 9, and 13 for the genome-wide background and the four supergene regions, respectively). **a** Repeat content per window was quantified using the repeat annotation generated by Tørresen et al.[10] for the gadMor2 assembly. **b–d** Mutational load was calculated based on the three-way genome alignment, in which the gadMor_Stat and melAeg[25] assemblies were aligned to the gadMor2 assembly. As a first measure of mutational load, we quantified, per window, the proportion of stop codons among all codons in the gadMor_Stat sequences of the three-way alignment (**b**), according to gene annotation produced by Tørresen et al.[10] for the gadMor2 assembly. As a second measure of mutational load, we calculated the proportions of amino acids that were changed, compared to the melAeg assembly, in the gadMor2 (**c**) and gadMor_Stat assemblies (**d**), according to the three-way alignment. Per supergene region, we tested for increased repeat content or mutational load compared to the genome-wide background; however, no measures were significantly increased at false discovery rate (FDR) 0.05 (one-sided $t$-test; $p > 0.46$). Box plots show the median as center line, box sizes indicate the first and third quartiles, and whiskers extend to the most extreme values or 1.5× the interquartile range from the box limits.

**Supplementary Figure 2:** Concordance between reference genomes for *Gadus morhua*.

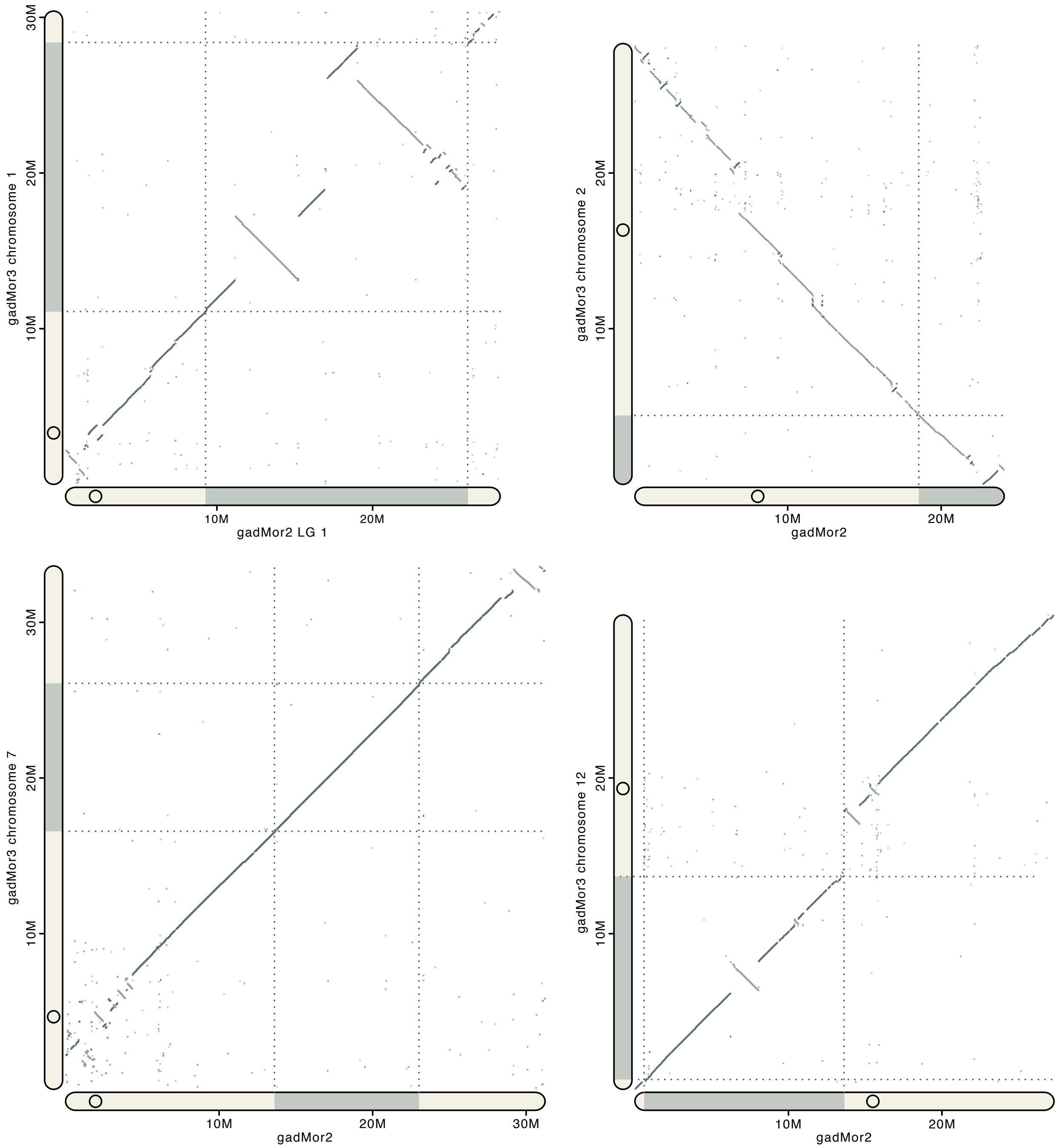


Dot plots illustrate regions of homology between the genome assemblies gadMor2[10] and gadMor3[26], for the four linkage groups with large genomic inversion regions in *Gadus morhua*. Gray shading and dotted lines indicate the beginning and end of supergene regions. Circles on scale bars indicate approximate centromere positions[26].

**Supplementary Figure 3:** Divergence times and introgression among *Gadus*, *Arctogadus*, and *Boreogadus*.

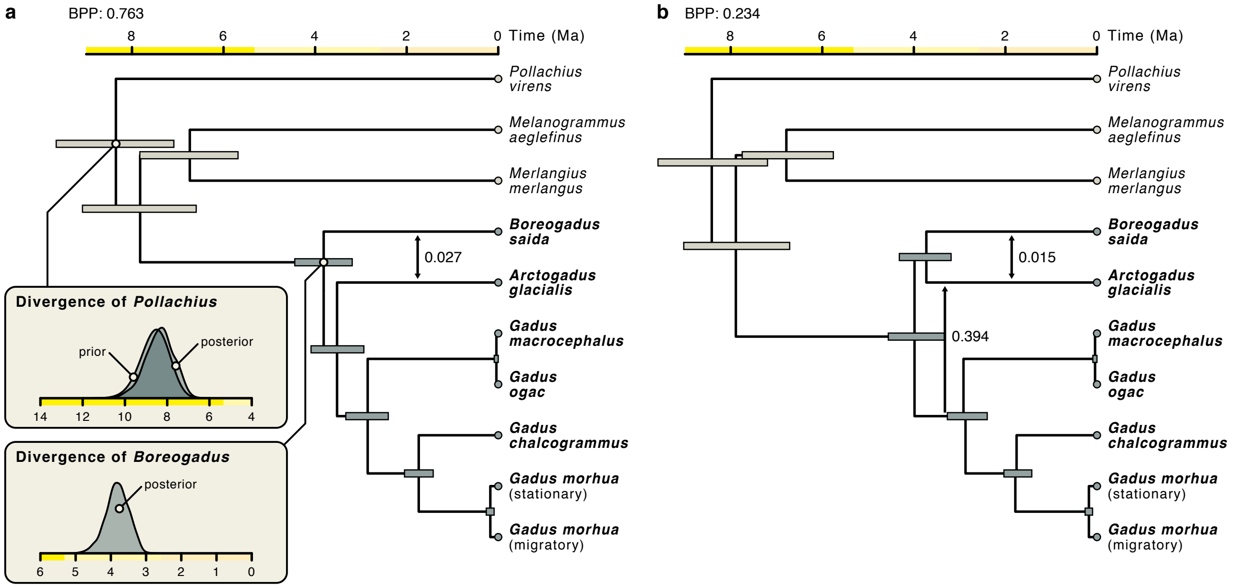


Maximum-clade-credibility species trees estimated with the AIM package for BEAST 2 under the isolation-with-migration model. Species of the three genera *Gadus*, *Arctogadus*, and *Boreogadus* are marked in bold; regular font is used for species considered as outgroups in this analysis. The tree was time-calibrated with a single prior distribution according to the results of our analysis of divergence times of Gadinae (Supplementary Figure 10), constraining the divergence of *Pollachius* to have occurred around 8.56 Ma (standard deviation: 0.08). Almost the entire posterior tree distribution (99.7%) supported one out of two constellations of topology and introgression. **a** Constellation of topology and introgression supported by a BPP of 0.763. The arrow between the branches leading to *Boreogadus saida* and *Arctogadus glacialis* indicates introgression supported with a Bayes factor greater than 10, the rate of which was estimated at 0.027 lineages per million years (95% HPD: 0–0.059). Insets show the prior distribution placed on the age of the divergence of *Pollachius virens*, the resulting posterior distribution for the same divergence event, and the posterior distribution for the divergence time of *Boreogadus saida*, which we used for time calibration in the downstream analyses of divergence times among *Gadus morhua* population. **b** Constellation of topology and introgression supported by a BPP of 0.234. Bayes factors greater than 10 support introgression between two pairs of lineages: from the common ancestor of the genus *Gadus* to *Arctogadus glacialis* with an estimated rate of 0.394 lineages per million years (95% HPD: 0.245–0.601) and between *Boreogadus saida* and *Arctogadus glacialis* with an estimated rate of 0.015 lineages per million years (95% HPD: 0–0.059). In both **a** and **b**, node bars indicate 95% HPD intervals for node ages.

**Supplementary Figure 4:** Distribution ranges of sampled species of the genera *Gadus*, *Arctogadus*, and *Boreogadus*.

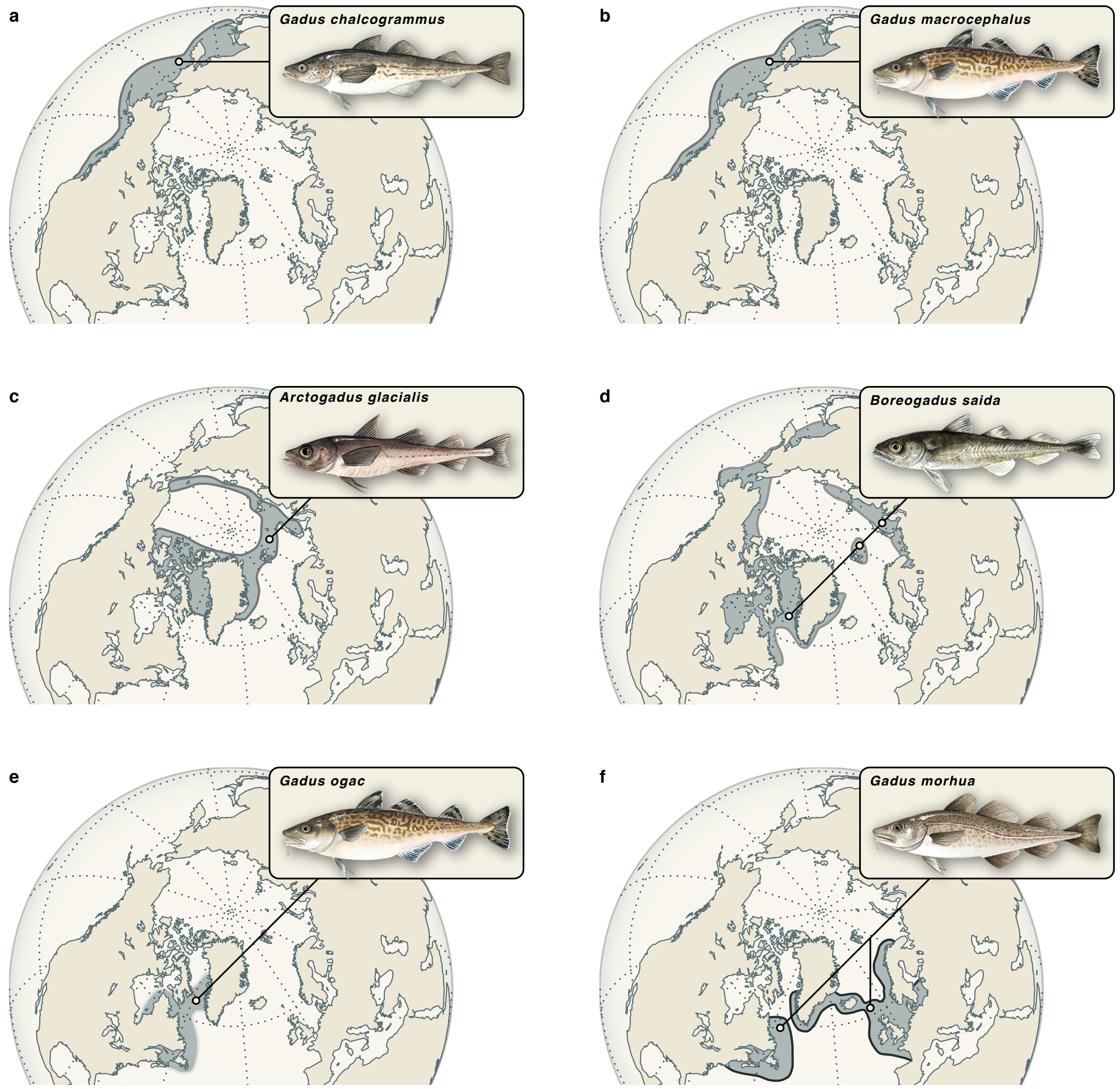


The distributions of the six species *G. chalcogrammus*, *G. macrocephalus*, *A. glacialis*, *B. saida*, *G. ogac*, and *G. morhua* are as shown in Fig. 2a but displayed separately.

**Supplementary Figure 5:** Tree-based signals for introgression among *Gadus*, *Arctogadus*, and *Boreogadus*.

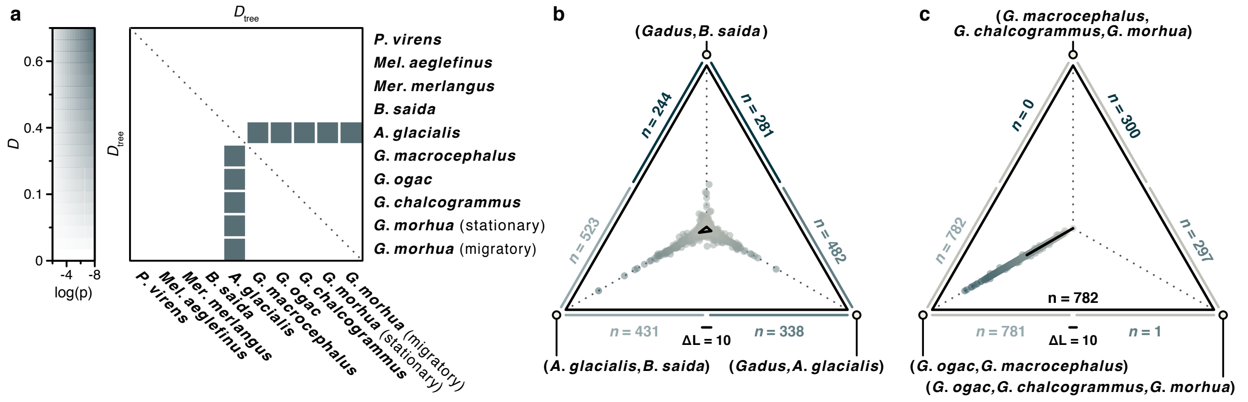


Signals of introgression among the genera *Gadus*, *Arctogadus*, and *Boreogadus*, obtained from sets of maximum-likelihood phylogenies. **a** The heatmap shows the tree-based equivalent to Patterson's $D$-statistic, $D_{\text{tree}}$[27], and its significance. Significant results ($p < 10^{-8}$) were only found in trios where *Boreogadus saida* and *Arctogadus glacialis* were placed as sister taxa, P1 and P2, and one of the species of the genus *Gadus* was in the position of the third species, P3. The grouping of P1 and P2 was then supported by 203 trees while 145 alignments supported the grouping of P2 and P3 and only 25 alignments supported the grouping of P1 and P3. The $D_{\text{tree}}$-statistic was thus the same for all these trios; $D_{\text{tree}} = (145 - 25)/(145 + 25) = 0.706$. **b** Likelihood support from alignments sampled across the genome for three alternative positions of *Arctogadus glacialis*: As the sister to *Boreogadus saida* (bottom left), as the sister to the genus *Gadus* (bottom right), and as an outgroup to a clade combining *Boreogadus saida* and the genus *Gadus* (top). Each dot in the triangle plot indicates the relative likelihood support that one alignment provides for each of the three hypotheses. Per dot, the distance from the center corresponds to the difference in likelihood between the best-supported topology and the alternative ones; the scale bar indicates the distance corresponding to a difference in log-likelihood support of 10. The small triangle in the center of the plot indicates the mean relative likelihood support for each topology, across all alignments. Sample sizes given on the edges of the triangle report the numbers of alignments that support one of the topologies connected by the edge over the other one. **c** As **b** but comparing the positions of *Gadus ogac* as the sister of *Gadus macrocephalus*, as the sister to *Gadus chalcogrammus* and *Gadus morhua*, and as the outgroup to a clade formed by *Gadus macrocephalus*, *Gadus chalcogrammus*, and *Gadus morhua*.

The tree-based tests for introgression corroborate the identification of *Arctogadus glacialis* as a species that introgressed either with *Boreogadus saida* or with the common ancestor of the genus *Gadus*, but do not further support the evidence coming from Patterson's $D$-statistic for introgression between *Gadus ogac* and *Gadus morhua*, perhaps because the introgressed regions were in this case too short to affect tree topologies.

**Supplementary Figure 6:** Divergence times among *Gadus morhua* populations.

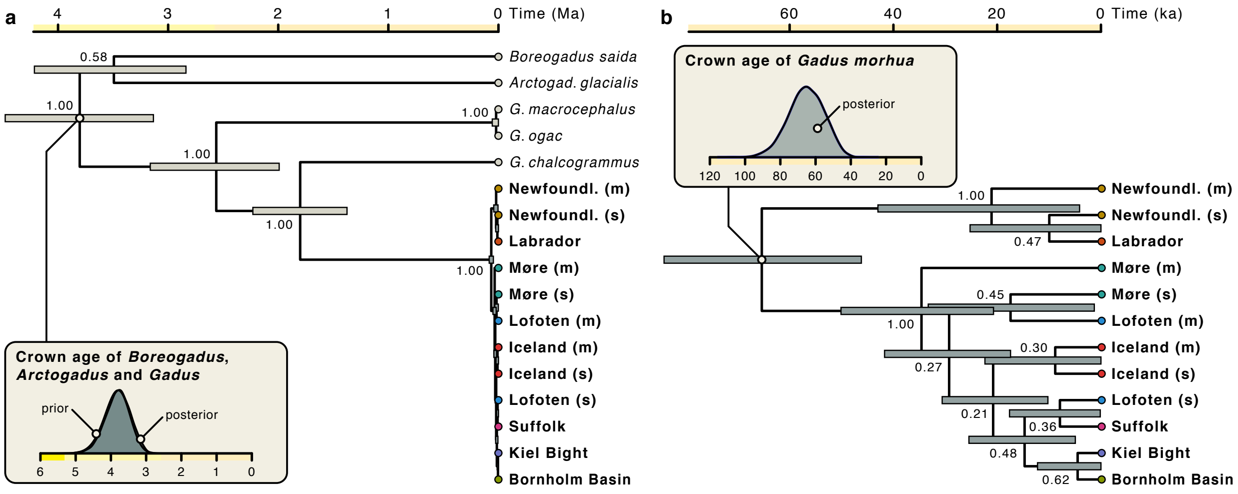


**a** Maximum-clade-credibility tree of *Gadus morhua* populations and outgroups inferred from genome-wide SNPs while accounting for incomplete lineage sorting with SNAPP. Single-nucleotide polymorphisms from within the four supergenes were excluded in the inference. *Gadus morhua* populations are marked in bold; regular font is used for species here considered as outgroups. The tree was time-calibrated with a single prior distribution to constrain the crown age of *Gadus*, *Arctogadus*, and *Boreogadus* to around 3.83 Ma (standard deviation: 0.093), according to the results of our analysis of divergence times among the three genera with the isolation-with-migration model (Fig. 2a; Supplementary Figure 3). The prior distribution is shown in the inset together with the posterior distribution for the same divergence event; these two distributions are nearly identical. **b** Enlarged section of the maximum-clade-credibility tree showing divergences among *Gadus morhua* populations in more detail. The posterior distribution for the crown age of *Gadus morhua* is shown in the inset. In both **a** and **b**, numbers next to nodes report BPP support values and node bars indicate 95% HPD intervals for node ages. This tree is identical to the maximum-clade-credibility tree plotted in Figure 3.

**Supplementary Figure 7:** Divergence times among *Gadus morhua* populations, specific to supergenes.

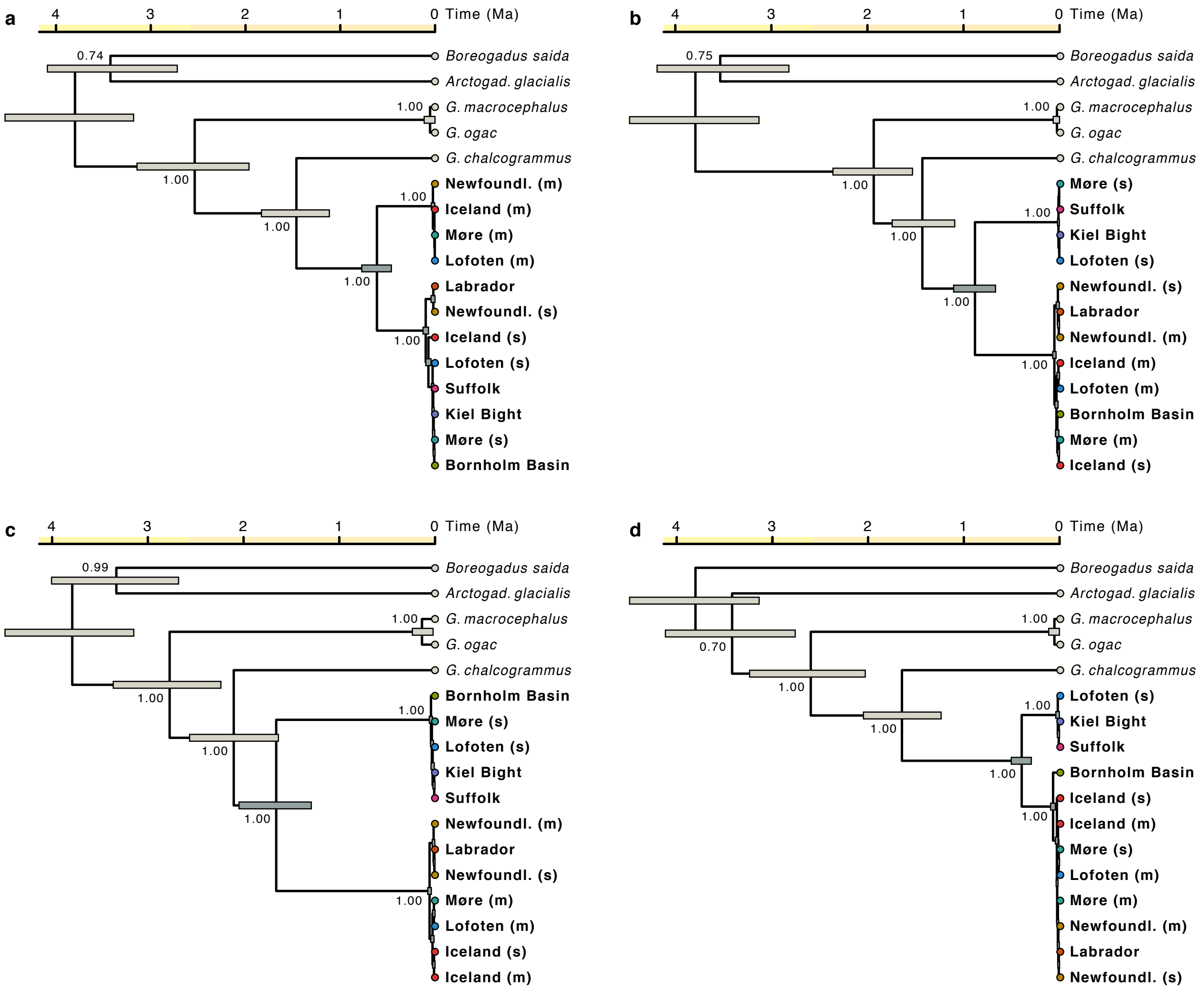


Maximum-clade-credibility trees of *Gadus morhua* populations and outgroups, separately inferred with SNAPP for each of the four supergenes. Inference settings were identical to those used to infer the MCC tree shown in Supplementary Figure 6a, except that only SNPs from supergene regions were used. **a–d** Maximum-clade-credibility trees for the supergene on linkage groups (LGs) 1 (**a**), 2 (**b**), 7 (**c**), and 12 (**d**). Numbers next to nodes report BPP support values and node bars indicate 95% HPD intervals for node ages. These trees are identical to the ones shown in Figures 4a, 5a, 5e, and 6a.

**Supplementary Figure 8:** Between-population divergence along linkage groups.

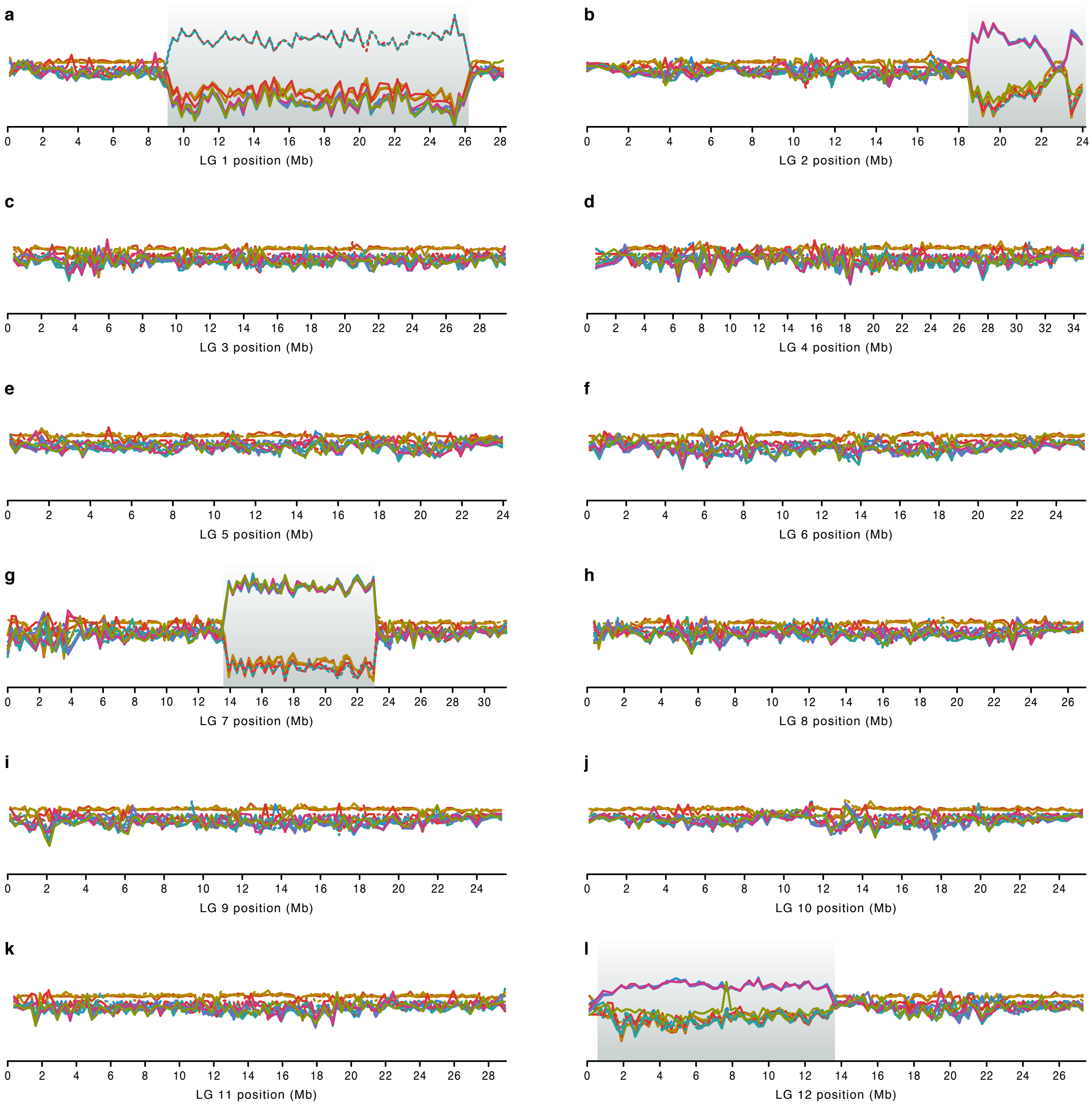


Divergence times were estimated with SNAPP from SNPs in sliding windows. **a**–**l** Between-population mean divergence-time estimates along LGs 1 (**a**), 2 (**b**), 3 (**c**), 4 (**d**), 5 (**e**), 6 (**f**), 7 (**g**), 8 (**h**), 9 (**i**), 10 (**j**), 11 (k), and 12 (**l**), plotted as in Figures 4d, 5d,h and 6d. The four supergene regions on LGs 1, 2, 7, and 12 are indicated with gray background.

**Supplementary Figure 8 (continued):** Between-population divergence along linkage groups.

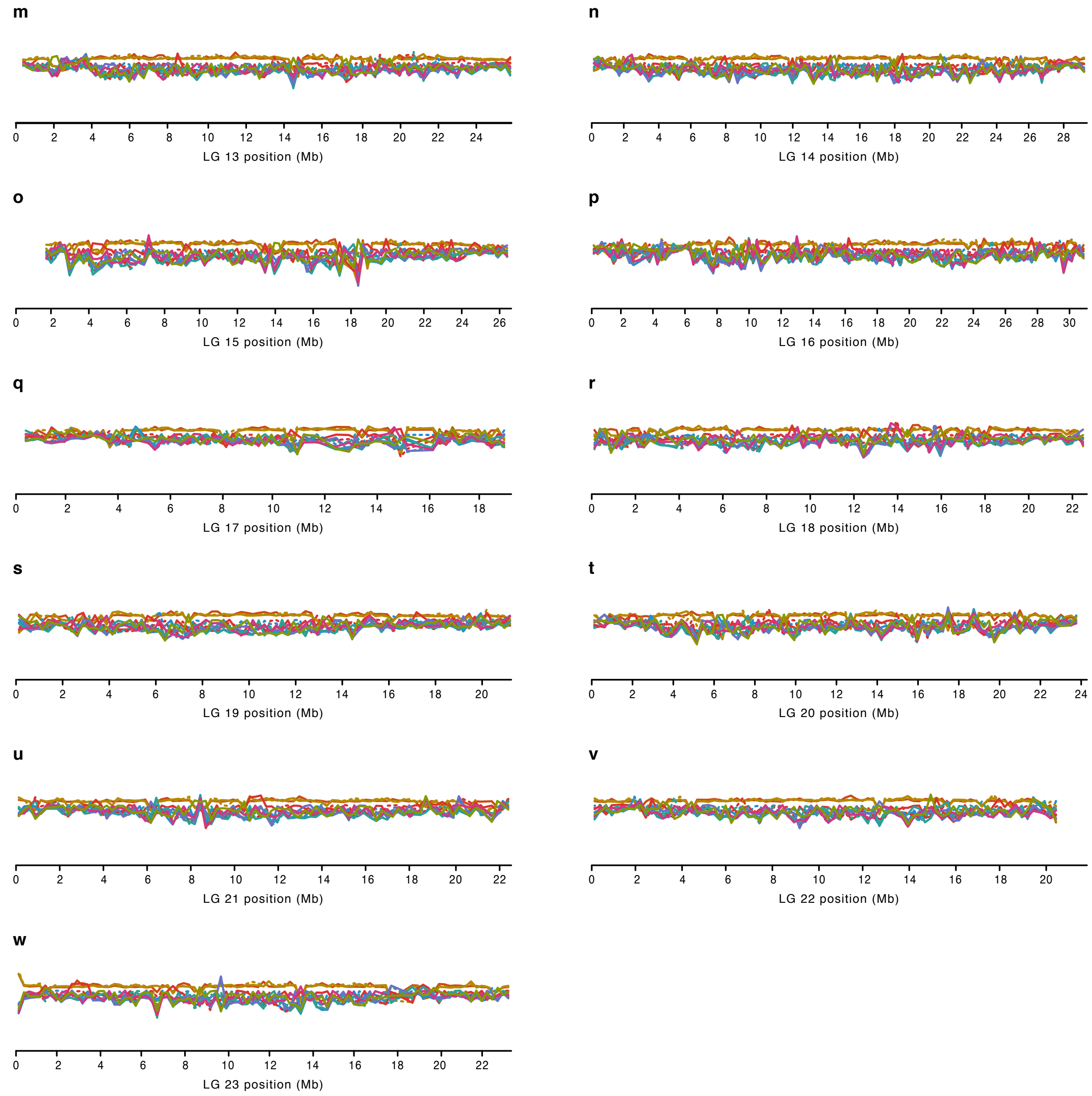


**m–w** Between-population divergence times along LGs 13 (**m**), 14 (**n**), 15 (**o**), 16 (**p**), 17 (**q**), 18 (**r**), 19 (**s**), 20 (**t**), 21 (**u**), 22 (**v**), and 23 (**w**).

**Supplementary Figure 9:** Measures of differentiation and divergence for linkage groups with supergenes.

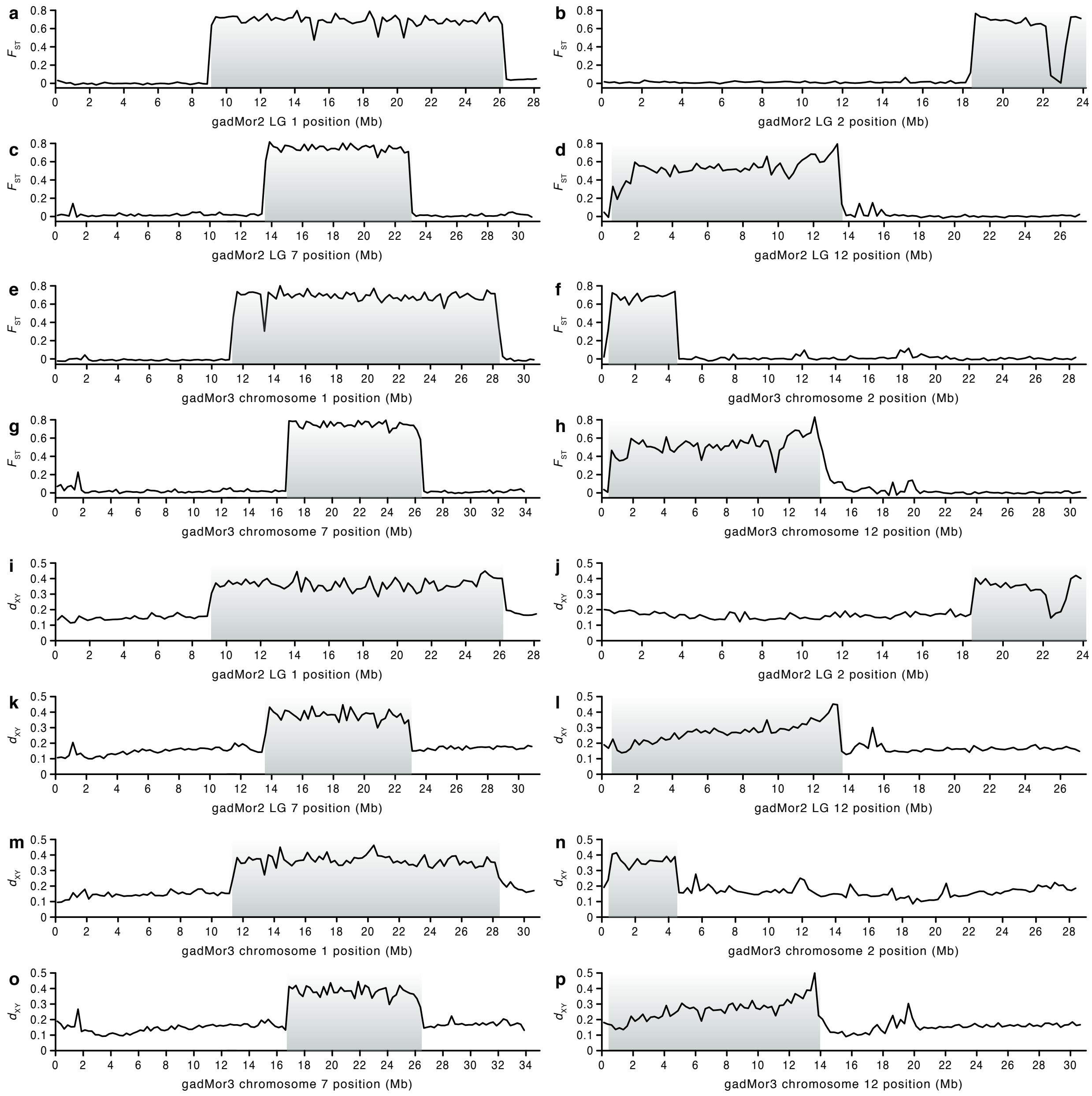


As a complement to the patterns of temporal divergence shown in Fig. 5, differentiation and divergence across linkage groups with supergenes were also quantified as $F_{\mathrm{st}}$ (**a-h**) and $d_{\mathrm{xy}}$ (**i-p**). As in Fig. 5, the two measures were calculated in sliding windows with a length of 250 kbp, for predefined groups of populations that separated those with the ancestral and derived supergene orientations. Both measures are plotted across linkage groups of the gadMor2 assembly (**a-d**, **i-l**) and chromosomes of the newer gadMor3[26] (**e-h**, **m-p**) assembly. Note that gadMor3 chromosome 2 is inverted relative to gadMor2 LG 2 (see Supplementary Figure 2). Comparable results were obtained by Barth et al.[28] for gadMor2 LGs 2, 7, and 12, using a different dataset and shorter window sizes of 50 and 100 kbp.

**Supplementary Figure 10:** Divergence times of Gadinae.

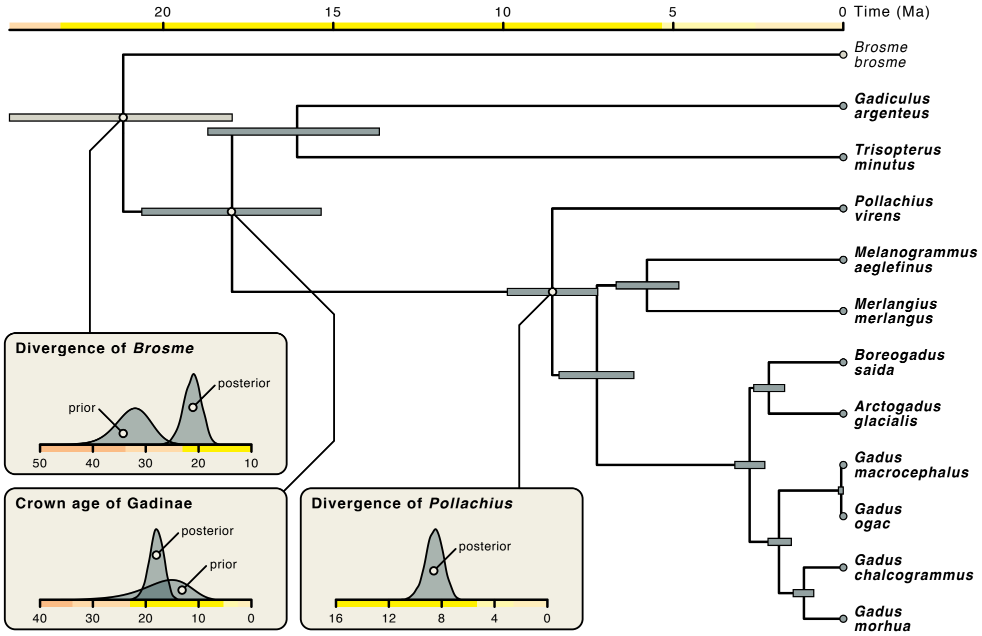


Maximum-clade-credibility species tree of Gadinae estimated with StarBEAST from 91 gene alignments under the multi-species coalescent model. Species included in the subfamily Gadinae are marked in bold. The tree was time-calibrated with two prior distributions according to Musilova et al.[29], constraining the divergence of the outgroup *Brosme brosme* to have occurred around 32.325 Ma (standard deviation: 0.1), and the first divergence within crown Gadinae to have occurred around 18.1358 Ma (standard deviation: 0.28). These prior distributions are shown in insets together with the respective posterior distributions; also shown is the posterior distribution for the divergence time of *Pollachius virens*, which we used for time calibration of the downstream estimation of divergence times among *Gadus*, *Arctogadus*, and *Boreogadus* with the isolation-with-migration model (Fig. 2; Supplementary Figure 3). Node bars indicate 95% highest-posterior-density (HPD) intervals for node ages. All nodes were supported by Bayesian posterior probabilities (BPP) of 1.0.

**Supplementary Figure 11:** Population sizes over time estimated with Relate from simulated data.

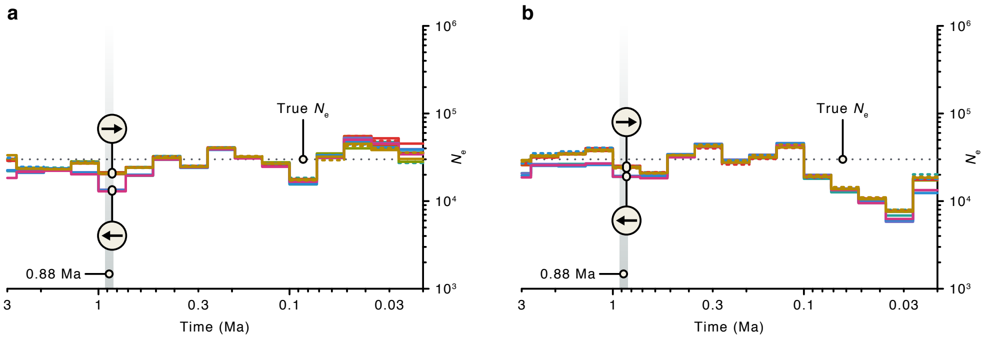


To test the ability of Relate to recover bottleneck events associated with old genomic inversions, we simulated chromosome-scale genetic data under a model of population diversification with msprime v.0.7.4[21], and inferred population sizes over time from these data with Relate v.1.1.2[30]. The selected model mimicked diversification of *Gadus morhua* populations as inferred from the supergene on LG 2 (Fig. 5a), with an initial divergence separating two haplotypes (labelled with forward and reverse arrows) 880,823 years ago, immediately followed by an extreme bottleneck in one of the two descending lineages during which the population size was reduced to a single sequence. The length of the bottleneck was set to ten generations. Outside of the bottleneck, all population sizes were set to a diploid size of $N_{\mathrm{e}} = 30,000$. The generation time was set to 10 years and mutations were added with a rate identical to the one inferred for our empirical dataset ($\mu = 1.64 \times 10^{-9}$ per bp and year). To model recombination rate variation in realistic detail we used a recombination map for a human chromosome from the HapMap project[31] because we assumed that the resolution of these maps would exceed those of available maps for teleost fishes. We selected the recombination map for human chromosome 20, which did not contain large regions without information and was more comparable in size to fish chromosomes than other maps from the HapMap project. **a,b** The inference of population sizes over time with Relate used the same settings as our inference from empirical data (Fig. 4f), except that, to allow an assessment of the effect of unknown recombination rate variation, we inferred population sizes from the simulated data once with the true recombination map as input to Relate (**a**) and once assuming homogeneous recombination across the chromosome (**b**). The dotted line shows the true population size used in simulations and the gray region indicates the time of the simulated bottleneck (the width of this region is not drawn to scale). Regardless of whether the true recombination map was used or homogenous recombination was assumed, the signature of the bottleneck was recognizable as a separation of the estimated population sizes per haplotype around the time of the bottleneck, with temporarily decreased estimates for the populations carrying the affected haplotype (marked with "i"). The true depth of the simulated bottleneck (an $N_{\mathrm{e}}$ corresponding to a single sequence), however, was not recovered, probably because Relate estimates population sizes in time bins that are much wider than the duration of the bottleneck, spanning up to several hundred thousand years.

In most time bins outside the bottleneck, the inferred population size was close to the true $N_\mathrm{e}$ used in the simulation.

### Supplementary Tables

**Supplementary Table 1:** Assembly completeness.

Assembly completeness was assessed with the program BUSCO v.5.0, using the Actinopterygii set of 3,640 conserved and single-copy orthologs.

| Category | Frequency | Percent of total |
|---|---|---|
| Complete BUSCO genes | 3,061 | 84.1 |
| Complete and single-copy BUSCO genes | 3,020 | 83.0 |
| Complete and duplicated BUSCO genes | 41 | 1.1 |
| Fragmented BUSCO genes | 218 | 6.0 |
| Missing BUSCO genes | 361 | 9.9 |

**Supplementary Table 2:** Genetic distances between gadiform genome assemblies.
Genetic distances were calculated from the three-way whole-genome alignment in which the gadMor_Stat and melAeg assemblies were aligned to the gadMor2 assembly. The three-way alignment was carefully filtered for reliably aligned sites, and pairwise genetic distances were calculated as the proportion of sites considered reliably aligned that differed between a pair of assemblies. We report, per gadMor2 linkage group, the genetic distances between the gadMor_Stat and gadMor2 assemblies ($d_{\text{gadMor\_Stat}-\text{gadMor2}}$), the genetic distance between the melAeg and gadMor2 assemblies ($d_{\text{melAeg}-\text{gadMor2}}$), and the relative difference between the gadMor_Stat and gadMor2 assemblies, compared to the difference between the melAeg and gadMor2 assemblies ($d_{\text{gadMor\_Stat}-\text{gadMor2}}$ / $d_{\text{melAeg}-\text{gadMor2}}$).

| LG | $d_{\text{gadMor\_Stat}-\text{gadMor2}}$ | $d_{\text{melAeg}-\text{gadMor2}}$ | $d_{\text{gadMor\_Stat}-\text{gadMor2}}/d_{\text{melAeg}-\text{gadMor2}}$ |
|---|---|---|---|
| 1 | 0.00833 | 0.04675 | 0.17816 |
| 2 | 0.00615 | 0.05136 | 0.11980 |
| 3 | 0.00454 | 0.04696 | 0.09659 |
| 4 | 0.00534 | 0.05212 | 0.10251 |
| 5 | 0.00459 | 0.04726 | 0.09701 |
| 6 | 0.00411 | 0.04564 | 0.08994 |
| 7 | 0.00681 | 0.04756 | 0.14327 |
| 8 | 0.00471 | 0.04842 | 0.09723 |
| 9 | 0.00407 | 0.04752 | 0.08560 |
| 10 | 0.00443 | 0.05032 | 0.08800 |
| 11 | 0.00423 | 0.04523 | 0.09362 |
| 12 | 0.00581 | 0.04941 | 0.11756 |
| 13 | 0.00401 | 0.04782 | 0.08386 |
| 14 | 0.00404 | 0.04637 | 0.08720 |
| 15 | 0.00421 | 0.04504 | 0.09356 |
| 16 | 0.00419 | 0.04636 | 0.09040 |
| 17 | 0.00518 | 0.05474 | 0.09462 |
| 18 | 0.00436 | 0.04926 | 0.08849 |
| 19 | 0.00450 | 0.05136 | 0.08759 |
| 20 | 0.00435 | 0.04563 | 0.09539 |
| 21 | 0.00449 | 0.05026 | 0.08942 |
| 22 | 0.00486 | 0.04998 | 0.09714 |
| 23 | 0.00466 | 0.04954 | 0.09411 |

| LG | Region | $d_{\text{gadMor\_Stat}-\text{gadMor2}}$ | $d_{\text{melAeg}-\text{gadMor2}}$ | $d_{\text{gadMor\_Stat}-\text{gadMor2}}/d_{\text{melAeg}-\text{gadMor2}}$ |
|---|---|---|---|---|
| 1 | 9,114,741–26,192,489 | 0.01069 | 0.04504 | 0.23736 |
| 2 | 18,489,307–24,050,282 | 0.01285 | 0.05805 | 0.22141 |
| 7 | 13,606,502–23,016,726 | 0.00909 | 0.04257 | 0.21348 |
| 12 | 589,105–13,631,347 | 0.00658 | 0.04868 | 0.13523 |

**Supplementary Table 3:** Contigs of the gadMor_Stat and melAeg assemblies aligning near supergene boundaries.
Contigs were aligned to the gadMor2 assembly[10] with BLASTN searches. Per alignment, we report the start and end positions of aligning regions on both the gadmor2 assembly (in the form "LG:start position–end position") and on the contigs from the gadMor_Stat and melAeg[25] assemblies. Contig labels correspond to those used in Figure 1b.

| Label | Assembly | Contig | gadMor2 position | Contig position | Orientation |
|---|---|---|---|---|---|
| 1 | gadMor_Stat | scf7180000174049 | 1:8,943,539–9,128,372 | 1–200,341 | forward |
| | | | 1:16,848,173–16,296,669 | 200,341–707,031 | reverse |
| | | | 1:26,322,778–26,326,857 | 706,985–710,054 | forward |
| 2 | gadMor_Stat | scf7180000173314 | 1:9,130,274–9,192,260 | 253,972–314,079 | forward |
| | | | 1:18,622,815–18,890,477 | 254–258,023 | forward |
| 3 | melAeg | scf1854:1-47276 | 1:9,090,044–9,120,848 | 4–47,275 | forward |
| 4 | melAeg | scg895:1-207496 | 1:9,130,297–9,187,321 | 1–59,877 | reverse |
| | | | 1:18,731,791–18,890,477 | 56,307–205,627 | reverse |
| 5 | gadMor_Stat | scf7180000174960 | 1:26,199,094–26,219,392 | 22–28,398 | forward |
| 6 | gadMor_Stat | scf7180000173351 | 2:18,399,575–18,494,225 | 1–89,835 | forward |
| 7 | gadMor_Stat | scf7180000176322 | 2:18,567,231–18,487,151 | 6–75,867 | reverse |
| | | | 2:23,150,464–23,200,546 | 68,286–108,674 | reverse |
| 8 | melAeg | scf170:27711-284173 | 2:18,360,264–18,642,641 | 1–255901 | forward |
| 9 | gadMor_Stat | scf7180000175069 | 2:23,939,789–24,054,399 | 4–107,435 | forward |
| 10 | melAeg | scf874:12556-53982 | 2:24,007,915–24,049,209 | 1–39,552 | reverse |
| 11 | gadMor_Stat | scf7180000173083 | 7:13,241,739–13,642,122 | 1–403,620 | reverse |
| 12 | gadMor_Stat | scf7180000173082 | 7:13,645,465–13651003 | 30,270–37,901 | reverse |
| | | | 7:22,971,316–23,002,424 | 4–28,473 | forward |
| 13 | gadMor_Stat | scf7180000177071 | 7:13,652,432–13,689,724 | 33–23,593 | reverse |
| | | | 7:23,269,226–23,270,717 | 21,916–23,360 | forward |
| 14 | melAeg | scf271:1-285897 | 7:13,585,625–13,651,514 | 6–66,452 | forward |
| | | | 7:22,773,521–23,002,439 | 68,044–285,897 | reverse |
| 15 | gadMor_Stat | scf7180000175147 | 7:23,043,967–23,074,768 | 1–31,361 | forward |
| 16 | gadMor_Stat | scf7180000176674 | 12:579,198–607,782 | 20,228–51,625 | reverse |
| | | | 12:13,378,418–13,386,293 | 1–10,649 | forward |
| 17 | gadMor_Stat | scf7180000174053 | 12:662,878–694,568 | 8–27,278 | forward |
| | | | 12:13,614,908–13,616,907 | 23,719–25,709 | reverse |

**Supplementary Table 4:** Sequence data used in analyses of divergence times of Gadinae. Illumina reads for *Gadus macrocephalus* and *Gadus ogac* were used for local assembly of candidate phylogenetic markers, all local and whole-genome assemblies were used to identify sets of orthologuous sequences for 91 genes, and alignments of these sequences were used to estimate divergence times with StarBEAST2[32, 33] (Supplementary Figure 10).

| Species | Data type | Reference | Accession |
|---|---|---|---|
| *Brosme brosme* | Genome assemby | Malmstrøm et al.[8] | Dryad: 326r8 |
| *Gadiculus argenteus* | Genome assemby | Malmstrøm et al.[8] | Dryad: 326r8 |
| *Trisopterus minutus* | Genome assemby | Malmstrøm et al.[8] | Dryad: 326r8 |
| *Pollachius virens* | Genome assemby | Malmstrøm et al.[8] | Dryad: 326r8 |
| *Melanogrammus aeglefinus* | Genome assemby | Malmstrøm et al.[8] | Dryad: 326r8 |
| *Merlangius merlangus* | Genome assemby | Malmstrøm et al.[8] | Dryad: 326r8 |
| *Boreogadus saida* | Genome assemby | Malmstrøm et al.[8] | Dryad: 326r8 |
| *Arctogadus glacialis* | Genome assemby | Malmstrøm et al.[8] | Dryad: 326r8 |
| *Gadus macrocephalus* | Illumina reads | Árnason and Halldórsdóttir[1] | ENA: SRR2906345 |
| *Gadus ogac* | Illumina reads | Árnason and Halldórsdóttir[1] | ENA: SRR2906193 |
| *Gadus chalcogrammus* | Genome assemby | Malmstrøm et al.[8] | Dryad: 326r8 |
| *Gadus morhua* | Genome assemby[1] | Tørresen et al.[10] | ENA: ERR1551885 |

[1]gadMor2 assembly[10]

**Supplementary Table 5:** Read data used in analyses of divergence times and introgression among species of the genera *Gadus*, *Arctogadus*, and *Boreogadus*.
Illumina reads were mapped to the gadMor2 assembly[10] and used to generate alignments for 109 phylogenetic markers, which were then analyzed with the isolation-with-migration model implemented in the AIM package[34] for BEAST 2[32].

| Species | Reference | ENA accession(s) |
|---|---|---|
| *Pollachius virens* | Malmstrøm et al.[8] | ERR1473878 |
| *Melanogrammus aeglefinus* | Tørresen et al.[25] | ERR2028455,ERR2028456 |
| *Merlangius merlangus* | Malmstrøm et al.[8] | ERR1473880,ERR1473881 |
| *Boreogadus saida* | Malmstrøm et al.[8] | ERR1473884,ERR1473885 |
| *Arctogadus glacialis* | Malmstrøm et al.[8] | ERR1473882,ERR1473883 |
| *Gadus macrocephalus* | Árnason and Halldórsdóttir[1] | SRR2906345 |
| *Gadus ogac* | Árnason and Halldórsdóttir[1] | SRR2906193 |
| *Gadus chalcogrammus* | Malmstrøm et al.[8] | ERR1473886 |
| *Gadus morhua* (migratory) | This study | ERR5330167[1] |
| *Gadus morhua* (stationary) | This study | ERR5321904[2] |

[1]Migratory *Gadus morhua* individual LOF1103Z24 sampled at the Lofoten islands, with a genome similar to the gadMor2 genome (2,940 differences excluding heterozygous sites)
[2]Stationary *Gadus morhua* individual LOF1106Z11 used for the gadMor_Stat assembly

**Supplementary Table 6:** $D_{\mathrm{BBAA}}$-statistics for species of the genera *Gadus*, *Arctogadus*, and *Boreogadus* and outgroups.

For the calculation of the $D_{\mathrm{BBAA}}$ version of the $D$-statistic[35], all trios are oriented so that the number of "BBAA" patterns ($C_{\mathrm{BBAA}}$) is greater than the numbers of "ABBA" ($C_{\mathrm{ABBA}}$) and "BBAA" ($C_{\mathrm{BABA}}$) patterns. Additionally, the trio is arbitrarily oriented so that the number of "ABBA" ($C_{\mathrm{ABBA}}$) patterns exceeds that of "BBAA" ($C_{\mathrm{BABA}}$) patterns. When the trio is oriented in this way, the $D_{\mathrm{BBAA}}$-statistic is never negative and values significantly greater than zero support introgression between the two species in positions P2 and P3. If the same pair of species in positions P2 and P3 produced significant results with several third species in position P1, only the trio with the highest $D_{\mathrm{BBAA}}$ is listed. Significance was assessed with block-jackknifing. The highest $D$ values are underlined.

| P1 | P2 | P3 | $C_{\mathrm{BBAA}}$ | $C_{\mathrm{ABBA}}$ | $C_{\mathrm{BABA}}$ | $D_{\mathrm{BBAA}}$ | $p$ |
|---|---|---|---|---|---|---|---|
| *G. morhua* (m)[1] | *B. saida* | *Mel. aeglefinus* | 999,222.0 | 74,770.5 | 70,196.2 | 0.032 | 0.0000 |
| *G. morhua* (m) | *A. glacialis* | *Mel. aeglefinus* | 1043,480.0 | 65,118.6 | 63,258.9 | 0.014 | 0.0000 |
| *G. morhua* (m) | *G. macrocephal.* | *Mel. aeglefinus* | 1,381,730.0 | 48,034.4 | 46,407.9 | 0.017 | 0.0000 |
| *G. morhua* (m) | *G. ogac* | *Mel. aeglefinus* | 1,378,670.0 | 47,735.4 | 45,982.6 | 0.019 | 0.0000 |
| *G. morhua* (m) | *G. chalcogram.* | *Mel. aeglefinus* | 1,630,450.0 | 32,523.8 | 31,345.2 | 0.018 | 0.0000 |
| *G. morhua* (m) | *G. morhua* (s)[2] | *Mel. aeglefinus* | 2,128,480.0 | 6,106.4 | 5,964.9 | 0.012 | 0.0360 |
| *Mel. aeglefinus* | *Mer. merlangus* | *B. saida* | 524,569.0 | 109,421.0 | 104,705.0 | 0.022 | 0.0000 |
| *Mel. aeglefinus* | *Mer. merlangus* | *A. glacialis* | 526,258.0 | 103,925.0 | 99,293.4 | 0.023 | 0.0000 |
| *Mel. aeglefinus* | *Mer. merlangus* | *G. macrocephal.* | 535,103.0 | 106,401.0 | 101,740.0 | 0.022 | 0.0000 |
| *Mel. aeglefinus* | *Mer. merlangus* | *G. ogac* | 532,068.0 | 105,833.0 | 101,330.0 | 0.022 | 0.0000 |
| *Mel. aeglefinus* | *Mer. merlangus* | *G. chalcogram.* | 534,655.0 | 107,694.0 | 101,792.0 | 0.028 | 0.0000 |
| *Mel. aeglefinus* | *Mer. merlangus* | *G. morhua* (s) | 544,694.0 | 110,805.0 | 103,491.0 | 0.034 | 0.0000 |
| *Mel. aeglefinus* | *Mer. merlangus* | *G. morhua* (m) | 544,885.0 | 110,788.0 | 103,421.0 | 0.034 | 0.0000 |
| *G. morhua* (m) | *G. macrocephal.* | *B. saida* | 461,022.0 | 87,150.0 | 74,312.0 | 0.080 | 0.0000 |
| *G. morhua* (m) | *G. ogac* | *B. saida* | 463,606.0 | 86,146.6 | 73,616.9 | 0.078 | 0.0000 |
| *G. morhua* (m) | *G. chalcogram.* | *B. saida* | 699,019.0 | 51,353.8 | 42,041.8 | 0.100 | 0.0000 |
| *G. morhua* (m) | *G. morhua* (s) | *B. saida* | 1,178,120.0 | 6,965.9 | 6,699.1 | 0.020 | 0.0004 |
| *B. saida* | *A. glacialis* | *G. macrocephal.* | 267,280.0 | 170,777.0 | 132,905.0 | 0.125 | 0.0000 |
| *B. saida* | *A. glacialis* | *G. ogac* | 265,999.0 | 169,897.0 | 132,210.0 | 0.125 | 0.0000 |
| *B. saida* | *A. glacialis* | *G. chalcogram.* | 268,779.0 | 170,122.0 | 131,604.0 | 0.128 | 0.0000 |
| *B. saida* | *A. glacialis* | *G. morhua* (s) | 279,966.0 | 170,630.0 | 131,786.0 | 0.128 | 0.0000 |
| *B. saida* | *A. glacialis* | *G. morhua* (m) | 280,258.0 | 170,613.0 | 131,776.0 | 0.128 | 0.0000 |
| *G. morhua* (m) | *G. chalcogram.* | *G. macrocephal.* | 337,874.0 | 74,069.5 | 55,581.5 | 0.143 | 0.0000 |
| *G. morhua* (m) | *G. morhua* (s) | *G. macrocephal.* | 820,109.0 | 7,930.4 | 7,495.9 | 0.028 | 0.0000 |
| *G. macrocephal.* | *G. ogac* | *G. chalcogram.* | 941,162.0 | 9,380.4 | 7,456.4 | 0.114 | 0.0000 |
| *G. macrocephal.* | *G. ogac* | *G. morhua* (s) | 980,086.0 | 13,956.1 | 7,797.4 | <u>0.283</u> | 0.0000 |
| *G. macrocephal.* | *G. ogac* | *G. morhua* (m) | 980,709.0 | 14,167.9 | 7,875.9 | <u>0.285</u> | 0.0000 |
| *G. morhua* (m) | *G. morhua* (s) | *G. chalcogram.* | 532,010.0 | 10,099.0 | 9,619.8 | 0.024 | 0.0000 |

[1]Migratory *Gadus morhua* individual LOF1103Z24 sampled at the Lofoten islands

[2]Stationary *Gadus morhua* individual LOF1106Z11 sampled at the Lofoten islands

**Supplementary Table 7:** $D_{\text{fix}}$-statistics for species of the genera *Gadus*, *Arctogadus*, and *Boreogadus* and outgroups.

For the calculation of the $D_{\text{fix}}$ version of the $D$-statistic[35], all trios are fixed in their orientation according to an assumed species tree, for which we here use the more strongly supported one out of two trees resulting from the inference under the isolation-with-migration model (Fig. 2b). Additionally, the trio is arbitrarily oriented so that the number of "ABBA" ($C_{\text{ABBA}}$) patterns exceeds that of "BBAA" ($C_{\text{BABA}}$) patterns. When the trio is oriented in this way, the $D_{\text{BBAA}}$-statistic is never negative and values significantly greater than zero support introgression between the two species in positions P2 and P3. If the same pair of species in positions P2 and P3 produced significant results with several third species in position P1, only the trio with the highest $D_{\text{BBAA}}$ is listed. Significance was assessed with block-jackknifing. The highest $D$ values are underlined.

| P1 | P2 | P3 | $C_{\text{BBAA}}$ | $C_{\text{ABBA}}$ | $C_{\text{BABA}}$ | $D_{\text{fix}}$ | $p$ |
|---|---|---|---:|---:|---:|---:|---:|
| *G. morhua* (m)[1] | *B. saida* | *Mel. aeglefinus* | 999,222.0 | 74,770.5 | 70,196.2 | 0.032 | 0.0000 |
| *G. morhua* (m) | *A. glacialis* | *Mel. aeglefinus* | 1,043,480.0 | 65,118.6 | 63,258.9 | 0.014 | 0.0000 |
| *G. morhua* (m) | *G. macrocephalus* | *Mel. aeglefinus* | 1,381,730.0 | 48,034.4 | 46,407.9 | 0.017 | 0.0000 |
| *G. morhua* (m) | *G. ogac* | *Mel. aeglefinus* | 1,378,670.0 | 47,735.4 | 45,982.6 | 0.019 | 0.0000 |
| *G. morhua* (m) | *G. chalcogrammus* | *Mel. aeglefinus* | 1,630,450.0 | 32,523.8 | 31,345.2 | 0.018 | 0.0000 |
| *G. morhua* (m) | *G. morhua* (s)[2] | *Mel. aeglefinus* | 2,128,480.0 | 6,106.4 | 5,964.9 | 0.012 | 0.0360 |
| *Mel. aeglefinus* | *Mer. merlangus* | *B. saida* | 524,569.0 | 109,421.0 | 104,705.0 | 0.022 | 0.0000 |
| *Mel. aeglefinus* | *Mer. merlangus* | *A. glacialis* | 526,258.0 | 103,925.0 | 99,293.4 | 0.023 | 0.0000 |
| *Mel. aeglefinus* | *Mer. merlangus* | *G. macrocephalus* | 535,103.0 | 106,401.0 | 101,740.0 | 0.022 | 0.0000 |
| *Mel. aeglefinus* | *Mer. merlangus* | *G. ogac* | 532,068.0 | 105,833.0 | 101,330.0 | 0.022 | 0.0000 |
| *Mel. aeglefinus* | *Mer. merlangus* | *G. chalcogrammus* | 534,655.0 | 107,694.0 | 101,792.0 | 0.028 | 0.0000 |
| *Mel. aeglefinus* | *Mer. merlangus* | *G. morhua* (s) | 544,694.0 | 110,805.0 | 103,491.0 | 0.034 | 0.0000 |
| *Mel. aeglefinus* | *Mer. merlangus* | *G. morhua* (m) | 544,885.0 | 110,788.0 | 103,421.0 | 0.034 | 0.0000 |
| *G. morhua* (m) | *A. glacialis* | *B. saida* | 170,613.0 | 280,258.0 | 131,776.0 | 0.360 | 0.0000 |
| *G. morhua* (m) | *G. macrocephalus* | *B. saida* | 461,022.0 | 87,150.0 | 74,312.0 | 0.080 | 0.0000 |
| *G. morhua* (m) | *G. ogac* | *B. saida* | 463,606.0 | 86,146.6 | 73,616.9 | 0.078 | 0.0000 |
| *G. morhua* (m) | *G. chalcogrammus* | *B. saida* | 699,019.0 | 51,353.8 | 42,041.8 | 0.100 | 0.0000 |
| *G. morhua* (m) | *G. morhua* (s) | *B. saida* | 1,178,120.0 | 6,965.9 | 6,699.1 | 0.020 | 0.0004 |
| *G. morhua* (m) | *G. macrocephalus* | *A. glacialis* | 425,364.0 | 85,291.4 | 71,895.4 | 0.085 | 0.0000 |
| *G. morhua* (m) | *G. ogac* | *A. glacialis* | 428,397.0 | 84,501.5 | 71,280.5 | 0.085 | 0.0000 |
| *G. morhua* (m) | *G. chalcogrammus* | *A. glacialis* | 669,250.0 | 50,390.8 | 40,088.8 | 0.114 | 0.0000 |
| *G. morhua* (m) | *G. morhua* (s) | *A. glacialis* | 1,159,190.0 | 6,748.2 | 6,422.2 | 0.025 | 0.0000 |
| *G. morhua* (m) | *G. chalcogrammus* | *G. macrocephalus* | 337,874.0 | 74,069.5 | 55,581.5 | 0.143 | 0.0000 |
| *G. morhua* (m) | *G. morhua* (s) | *G. macrocephalus* | 820,109.0 | 7,930.4 | 7,495.9 | 0.028 | 0.0000 |
| *G. macrocephalus* | *G. ogac* | *G. chalcogrammus* | 941,162.0 | 9,380.4 | 7,456.4 | 0.114 | 0.0000 |
| *G. macrocephalus* | *G. ogac* | *G. morhua* (s) | 980,086.0 | 13,956.1 | 7,797.4 | 0.283 | 0.0000 |
| *G. macrocephalus* | *G. ogac* | *G. morhua* (m) | 980,709.0 | 14,167.9 | 7,875.9 | 0.285 | 0.0000 |
| *G. morhua* (m) | *G. morhua* (s) | *G. chalcogrammus* | 532,010.0 | 10,099.0 | 9,619.8 | 0.024 | 0.0000 |

[1]Migratory *Gadus morhua* individual LOF1103Z24 sampled at the Lofoten islands

[2]Stationary *Gadus morhua* individual LOF1106Z11 sampled at the Lofoten islands

**Supplementary Table 8:** *Gadus morhua* specimens used for population-level analyses.
For the Newfoundland, Iceland, Møre, and Lofoten populations, we distinguished "migratory" (m) and "stationary" (s) individuals. At all other sampling locations, all individuals were considered stationary. Coverage was measured based on an assumed genome size of 650 Mbp[10]. TL: terminal length in centimeters.

| Specimen ID | ENA accession(s) | Population | Country | Lat. | Lon. | Date | Sampling | TL (cm) | # reads | Coverage |
|---|---|---|---|---|---|---|---|---|---|---|
| TWI1307Z12 | ERR5321892 | Newfoundland (m) | Canada | 49.68 | -54.80 | 07/2013 | Hand line | 76 | 66,970,106 | 10.3 |
| TWI1307Z17 | ERR5321893 | Newfoundland (m) | Canada | 49.68 | -54.80 | 07/2013 | Hand line | 70 | 80,309,398 | 12.4 |
| TWI1307Z07 | ERR5321894 | Newfoundland (s) | Canada | 49.68 | -54.80 | 07/2013 | Hand line | 51 | 76,098,264 | 11.7 |
| TWI1307Z15 | ERR5321895 | Newfoundland (s) | Canada | 49.68 | -54.80 | 07/2013 | Hand line | 67 | 66,823,400 | 10.3 |
| BAT1309Z03 | ERR5321896 | Labrador | Canada | 52.27 | -55.58 | 09/2013 | Hand line | 55 | 60,329,852 | 9.3 |
| BAT1309Z10 | ERR5321897 | Labrador | Canada | 52.27 | -55.58 | 09/2013 | Hand line | 62 | 55,045,914 | 8.5 |
| ICO0304Z23 | ERR5321898 | Iceland (m) | Iceland | 63.50 | -22.11 | 04/2003 | Gillnet | 86 | 110,796,682 | 17.0 |
| ICO0304Z20 | ERR5321899 | Iceland (m) | Iceland | 63.50 | -22.11 | 04/2003 | Gillnet | 81 | 69,704,806 | 10.7 |
| ICC0304Z11 | ERR5321900 | Iceland (s) | Iceland | 63.81 | -21.98 | 04/2003 | Gillnet | NA | 57,098,960 | 8.8 |
| LOF1103Z11 | ERR5321901 | Lofoten (m) | Norway | 68.07 | 13.58 | 03/2011 | Longline | 93 | 67,562,048 | 10.4 |
| NA | ERR1551885[1] | Lofoten (m) | Norway | 67.60 | 12.54 | NA | Longline | NA | 483,953,674 | 74.5 |
| LOF1106Z04 | ERR5321902 | Lofoten (s) | Norway | 68.07 | 13.69 | 06/2011 | Hand line | 64 | 57,883,292 | 8.9 |
| LOF1106Z24 | ERR5321903 | Lofoten (s) | Norway | 68.07 | 13.69 | 06/2011 | Hand line | 68 | 60,765,564 | 9.3 |
| LOF1106Z11[2] | ERR5321904 | Lofoten (s) | Norway | 68.07 | 13.69 | 06/2011 | Hand line | 62 | 61,641,456 | 9.5 |
| AVE1403Z10 | ERR5321905 | Møre (m) | Norway | 63.08 | 7.11 | 03/2014 | Gillnet | 80 | 63,759,112 | 9.8 |
| AVE1403Z11 | ERR5321906 | Møre (m) | Norway | 63.08 | 7.11 | 03/2014 | Gillnet | 102 | 106,219,192 | 16.3 |
| AVE1409Z09[3] | ERR2850387 | Møre (s) | Norway | 63.12 | 7.23 | 09/2014 | Hand line | 74 | 57,939,902 | 8.9 |
| AVE1409Z10[3] | ERR2850388 | Møre (s) | Norway | 63.12 | 7.23 | 09/2014 | Hand line | 65 | 60,093,966 | 9.2 |
| LOW1503Z06[3] | ERR2850515-18 | Suffolk | UK | 52.12 | 1.98 | 03/2015 | Trawling | 58 | 68,126,966 | 10.5 |
| LOW1504Z07[3] | ERR2850579-82 | Suffolk | UK | 52.04 | 1.51 | 04/2015 | Trawling | 53 | 67,523,264 | 10.4 |
| KIE1102Z06[3] | ERR2850464 | Kiel Bight | Germany | 54.76 | 10.05 | 02/2011 | Trawling | 51 | 79,339,236 | 12.2 |
| KIE1103Z20[3] | ERR2850485 | Kiel Bight | Germany | 54.76 | 10.05 | 03/2011 | Trawling | 42 | 89,153,264 | 13.7 |
| BOR1205Z03[3] | ERR2850424-26 | Bornholm Basin | Sweden | 55.29 | 15.72 | 05/2012 | Trawling | 38 | 52,748,670 | 8.1 |
| BOR1205Z07[3] | ERR2850436-38, | Bornholm Basin | Sweden | 55.29 | 15.72 | 05/2012 | Trawling | 45 | 54,990,446 | 8.5 |

[1]Accession of Illumina data from the migratory *Gadus morhua* specimen used for the gadMor2 assembly[10]
[2]Stationary *Gadus morhua* individual used for the gadMor_Stat assembly
[3]Published in Barth et al.[28]

**Supplementary Table 9:** Genetic diversity of *Gadus morhua* populations and outgroups. Genetic diversity $\pi$ was calculated following Ruegg et al.[36]. Genetic diversities for populations inferred to be carriers of the derived arrangement are underlined.

| Species | Population | $\pi$ outside of supergene regions ($\times 10^{-3}$) | $\pi$ in supergene regions ($\times 10^{-3}$) LG 1 | LG 2 | LG 7 | LG 12 |
|---|---|---|---|---|---|---|
| *G. macrocephalus* | NA | 0.363 | 0.302 | 0.283 | 0.369 | 0.376 |
| *G. ogac* | NA | 0.323 | 0.299 | 0.268 | 0.316 | 0.365 |
| *G. chalcogrammus* | NA | 0.905 | 1.031 | 0.993 | 1.213 | 1.068 |
| *G. morhua* | Newfoundland (m) | 0.934 | 0.522 | 0.593 | 0.799 | 0.707 |
| *G. morhua* | Newfoundland (s) | 0.934 | 0.768 | 0.594 | 0.789 | 0.711 |
| *G. morhua* | Labrador | 0.942 | 0.630 | 0.611 | 0.808 | 0.723 |
| *G. morhua* | Iceland (m) | 0.942 | 0.353 | 0.578 | 0.782 | 0.678 |
| *G. morhua* | Iceland (s) | 0.882 | 0.846 | 0.550 | 0.680 | 0.652 |
| *G. morhua* | Lofoten (m) | 1.084 | 0.394 | 0.665 | 0.871 | 0.776 |
| *G. morhua* | Lofoten (s) | 1.061 | 0.843 | 0.683 | 0.812 | 0.489[1] |
| *G. morhua* | Møre (m) | 0.947 | 0.371 | 0.589 | 0.746 | 0.700 |
| *G. morhua* | Møre (s) | 0.974 | 0.780 | 0.619 | 0.694 | 0.660 |
| *G. morhua* | Suffolk | 0.947 | 0.753 | 0.608 | 0.729 | 0.532[1] |
| *G. morhua* | Kiel Bight | 0.935 | 0.738 | 0.607 | 0.769 | 0.518[1] |
| *G. morhua* | Bornholm Basin | 0.904 | 0.722 | 0.379 | 0.680 | 0.487 |
| *G. morhua* | All with derived arrangement[2] | NA | 0.410 | 0.629 | 0.782 | 0.513 |
| *G. morhua* | All with ancestral arrangement[3] | NA | 0.760 | 0.570 | 0.737 | 0.701 |

[1] The derived and ancestral arrangements for the supergene on LG 12 could not be identified with the contig-mapping approach; however, reduced population size in a time inverval after the supergene origin suggests that it was the individuals from the Lofoten (s), Suffolk, and Kiel Bight populations that carried the derived arrangments.

[2] Average genetic diversity across all populations carrying the derived arrangement

[3] Average genetic diversity across all populations carrying the ancestral arrangement

**Supplementary Table 10:** $D_{\mathrm{BBAA}}$-statistics for *Gadus morhua* populations and outgroups. For the calculation of the $D_{\mathrm{BBAA}}$ version of the $D$-statistic[35], all trios are oriented so that the number of "BBAA" patterns ($C_{\mathrm{BBAA}}$) is greater than the numbers of "ABBA" ($C_{\mathrm{ABBA}}$) and "BBAA" ($C_{\mathrm{BABA}}$) patterns. Additionally, the trio is arbitrarily oriented so that the number of "ABBA" ($C_{\mathrm{ABBA}}$) patterns exceeds that of "BBAA" ($C_{\mathrm{BABA}}$) patterns. When the trio is oriented in this way, the $D_{\mathrm{BBAA}}$-statistic is never negative and values significantly greater than zero support introgression between the two species in positions P2 and P3. If the same pair of species in positions P2 and P3 produced significant results with several third species in position P1, only the trio with the highest $D_{\mathrm{BBAA}}$ is listed. Significance was assessed with block-jackknifing. Sites from supergene regions were excluded from this analysis. The highest $D$ values are underlined.

| P1 | P2 | P3 | $C_{\mathrm{BBAA}}$ | $C_{\mathrm{ABBA}}$ | $C_{\mathrm{BABA}}$ | $D_{\mathrm{BBAA}}$ | $p$ |
|---|---|---|---:|---:|---:|---:|---:|
| Lofoten (m) | *G. chalcogrammus* | *G. macrocephalus* | 520,663.0 | 99,351.4 | 84,913.6 | 0.078 | 0.0000 |
| Lofoten (m) | Newfoundland (m) | *G. macrocephalus* | 1,315,150.0 | 13,951.5 | 12,893.6 | 0.039 | 0.0000 |
| Lofoten (m) | Newfoundland (s) | *G. macrocephalus* | 1,315,950.0 | 13,995.8 | 12,968.8 | 0.038 | 0.0000 |
| Lofoten (m) | Labrador | *G. macrocephalus* | 1,283,140.0 | 14,079.2 | 13,066.8 | 0.037 | 0.0000 |
| Lofoten (m) | Møre (m) | *G. macrocephalus* | 1,359,260.0 | 13,246.6 | 12,503.9 | 0.029 | 0.0000 |
| Lofoten (m) | Møre (s) | *G. macrocephalus* | 1,290,310.0 | 12,728.2 | 12,083.8 | 0.026 | 0.0000 |
| Lofoten (m) | Iceland (m) | *G. macrocephalus* | 1,360,800.0 | 13,236.4 | 12,506.8 | 0.028 | 0.0000 |
| Lofoten (m) | Iceland (s) | *G. macrocephalus* | 1,195,570.0 | 11,584.5 | 10,701.2 | 0.040 | 0.0000 |
| Lofoten (m) | Lofoten (s) | *G. macrocephalus* | 1,271,080.0 | 13,134.9 | 12,618.3 | 0.020 | 0.0000 |
| Lofoten (m) | Suffolk | *G. macrocephalus* | 1,337,050.0 | 13,094.1 | 12,316.8 | 0.031 | 0.0000 |
| Lofoten (m) | Kiel Bight | *G. macrocephalus* | 1,349,250.0 | 13,088.5 | 12,375.4 | 0.028 | 0.0000 |
| Lofoten (m) | Bornholm Basin | *G. macrocephalus* | 1,249,840.0 | 12,723.0 | 12,013.0 | 0.029 | 0.0000 |
| *G. macrocephalus* | *G. ogac* | *G. chalcogrammus* | 1,381,720.0 | 11,877.9 | 9,868.6 | 0.092 | 0.0000 |
| *G. macrocephalus* | *G. ogac* | Newfoundland (m) | 1,407,000.0 | 16,512.3 | 9,997.9 | 0.246 | 0.0000 |
| *G. macrocephalus* | *G. ogac* | Newfoundland (s) | 1,409,330.0 | 16,543.0 | 9,954.9 | <u>0.249</u> | 0.0000 |
| *G. macrocephalus* | *G. ogac* | Labrador | 1,362,790.0 | 16,024.5 | 9,682.8 | 0.247 | 0.0000 |
| *G. macrocephalus* | *G. ogac* | Møre (m) | 1,428,900.0 | 16,894.6 | 10,179.4 | 0.248 | 0.0000 |
| *G. macrocephalus* | *G. ogac* | Møre (s) | 1,337,080.0 | 15,864.3 | 9,578.4 | 0.247 | 0.0000 |
| *G. macrocephalus* | *G. ogac* | Lofoten (m) | 1,243,990.0 | 14,961.9 | 8,992.1 | 0.249 | 0.0000 |
| *G. macrocephalus* | *G. ogac* | Iceland (m) | 1,433,930.0 | 16,947.1 | 10,201.1 | 0.248 | 0.0000 |
| *G. macrocephalus* | *G. ogac* | Iceland (s) | 1,244,640.0 | 14,521.0 | 8,801.2 | 0.245 | 0.0000 |
| *G. macrocephalus* | *G. ogac* | Lofoten (s) | 1,306,910.0 | 15,591.3 | 9,381.0 | 0.249 | 0.0000 |
| *G. macrocephalus* | *G. ogac* | Suffolk | 1,399,690.0 | 16,515.4 | 9,967.1 | 0.247 | 0.0000 |
| *G. macrocephalus* | *G. ogac* | Kiel Bight | 1,416,600.0 | 16,741.8 | 10,039.2 | <u>0.250</u> | 0.0000 |
| *G. macrocephalus* | *G. ogac* | Bornholm Basin | 1,303,870.0 | 15,412.2 | 9,298.4 | 0.247 | 0.0000 |
| Lofoten (m) | Newfoundland (m) | *G. chalcogrammus* | 899,928.0 | 19,121.2 | 17,064.7 | 0.057 | 0.0000 |
| Lofoten (m) | Newfoundland (s) | *G. chalcogrammus* | 900,322.0 | 19,129.8 | 17,145.6 | 0.055 | 0.0000 |
| Lofoten (m) | Labrador | *G. chalcogrammus* | 879,634.0 | 19,307.5 | 17,313.0 | 0.054 | 0.0000 |
| Lofoten (m) | Møre (m) | *G. chalcogrammus* | 940,601.0 | 17,992.4 | 16,643.0 | 0.039 | 0.0000 |
| Lofoten (m) | Møre (s) | *G. chalcogrammus* | 892,834.0 | 17,119.2 | 16,108.0 | 0.030 | 0.0000 |
| Lofoten (m) | Iceland (m) | *G. chalcogrammus* | 940,877.0 | 17,881.4 | 16,644.0 | 0.036 | 0.0000 |
| Lofoten (m) | Iceland (s) | *G. chalcogrammus* | 824,358.0 | 15,499.5 | 14,249.0 | 0.042 | 0.0000 |
| Lofoten (m) | Lofoten (s) | *G. chalcogrammus* | 880,879.0 | 17,736.3 | 16,817.5 | 0.027 | 0.0000 |
| Lofoten (m) | Suffolk | *G. chalcogrammus* | 925,054.0 | 17,621.4 | 16,512.5 | 0.032 | 0.0000 |
| Lofoten (m) | Kiel Bight | *G. chalcogrammus* | 933,388.0 | 17,530.1 | 16,578.2 | 0.028 | 0.0000 |
| Lofoten (m) | Bornholm Basin | *G. chalcogrammus* | 864,946.0 | 17,173.2 | 16,101.7 | 0.032 | 0.0000 |
| Lofoten (m) | Møre (m) | Newfoundland (m) | 101,116.0 | 76,042.8 | 73,359.0 | 0.018 | 0.0000 |

**Supplementary Table 10 (continued):** $D_{\mathrm{BBAA}}$-statistics for *Gadus morhua* populations and outgroups.

| P1 | P2 | P3 | $C_{\mathrm{BBAA}}$ | $C_{\mathrm{ABBA}}$ | $C_{\mathrm{BABA}}$ | $D_{\mathrm{BBAA}}$ | $p$ |
|---|---|---|---|---|---|---|---|
| Lofoten (m) | Møre (s) | Newfoundland (m) | 97,687.3 | 73,463.8 | 71,353.9 | 0.015 | 0.0000 |
| Newfoundland (s) | Newfoundland (m) | Lofoten (m) | 100,337.0 | 73,708.3 | 73,166.3 | 0.004 | 0.0199 |
| Lofoten (m) | Iceland (m) | Newfoundland (m) | 99,598.9 | 76,880.4 | 73,436.2 | 0.023 | 0.0000 |
| Lofoten (m) | Iceland (s) | Newfoundland (m) | 87,416.8 | 67,254.8 | 64,090.8 | 0.024 | 0.0000 |
| Lofoten (m) | Lofoten (s) | Newfoundland (m) | 99,879.5 | 75,403.2 | 73,398.0 | 0.013 | 0.0000 |
| Lofoten (m) | Suffolk | Newfoundland (m) | 99,806.2 | 75,106.4 | 72,659.9 | 0.017 | 0.0000 |
| Lofoten (m) | Kiel Bight | Newfoundland (m) | 100,081.0 | 75,274.3 | 72,882.2 | 0.016 | 0.0000 |
| Lofoten (m) | Bornholm Basin | Newfoundland (m) | 95,876.1 | 72,700.8 | 71,137.0 | 0.011 | 0.0000 |
| Lofoten (m) | Møre (m) | Newfoundland (s) | 101,717.0 | 76,179.5 | 73,376.7 | 0.019 | 0.0000 |
| Lofoten (m) | Møre (s) | Newfoundland (s) | 98,290.6 | 73,381.3 | 71,418.9 | 0.014 | 0.0000 |
| Lofoten (m) | Iceland (m) | Newfoundland (s) | 100,066.0 | 77,109.0 | 73,368.2 | 0.025 | 0.0000 |
| Lofoten (m) | Iceland (s) | Newfoundland (s) | 87,938.8 | 67,313.5 | 64,024.0 | 0.025 | 0.0000 |
| Lofoten (m) | Lofoten (s) | Newfoundland (s) | 100,514.0 | 75,274.5 | 73,461.6 | 0.012 | 0.0000 |
| Lofoten (m) | Suffolk | Newfoundland (s) | 100,406.0 | 75,163.5 | 72,707.5 | 0.017 | 0.0000 |
| Lofoten (m) | Kiel Bight | Newfoundland (s) | 100,678.0 | 75,348.4 | 72,961.2 | 0.016 | 0.0000 |
| Lofoten (m) | Bornholm Basin | Newfoundland (s) | 96,373.6 | 72,764.8 | 71,170.7 | 0.011 | 0.0000 |
| Lofoten (m) | Møre (m) | Labrador | 101,470.0 | 76,990.3 | 73,939.4 | 0.020 | 0.0000 |
| Lofoten (m) | Møre (s) | Labrador | 98,044.0 | 74,081.3 | 71,929.0 | 0.015 | 0.0000 |
| Newfoundland (s) | Labrador | Lofoten (m) | 99,742.5 | 75,063.6 | 74,447.7 | 0.004 | 0.0279 |
| Lofoten (m) | Iceland (m) | Labrador | 99,959.3 | 77,749.7 | 73,957.8 | 0.025 | 0.0000 |
| Lofoten (m) | Iceland (s) | Labrador | 87,825.1 | 67,579.0 | 64,608.9 | 0.022 | 0.0000 |
| Lofoten (m) | Lofoten (s) | Labrador | 100,116.0 | 76,114.1 | 73,845.5 | 0.015 | 0.0000 |
| Lofoten (m) | Suffolk | Labrador | 100,325.0 | 75,694.3 | 73,373.0 | 0.016 | 0.0000 |
| Lofoten (m) | Kiel Bight | Labrador | 100,604.0 | 75,824.4 | 73,680.3 | 0.014 | 0.0000 |
| Lofoten (m) | Bornholm Basin | Labrador | 96,187.9 | 73,617.0 | 71,666.0 | 0.013 | 0.0000 |
| Bornholm Basin | Møre (s) | Møre (m) | 82,381.2 | 82,054.0 | 81,135.4 | 0.006 | 0.0001 |
| Iceland (m) | Møre (m) | Lofoten (m) | 86,643.9 | 84,266.6 | 82,604.6 | 0.010 | 0.0000 |
| Bornholm Basin | Møre (m) | Iceland (m) | 85,019.3 | 84,942.5 | 83,017.4 | 0.011 | 0.0000 |
| Bornholm Basin | Møre (m) | Iceland (s) | 73,998.2 | 73,801.4 | 72,299.6 | 0.010 | 0.0000 |
| Bornholm Basin | Lofoten (s) | Møre (m) | 85,727.1 | 83,660.2 | 83,127.0 | 0.003 | 0.0466 |
| Lofoten (s) | Suffolk | Møre (m) | 86,796.1 | 86,639.3 | 86,067.9 | 0.003 | 0.0241 |
| Bornholm Basin | Kiel Bight | Møre (m) | 89,156.6 | 82,461.2 | 81,638.7 | 0.005 | 0.0018 |
| Bornholm Basin | Møre (s) | Lofoten (m) | 81,360.8 | 77,833.7 | 76,767.9 | 0.007 | 0.0000 |
| Bornholm Basin | Møre (s) | Iceland (m) | 83,200.5 | 81,196.5 | 80,013.6 | 0.007 | 0.0000 |
| Bornholm Basin | Møre (s) | Iceland (s) | 73,138.3 | 71,373.7 | 70,332.3 | 0.007 | 0.0051 |
| Bornholm Basin | Lofoten (s) | Møre (s) | 82,126.2 | 81,588.3 | 80,904.9 | 0.004 | 0.0320 |
| Lofoten (s) | Møre (s) | Suffolk | 84,088.9 | 83,849.0 | 83,204.8 | 0.004 | 0.0021 |
| Bornholm Basin | Kiel Bight | Møre (s) | 85,114.4 | 80,467.4 | 79,193.1 | 0.008 | 0.0003 |
| Suffolk | Møre (s) | Bornholm Basin | 81,267.9 | 80,920.2 | 80,378.3 | 0.003 | 0.0523 |
| Iceland (m) | Lofoten (s) | Lofoten (m) | 84,955.3 | 84,141.1 | 82,642.6 | 0.009 | 0.0000 |
| Iceland (m) | Suffolk | Lofoten (m) | 85,394.6 | 83,070.4 | 81,713.4 | 0.008 | 0.0000 |
| Iceland (m) | Kiel Bight | Lofoten (m) | 85,257.8 | 83,440.5 | 82,256.6 | 0.007 | 0.0000 |
| Lofoten (s) | Suffolk | Iceland (m) | 87,585.4 | 86,223.8 | 84,759.5 | 0.009 | 0.0000 |
| Bornholm Basin | Kiel Bight | Iceland (m) | 90,081.5 | 81,472.2 | 80,596.6 | 0.005 | 0.0017 |
| Bornholm Basin | Lofoten (s) | Iceland (s) | 75,788.9 | 72,473.6 | 71,901.6 | 0.004 | 0.0698 |
| Bornholm Basin | Suffolk | Iceland (s) | 73,774.8 | 72,801.1 | 71,399.4 | 0.010 | 0.0001 |
| Bornholm Basin | Kiel Bight | Iceland (s) | 78,755.6 | 71,169.0 | 70,455.3 | 0.005 | 0.0647 |

**Supplementary Table 10 (continued):** $D_{\mathrm{BBAA}}$-statistics for *Gadus morhua* populations and outgroups.

| P1 | P2 | P3 | $C_{\mathrm{BBAA}}$ | $C_{\mathrm{ABBA}}$ | $C_{\mathrm{BABA}}$ | $D_{\mathrm{BBAA}}$ | $p$ |
|---|---|---|---|---|---|---|---|
| Møre (s) | Kiel Bight | Lofoten (s) | 84,291.6 | 84,153.6 | 83,616.2 | 0.003 | 0.0368 |
| Lofoten (s) | Kiel Bight | Suffolk | 86,514.7 | 85,681.9 | 84,988.6 | 0.004 | 0.0018 |
| Møre (m) | Suffolk | Bornholm Basin | 84,024.6 | 83,775.7 | 83,044.5 | 0.004 | 0.0267 |

**Supplementary Table 11:** $D_{\mathrm{fix}}$-statistics for *Gadus morhua* populations and outgroups.
For the calculation of the $D_{\mathrm{fix}}$ version of the $D$-statistic[35], all trios are fixed in their orientation according to an assumed species tree, for which we here use the one inferred under the multi-species coalescent model (Fig. 3b). Additionally, the trio is arbitrarily oriented so that the number of "ABBA" ($C_{\mathrm{ABBA}}$) patterns exceeds that of "BBAA" ($C_{\mathrm{BABA}}$) patterns. When the trio is oriented in this way, the $D_{\mathrm{BBAA}}$-statistic is never negative and values significantly greater than zero support introgression between the two species in positions P2 and P3. If the same pair of species in positions P2 and P3 produced significant results with several third species in position P1, only the trio with the highest $D_{\mathrm{BBAA}}$ is listed. Significance was assessed with block-jackknifing. Sites from supergene regions were excluded from this analysis. The highest $D$ values are underlined.

| P1 | P2 | P3 | $C_{\mathrm{BBAA}}$ | $C_{\mathrm{ABBA}}$ | $C_{\mathrm{BABA}}$ | $D_{\mathrm{BBAA}}$ | $p$ |
|---|---|---|---|---|---|---|---|
| Lofoten (m) | *G. chalcogrammus* | *G. macrocephalus* | 520,663.0 | 99,351.4 | 84,913.6 | 0.078 | 0.0000 |
| Lofoten (m) | Newfoundland (m) | *G. macrocephalus* | 1,315,150.0 | 13,951.5 | 12,893.6 | 0.039 | 0.0000 |
| Lofoten (m) | Newfoundland (s) | *G. macrocephalus* | 1,315,950.0 | 13,995.8 | 12,968.8 | 0.038 | 0.0000 |
| Lofoten (m) | Labrador | *G. macrocephalus* | 1,283,140.0 | 14,079.2 | 13,066.8 | 0.037 | 0.0000 |
| Lofoten (m) | Møre (m) | *G. macrocephalus* | 1,359,260.0 | 13,246.6 | 12,503.9 | 0.029 | 0.0000 |
| Lofoten (m) | Møre (s) | *G. macrocephalus* | 1,290,310.0 | 12,728.2 | 12,083.8 | 0.026 | 0.0000 |
| Lofoten (m) | Iceland (m) | *G. macrocephalus* | 1,360,800.0 | 13,236.4 | 12,506.8 | 0.028 | 0.0000 |
| Lofoten (m) | Iceland (s) | *G. macrocephalus* | 1,195,570.0 | 11,584.5 | 10,701.2 | 0.040 | 0.0000 |
| Lofoten (m) | Lofoten (s) | *G. macrocephalus* | 1,271,080.0 | 13,134.9 | 12,618.3 | 0.020 | 0.0000 |
| Lofoten (m) | Suffolk | *G. macrocephalus* | 1,337,050.0 | 13,094.1 | 12,316.8 | 0.031 | 0.0000 |
| Lofoten (m) | Kiel Bight | *G. macrocephalus* | 1,349,250.0 | 13,088.5 | 12,375.4 | 0.028 | 0.0000 |
| Lofoten (m) | Bornholm Basin | *G. macrocephalus* | 1,249,840.0 | 12,723.0 | 12,013.0 | 0.029 | 0.0000 |
| *G. macrocephalus* | *G. ogac* | *G. chalcogrammus* | 1,381,720.0 | 11,877.9 | 9,868.6 | 0.092 | 0.0000 |
| *G. macrocephalus* | *G. ogac* | Newfoundland (m) | 1,407,000.0 | 16,512.3 | 9,997.9 | 0.246 | 0.0000 |
| *G. macrocephalus* | *G. ogac* | Newfoundland (s) | 1,409,330.0 | 16,543.0 | 9,954.9 | <u>0.249</u> | 0.0000 |
| *G. macrocephalus* | *G. ogac* | Labrador | 1,362,790.0 | 16,024.5 | 9,682.8 | 0.247 | 0.0000 |
| *G. macrocephalus* | *G. ogac* | Møre (m) | 1,428,900.0 | 16,894.6 | 10,179.4 | 0.248 | 0.0000 |
| *G. macrocephalus* | *G. ogac* | Møre (s) | 1,337,080.0 | 15,864.3 | 9,578.4 | 0.247 | 0.0000 |
| *G. macrocephalus* | *G. ogac* | Lofoten (m) | 1,243,990.0 | 14,961.9 | 8,992.1 | 0.249 | 0.0000 |
| *G. macrocephalus* | *G. ogac* | Iceland (m) | 1,433,930.0 | 16,947.1 | 10,201.1 | 0.248 | 0.0000 |
| *G. macrocephalus* | *G. ogac* | Iceland (s) | 1,244,640.0 | 14,521.0 | 8,801.2 | 0.245 | 0.0000 |
| *G. macrocephalus* | *G. ogac* | Lofoten (s) | 1,306,910.0 | 15,591.3 | 9,381.0 | 0.249 | 0.0000 |
| *G. macrocephalus* | *G. ogac* | Suffolk | 1,399,690.0 | 16,515.4 | 9,967.1 | 0.247 | 0.0000 |
| *G. macrocephalus* | *G. ogac* | Kiel Bight | 1,416,600.0 | 16,741.8 | 10,039.2 | <u>0.250</u> | 0.0000 |
| *G. macrocephalus* | *G. ogac* | Bornholm Basin | 1,303,870.0 | 15,412.2 | 9,298.4 | 0.247 | 0.0000 |
| Lofoten (m) | Newfoundland (m) | *G. chalcogrammus* | 899,928.0 | 19,121.2 | 17,064.7 | 0.057 | 0.0000 |

**Supplementary Table 11 (continued):** $D_{\text{fix}}$-statistics for *Gadus morhua* populations and outgroups.

| P1 | P2 | P3 | $C_{\text{BBAA}}$ | $C_{\text{ABBA}}$ | $C_{\text{BABA}}$ | $D_{\text{BBAA}}$ | $p$ |
|---|---|---|---|---|---|---|---|
| Lofoten (m) | Newfoundland (s) | *G. chalcogrammus* | 900,322.0 | 19,129.8 | 17,145.6 | 0.055 | 0.0000 |
| Lofoten (m) | Labrador | *G. chalcogrammus* | 879,634.0 | 19,307.5 | 17,313.0 | 0.054 | 0.0000 |
| Lofoten (m) | Møre (m) | *G. chalcogrammus* | 940,601.0 | 17,992.4 | 16,643.0 | 0.039 | 0.0000 |
| Lofoten (m) | Møre (s) | *G. chalcogrammus* | 892,834.0 | 17,119.2 | 16,108.0 | 0.030 | 0.0000 |
| Lofoten (m) | Iceland (m) | *G. chalcogrammus* | 940,877.0 | 17,881.4 | 16,644.0 | 0.036 | 0.0000 |
| Lofoten (m) | Iceland (s) | *G. chalcogrammus* | 824,358.0 | 15,499.5 | 14,249.0 | 0.042 | 0.0000 |
| Lofoten (m) | Lofoten (s) | *G. chalcogrammus* | 880,879.0 | 17,736.3 | 16,817.5 | 0.027 | 0.0000 |
| Lofoten (m) | Suffolk | *G. chalcogrammus* | 925,054.0 | 17,621.4 | 16,512.5 | 0.032 | 0.0000 |
| Lofoten (m) | Kiel Bight | *G. chalcogrammus* | 933,388.0 | 17,530.1 | 16,578.2 | 0.028 | 0.0000 |
| Lofoten (m) | Bornholm Basin | *G. chalcogrammus* | 864,946.0 | 17,173.2 | 16,101.7 | 0.032 | 0.0000 |
| Lofoten (m) | Møre (m) | Newfoundland (m) | 101,116.0 | 76,042.8 | 73,359.0 | 0.018 | 0.0000 |
| Lofoten (m) | Møre (s) | Newfoundland (m) | 97,687.3 | 73,463.8 | 71,353.9 | 0.015 | 0.0000 |
| Newfoundland (s) | Newfoundland (m) | Lofoten (m) | 100,337.0 | 73,708.3 | 73,166.3 | 0.004 | 0.0199 |
| Lofoten (m) | Iceland (m) | Newfoundland (m) | 99,598.9 | 76,880.4 | 73,436.2 | 0.023 | 0.0000 |
| Lofoten (m) | Iceland (s) | Newfoundland (m) | 87,416.8 | 67,254.8 | 64,090.8 | 0.024 | 0.0000 |
| Lofoten (m) | Lofoten (s) | Newfoundland (m) | 99,879.5 | 75,403.2 | 73,398.0 | 0.013 | 0.0000 |
| Lofoten (m) | Suffolk | Newfoundland (m) | 99,806.2 | 75,106.4 | 72,659.9 | 0.017 | 0.0000 |
| Lofoten (m) | Kiel Bight | Newfoundland (m) | 100,081.0 | 75,274.3 | 72,882.2 | 0.016 | 0.0000 |
| Lofoten (m) | Bornholm Basin | Newfoundland (m) | 95,876.1 | 72,700.8 | 71,137.0 | 0.011 | 0.0000 |
| Lofoten (m) | Møre (m) | Newfoundland (s) | 101,717.0 | 76,179.5 | 73,376.7 | 0.019 | 0.0000 |
| Lofoten (m) | Møre (s) | Newfoundland (s) | 98,290.6 | 73,381.3 | 71,418.9 | 0.014 | 0.0000 |
| Lofoten (m) | Iceland (m) | Newfoundland (s) | 100,066.0 | 77,109.0 | 73,368.2 | 0.025 | 0.0000 |
| Lofoten (m) | Iceland (s) | Newfoundland (s) | 87,938.8 | 67,313.5 | 64,024.0 | 0.025 | 0.0000 |
| Lofoten (m) | Lofoten (s) | Newfoundland (s) | 100,514.0 | 75,274.5 | 73,461.6 | 0.012 | 0.0000 |
| Lofoten (m) | Suffolk | Newfoundland (s) | 100,406.0 | 75,163.5 | 72,707.5 | 0.017 | 0.0000 |
| Lofoten (m) | Kiel Bight | Newfoundland (s) | 100,678.0 | 75,348.4 | 72,961.2 | 0.016 | 0.0000 |
| Lofoten (m) | Bornholm Basin | Newfoundland (s) | 96,373.6 | 72,764.8 | 71,170.7 | 0.011 | 0.0000 |
| Lofoten (m) | Møre (m) | Labrador | 101,470.0 | 76,990.3 | 73,939.4 | 0.020 | 0.0000 |
| Lofoten (m) | Møre (s) | Labrador | 98,044.0 | 74,081.3 | 71,929.0 | 0.015 | 0.0000 |
| Newfoundland (s) | Labrador | Lofoten (m) | 99,742.5 | 75,063.6 | 74,447.7 | 0.004 | 0.0279 |
| Lofoten (m) | Iceland (m) | Labrador | 99,959.3 | 77,749.7 | 73,957.8 | 0.025 | 0.0000 |
| Lofoten (m) | Iceland (s) | Labrador | 87,825.1 | 67,579.0 | 64,608.9 | 0.022 | 0.0000 |
| Lofoten (m) | Lofoten (s) | Labrador | 100,116.0 | 76,114.1 | 73,845.5 | 0.015 | 0.0000 |
| Lofoten (m) | Suffolk | Labrador | 100,325.0 | 75,694.3 | 73,373.0 | 0.016 | 0.0000 |
| Lofoten (m) | Kiel Bight | Labrador | 100,604.0 | 75,824.4 | 73,680.3 | 0.014 | 0.0000 |
| Lofoten (m) | Bornholm Basin | Labrador | 96,187.9 | 73,617.0 | 71,666.0 | 0.013 | 0.0000 |
| Lofoten (m) | Møre (s) | Møre (m) | 80,571.3 | 84,048.2 | 80,947.2 | 0.019 | 0.0000 |
| Lofoten (m) | Iceland (m) | Møre (m) | 82,604.6 | 86,643.9 | 84,266.6 | 0.014 | 0.0000 |
| Lofoten (m) | Iceland (s) | Møre (m) | 72,001.0 | 74,983.3 | 73,196.2 | 0.012 | 0.0000 |
| Lofoten (m) | Lofoten (s) | Møre (m) | 83,276.1 | 86,092.4 | 83,367.1 | 0.016 | 0.0000 |
| Lofoten (m) | Suffolk | Møre (m) | 82,333.2 | 86,034.3 | 82,638.9 | 0.020 | 0.0000 |
| Lofoten (m) | Kiel Bight | Møre (m) | 82,562.5 | 86,296.7 | 82,997.0 | 0.019 | 0.0000 |
| Lofoten (m) | Bornholm Basin | Møre (m) | 79,469.0 | 83,254.0 | 80,923.7 | 0.014 | 0.0000 |
| Lofoten (m) | Møre (s) | Iceland (m) | 81,388.9 | 83,186.0 | 80,164.7 | 0.018 | 0.0000 |
| Lofoten (m) | Møre (s) | Iceland (s) | 71,381.1 | 72,775.3 | 70,473.5 | 0.016 | 0.0000 |
| Lofoten (m) | Møre (s) | Lofoten (s) | 80,085.8 | 84,303.2 | 80,326.7 | 0.024 | 0.0000 |
| Lofoten (m) | Møre (s) | Suffolk | 79,554.5 | 83,557.4 | 79,595.2 | 0.024 | 0.0000 |

**Supplementary Table 11 (continued):** $D_{\text{fix}}$-statistics for *Gadus morhua* populations and outgroups.

| P1 | P2 | P3 | $C_{\text{BBAA}}$ | $C_{\text{ABBA}}$ | $C_{\text{BABA}}$ | $D_{\text{BBAA}}$ | $p$ |
|---|---|---|---|---|---|---|---|
| Lofoten (m) | Møre (s) | Kiel Bight | 79,506.0 | 84,530.7 | 79,462.2 | 0.031 | 0.0000 |
| Lofoten (m) | Møre (s) | Bornholm Basin | 77,833.7 | 81,360.8 | 76,767.9 | 0.029 | 0.0000 |
| Iceland (m) | Lofoten (s) | Lofoten (m) | 84,955.3 | 84,141.1 | 82,642.6 | 0.009 | 0.0000 |
| Iceland (m) | Suffolk | Lofoten (m) | 85,394.6 | 83,070.4 | 81,713.4 | 0.008 | 0.0000 |
| Iceland (m) | Kiel Bight | Lofoten (m) | 85,257.8 | 83,440.5 | 82,256.6 | 0.007 | 0.0000 |
| Lofoten (s) | Suffolk | Iceland (m) | 87,585.4 | 86,223.8 | 84,759.5 | 0.009 | 0.0000 |
| Bornholm Basin | Kiel Bight | Iceland (m) | 90,081.5 | 81,472.2 | 80,596.6 | 0.005 | 0.0017 |
| Bornholm Basin | Lofoten (s) | Iceland (s) | 75,788.9 | 72,473.6 | 71,901.6 | 0.004 | 0.0698 |
| Bornholm Basin | Suffolk | Iceland (s) | 73,774.8 | 72,801.1 | 71,399.4 | 0.010 | 0.0001 |
| Bornholm Basin | Kiel Bight | Iceland (s) | 78,755.6 | 71,169.0 | 70,455.3 | 0.005 | 0.0647 |
| Suffolk | Lofoten (s) | Kiel Bight | 84,988.6 | 86,514.7 | 85,681.9 | 0.005 | 0.0002 |
| Suffolk | Lofoten (s) | Bornholm Basin | 82,724.6 | 84,150.7 | 82,438.2 | 0.010 | 0.0000 |
| Bornholm Basin | Kiel Bight | Suffolk | 87,465.9 | 81,655.5 | 80,793.0 | 0.005 | 0.0022 |

**Supplementary Table 12:** $D_{\text{BBAA}}$-statistics for *Gadus morhua* populations and outgroups within the supergene region on LG 1.
The $D_{\text{BBAA}}$ version of the $D$-statistic[35] was calculated as for Supplementary Table 10, except that only sites from the supergene region on LG 1 were used. The highest $D$ values are underlined.

| P1 | P2 | P3 | $C_{\text{BBAA}}$ | $C_{\text{ABBA}}$ | $C_{\text{BABA}}$ | $D_{\text{BBAA}}$ | $p$ |
|---|---|---|---|---|---|---|---|
| Newfoundland (s) | *G. chalcogrammus* | *G. macrocephalus* | 18,487.3 | 3,724.6 | 3,100.6 | 0.091 | 0.0000 |
| Iceland (m) | Newfoundland (m) | *G. macrocephalus* | 49,542.8 | 212.7 | 185.5 | 0.068 | 0.0050 |
| Kiel Bight | Lofoten (s) | *G. macrocephalus* | 48,224.6 | 316.9 | 290.4 | 0.044 | 0.0169 |
| Kiel Bight | Suffolk | *G. macrocephalus* | 51,567.5 | 315.3 | 290.9 | 0.040 | 0.0312 |
| Kiel Bight | Møre (s) | *G. macrocephalus* | 49,294.1 | 315.4 | 283.0 | 0.054 | 0.0092 |
| Newfoundland (s) | *G. chalcogrammus* | *G. ogac* | 16,967.9 | 3,449.0 | 3,009.4 | 0.068 | 0.0000 |
| *G. macrocephalus* | *G. ogac* | Newfoundland (m) | 44,845.4 | 512.2 | 338.7 | 0.204 | 0.0000 |
| *G. macrocephalus* | *G. ogac* | Iceland (m) | 45,983.8 | 534.5 | 348.5 | 0.211 | 0.0000 |
| *G. macrocephalus* | *G. ogac* | Møre (m) | 45,841.7 | 537.7 | 345.2 | 0.218 | 0.0000 |
| *G. macrocephalus* | *G. ogac* | Lofoten (m) | 39,458.0 | 457.2 | 292.6 | 0.220 | 0.0000 |
| *G. macrocephalus* | *G. ogac* | Newfoundland (s) | 44,555.7 | 466.5 | 334.2 | 0.165 | 0.0000 |
| *G. macrocephalus* | *G. ogac* | Labrador | 42,906.4 | 460.1 | 324.2 | 0.173 | 0.0000 |
| *G. macrocephalus* | *G. ogac* | Iceland (s) | 38,876.5 | 410.5 | 298.0 | 0.159 | 0.0000 |
| *G. macrocephalus* | *G. ogac* | Lofoten (s) | 40,723.6 | 439.4 | 317.3 | 0.161 | 0.0000 |
| *G. macrocephalus* | *G. ogac* | Suffolk | 44,190.2 | 464.3 | 342.1 | 0.152 | 0.0000 |
| *G. macrocephalus* | *G. ogac* | Kiel Bight | 44,891.2 | 472.6 | 346.4 | 0.154 | 0.0000 |
| *G. macrocephalus* | *G. ogac* | Møre (s) | 41,842.6 | 446.7 | 323.9 | 0.159 | 0.0000 |
| *G. macrocephalus* | *G. ogac* | Bornholm Basin | 41,037.4 | 434.5 | 318.9 | 0.153 | 0.0000 |
| Lofoten (m) | Newfoundland (m) | *G. chalcogrammus* | 30,070.9 | 261.1 | 232.4 | 0.058 | 0.0153 |
| Lofoten (m) | Iceland (m) | *G. chalcogrammus* | 33,208.8 | 195.6 | 176.3 | 0.052 | 0.0145 |
| Møre (m) | Newfoundland (s) | *G. chalcogrammus* | 19,552.6 | 2,027.7 | 1,886.4 | 0.036 | 0.0224 |
| Møre (m) | Labrador | *G. chalcogrammus* | 18,468.8 | 2,112.1 | 1,945.4 | 0.041 | 0.0260 |

**Supplementary Table 12 (continued):** $D_{\mathrm{BBAA}}$-statistics for *Gadus morhua* populations and outgroups within the supergene region on LG 1.

| P1 | P2 | P3 | $C_{\mathrm{BBAA}}$ | $C_{\mathrm{ABBA}}$ | $C_{\mathrm{BABA}}$ | $D_{\mathrm{BBAA}}$ | $p$ |
|---|---|---|---|---|---|---|---|
| Bornholm Basin | Iceland (s) | *G. chalcogrammus* | 29,160.5 | 392.2 | 341.5 | 0.069 | 0.0117 |
| Møre (m) | Lofoten (s) | *G. chalcogrammus* | 17,429.4 | 2,049.7 | 1,928.4 | 0.031 | 0.0755 |
| Møre (m) | Suffolk | *G. chalcogrammus* | 18,897.4 | 2,137.4 | 1,989.1 | 0.036 | 0.0361 |
| Møre (m) | Kiel Bight | *G. chalcogrammus* | 19,207.2 | 2,176.5 | 2,019.2 | 0.037 | 0.0194 |
| Bornholm Basin | Møre (s) | *G. chalcogrammus* | 33,024.7 | 367.6 | 330.6 | 0.053 | 0.0139 |
| Møre (m) | Bornholm Basin | *G. chalcogrammus* | 17,629.4 | 2,016.5 | 1,912.0 | 0.027 | 0.0908 |
| Møre (m) | Iceland (m) | Newfoundland (m) | 2,630.7 | 1,295.3 | 1,199.3 | 0.038 | 0.0514 |
| Lofoten (m) | Newfoundland (m) | Newfoundland (s) | 14,465.7 | 1,040.8 | 464.5 | 0.383 | 0.0000 |
| Iceland (m) | Newfoundland (m) | Labrador | 16,935.5 | 1,152.4 | 461.4 | <u>0.428</u> | 0.0000 |
| Iceland (m) | Newfoundland (m) | Iceland (s) | 15,160.8 | 1,002.5 | 414.4 | 0.415 | 0.0000 |
| Iceland (m) | Newfoundland (m) | Lofoten (s) | 16,682.0 | 1,048.9 | 470.8 | 0.380 | 0.0000 |
| Lofoten (m) | Newfoundland (m) | Suffolk | 15,213.9 | 999.9 | 461.1 | 0.369 | 0.0000 |
| Iceland (m) | Newfoundland (m) | Kiel Bight | 17,605.9 | 1,105.1 | 505.8 | 0.372 | 0.0000 |
| Lofoten (m) | Newfoundland (m) | Møre (s) | 14,781.6 | 969.4 | 443.4 | 0.372 | 0.0000 |
| Iceland (m) | Newfoundland (m) | Bornholm Basin | 16,491.6 | 1,040.0 | 476.6 | 0.372 | 0.0000 |
| Lofoten (m) | Iceland (m) | Møre (m) | 1,423.2 | 1422.9 | 1,368.3 | 0.020 | 0.0832 |
| Labrador | Newfoundland (s) | Iceland (m) | 19,409.3 | 1280.1 | 711.1 | 0.286 | 0.0000 |
| Bornholm Basin | Labrador | Iceland (m) | 17,553.5 | 926.0 | 842.9 | 0.047 | 0.0695 |
| Møre (s) | Iceland (s) | Iceland (m) | 16,791.3 | 728.4 | 668.8 | 0.043 | 0.0583 |
| Møre (s) | Suffolk | Iceland (m) | 20,436.4 | 706.2 | 677.3 | 0.021 | 0.0679 |
| Møre (s) | Kiel Bight | Iceland (m) | 20,578.5 | 713.7 | 660.1 | 0.039 | 0.0066 |
| Labrador | Newfoundland (s) | Møre (m) | 19,308.6 | 1,279.2 | 730.8 | 0.273 | 0.0000 |
| Lofoten (m) | Møre (m) | Labrador | 17,641.5 | 535.2 | 426.9 | 0.113 | 0.0016 |
| Lofoten (m) | Møre (m) | Iceland (s) | 15,738.5 | 482.1 | 381.2 | 0.117 | 0.0026 |
| Iceland (m) | Møre (m) | Lofoten (s) | 19,879.4 | 557.8 | 451.5 | 0.105 | 0.0050 |
| Lofoten (m) | Møre (m) | Suffolk | 18,093.3 | 568.8 | 461.0 | 0.105 | 0.0034 |
| Lofoten (m) | Møre (m) | Kiel Bight | 18,248.7 | 581.2 | 455.5 | 0.121 | 0.0008 |
| Lofoten (m) | Møre (m) | Møre (s) | 17,509.5 | 554.2 | 441.1 | 0.114 | 0.0024 |
| Lofoten (m) | Møre (m) | Bornholm Basin | 17,149.7 | 531.9 | 427.6 | 0.109 | 0.0039 |
| Labrador | Newfoundland (s) | Lofoten (m) | 16,820.0 | 1,103.8 | 612.9 | 0.286 | 0.0000 |
| Møre (s) | Iceland (s) | Lofoten (m) | 14,644.3 | 640.7 | 577.0 | 0.052 | 0.0463 |
| Møre (s) | Suffolk | Lofoten (m) | 17,679.3 | 620.2 | 593.3 | 0.022 | 0.0902 |
| Møre (s) | Kiel Bight | Lofoten (m) | 17,772.6 | 629.1 | 568.9 | 0.050 | 0.0018 |
| Suffolk | Iceland (s) | Newfoundland (s) | 3,077.7 | 2,619.6 | 1,718.4 | 0.208 | 0.0000 |
| Newfoundland (s) | Labrador | Iceland (s) | 2,715.7 | 2,432.7 | 1,833.5 | 0.140 | 0.0000 |
| Newfoundland (s) | Labrador | Lofoten (s) | 3,801.3 | 2,357.9 | 1,949.4 | 0.095 | 0.0000 |
| Newfoundland (s) | Labrador | Suffolk | 3,993.3 | 2,425.8 | 1,961.8 | 0.106 | 0.0000 |
| Newfoundland (s) | Labrador | Kiel Bight | 4,008.5 | 2,415.2 | 1,981.8 | 0.099 | 0.0000 |
| Newfoundland (s) | Labrador | Møre (s) | 3,857.1 | 2,357.5 | 1,926.3 | 0.101 | 0.0000 |
| Newfoundland (s) | Labrador | Bornholm Basin | 3,739.3 | 2,312.7 | 1,873.5 | 0.105 | 0.0000 |
| Labrador | Iceland (s) | Lofoten (s) | 2,706.1 | 2,585.8 | 1,673.9 | 0.214 | 0.0000 |
| Labrador | Iceland (s) | Suffolk | 2,786.5 | 2,606.7 | 1,699.8 | 0.211 | 0.0000 |
| Labrador | Iceland (s) | Kiel Bight | 2,812.2 | 2,573.9 | 1,691.7 | 0.207 | 0.0000 |
| Labrador | Iceland (s) | Møre (s) | 2,719.0 | 2,529.8 | 1,643.2 | 0.212 | 0.0000 |
| Labrador | Iceland (s) | Bornholm Basin | 2,642.8 | 2,440.4 | 1,643.7 | 0.195 | 0.0000 |
| Bornholm Basin | Lofoten (s) | Suffolk | 2,372.5 | 2,362.2 | 2,279.9 | 0.018 | 0.0460 |
| Bornholm Basin | Møre (s) | Suffolk | 2,324.7 | 2,322.5 | 2,239.2 | 0.018 | 0.0311 |

**Supplementary Table 13:** $D_{\text{fix}}$-statistics for *Gadus morhua* populations and outgroups within the supergene region on LG 1.

The $D_{\text{fix}}$ version of the $D$-statistic[35] was calculated as for Supplementary Table 10, except that only sites from the supergene region on LG 1 were used and the input tree was the one inferred for the supergene on LG 1 under the multi-species coalescent model (Fig. 4a).

| P1 | P2 | P3 | $C_{\text{BBAA}}$ | $C_{\text{ABBA}}$ | $C_{\text{BABA}}$ | $D_{\text{BBAA}}$ | $p$ |
|---|---|---|---|---|---|---|---|
| Newfoundland (s) | *G. chalcogrammus* | *G. macrocephalus* | 18,487.3 | 3,724.6 | 3,100.6 | 0.091 | 0.0000 |
| Iceland (m) | Newfoundland (m) | *G. macrocephalus* | 49,542.8 | 212.7 | 185.5 | 0.068 | 0.0050 |
| Kiel Bight | Lofoten (s) | *G. macrocephalus* | 48,224.6 | 316.9 | 290.4 | 0.044 | 0.0169 |
| Kiel Bight | Suffolk | *G. macrocephalus* | 51,567.5 | 315.3 | 290.9 | 0.040 | 0.0312 |
| Kiel Bight | Møre (s) | *G. macrocephalus* | 49,294.1 | 315.4 | 283.0 | 0.054 | 0.0092 |
| Newfoundland (s) | *G. chalcogrammus* | *G. ogac* | 16,967.9 | 3,449.0 | 3,009.4 | 0.068 | 0.0000 |
| *G. macrocephalus* | *G. ogac* | Newfoundland (m) | 44,845.4 | 512.2 | 338.7 | 0.204 | 0.0000 |
| *G. macrocephalus* | *G. ogac* | Iceland (m) | 45,983.8 | 534.5 | 348.5 | 0.211 | 0.0000 |
| *G. macrocephalus* | *G. ogac* | Møre (m) | 45,841.7 | 537.7 | 345.2 | 0.218 | 0.0000 |
| *G. macrocephalus* | *G. ogac* | Lofoten (m) | 39,458.0 | 457.2 | 292.6 | 0.220 | 0.0000 |
| *G. macrocephalus* | *G. ogac* | Newfoundland (s) | 44,555.7 | 466.5 | 334.2 | 0.165 | 0.0000 |
| *G. macrocephalus* | *G. ogac* | Labrador | 42,906.4 | 460.1 | 324.2 | 0.173 | 0.0000 |
| *G. macrocephalus* | *G. ogac* | Iceland (s) | 38,876.5 | 410.5 | 298.0 | 0.159 | 0.0000 |
| *G. macrocephalus* | *G. ogac* | Lofoten (s) | 40,723.6 | 439.4 | 317.3 | 0.161 | 0.0000 |
| *G. macrocephalus* | *G. ogac* | Suffolk | 44,190.2 | 464.3 | 342.1 | 0.152 | 0.0000 |
| *G. macrocephalus* | *G. ogac* | Kiel Bight | 44,891.2 | 472.6 | 346.4 | 0.154 | 0.0000 |
| *G. macrocephalus* | *G. ogac* | Møre (s) | 41,842.6 | 446.7 | 323.9 | 0.159 | 0.0000 |
| *G. macrocephalus* | *G. ogac* | Bornholm Basin | 41,037.4 | 434.5 | 318.9 | 0.153 | 0.0000 |
| Lofoten (m) | Newfoundland (m) | *G. chalcogrammus* | 30,070.9 | 261.1 | 232.4 | 0.058 | 0.0153 |
| Lofoten (m) | Iceland (m) | *G. chalcogrammus* | 33,208.8 | 195.6 | 176.3 | 0.052 | 0.0145 |
| Møre (m) | Newfoundland (s) | *G. chalcogrammus* | 19,552.6 | 2,027.7 | 1,886.4 | 0.036 | 0.0224 |
| Møre (m) | Labrador | *G. chalcogrammus* | 18,468.8 | 2,112.1 | 1,945.4 | 0.041 | 0.0260 |
| Bornholm Basin | Iceland (s) | *G. chalcogrammus* | 29,160.5 | 392.2 | 341.5 | 0.069 | 0.0117 |
| Møre (m) | Lofoten (s) | *G. chalcogrammus* | 17,429.4 | 2,049.7 | 1,928.4 | 0.031 | 0.0755 |
| Møre (m) | Suffolk | *G. chalcogrammus* | 18,897.4 | 2,137.4 | 1,989.1 | 0.036 | 0.0361 |
| Møre (m) | Kiel Bight | *G. chalcogrammus* | 19,207.2 | 2,176.5 | 2,019.2 | 0.037 | 0.0194 |
| Bornholm Basin | Møre (s) | *G. chalcogrammus* | 33,024.7 | 367.6 | 330.6 | 0.053 | 0.0139 |
| Møre (m) | Bornholm Basin | *G. chalcogrammus* | 17,629.4 | 2,016.5 | 1,912.0 | 0.027 | 0.0908 |
| Møre (m) | Iceland (m) | Newfoundland (m) | 2,630.7 | 1,295.3 | 1,199.3 | 0.038 | 0.0514 |
| Lofoten (m) | Newfoundland (m) | Newfoundland (s) | 14,465.7 | 1,040.8 | 464.5 | 0.383 | 0.0000 |
| Iceland (m) | Newfoundland (m) | Labrador | 16,935.5 | 1,152.4 | 461.4 | <u>0.428</u> | 0.0000 |
| Iceland (m) | Newfoundland (m) | Iceland (s) | 15,160.8 | 1,002.5 | 414.4 | 0.415 | 0.0000 |
| Iceland (m) | Newfoundland (m) | Lofoten (s) | 16,682.0 | 1,048.9 | 470.8 | 0.380 | 0.0000 |
| Lofoten (m) | Newfoundland (m) | Suffolk | 15,213.9 | 999.9 | 461.1 | 0.369 | 0.0000 |
| Iceland (m) | Newfoundland (m) | Kiel Bight | 17,605.9 | 1,105.1 | 505.8 | 0.372 | 0.0000 |
| Lofoten (m) | Newfoundland (m) | Møre (s) | 14,781.6 | 969.4 | 443.4 | 0.372 | 0.0000 |
| Iceland (m) | Newfoundland (m) | Bornholm Basin | 16,491.6 | 1,040.0 | 476.6 | 0.372 | 0.0000 |
| Labrador | Newfoundland (s) | Iceland (m) | 19,409.3 | 1,280.1 | 711.1 | 0.286 | 0.0000 |
| Bornholm Basin | Labrador | Iceland (m) | 17,553.5 | 926.0 | 842.9 | 0.047 | 0.0695 |
| Møre (s) | Iceland (s) | Iceland (m) | 16,791.3 | 728.4 | 668.8 | 0.043 | 0.0583 |
| Møre (s) | Suffolk | Iceland (m) | 20,436.4 | 706.2 | 677.3 | 0.021 | 0.0679 |
| Møre (s) | Kiel Bight | Iceland (m) | 20,578.5 | 713.7 | 660.1 | 0.039 | 0.0066 |

**Supplementary Table 13 (continued):** $D_{\mathrm{fix}}$-statistics for *Gadus morhua* populations and outgroups within the supergene region on LG 1.

| P1 | P2 | P3 | $C_{\mathrm{BBAA}}$ | $C_{\mathrm{ABBA}}$ | $C_{\mathrm{BABA}}$ | $D_{\mathrm{BBAA}}$ | $p$ |
|---|---|---|---|---|---|---|---|
| Labrador | Newfoundland (s) | Møre (m) | 19,308.6 | 1,279.2 | 730.8 | 0.273 | 0.0000 |
| Lofoten (m) | Møre (m) | Labrador | 17,641.5 | 535.2 | 426.9 | 0.113 | 0.0016 |
| Lofoten (m) | Møre (m) | Iceland (s) | 15,738.5 | 482.1 | 381.2 | 0.117 | 0.0026 |
| Iceland (m) | Møre (m) | Lofoten (s) | 19,879.4 | 557.8 | 451.5 | 0.105 | 0.0050 |
| Lofoten (m) | Møre (m) | Suffolk | 18,093.3 | 568.8 | 461.0 | 0.105 | 0.0034 |
| Lofoten (m) | Møre (m) | Kiel Bight | 18,248.7 | 581.2 | 455.5 | 0.121 | 0.0008 |
| Lofoten (m) | Møre (m) | Møre (s) | 17,509.5 | 554.2 | 441.1 | 0.114 | 0.0024 |
| Lofoten (m) | Møre (m) | Bornholm Basin | 17,149.7 | 531.9 | 427.6 | 0.109 | 0.0039 |
| Labrador | Newfoundland (s) | Lofoten (m) | 16,820.0 | 1,103.8 | 612.9 | 0.286 | 0.0000 |
| Møre (s) | Iceland (s) | Lofoten (m) | 14,644.3 | 640.7 | 577.0 | 0.052 | 0.0463 |
| Møre (s) | Suffolk | Lofoten (m) | 17,679.3 | 620.2 | 593.3 | 0.022 | 0.0902 |
| Møre (s) | Kiel Bight | Lofoten (m) | 17,772.6 | 629.1 | 568.9 | 0.050 | 0.0018 |
| Suffolk | Iceland (s) | Newfoundland (s) | 3,077.7 | 2,619.6 | 1,718.4 | 0.208 | 0.0000 |
| Kiel Bight | Iceland (s) | Labrador | 2,573.9 | 2,812.2 | 1,691.7 | 0.249 | 0.0000 |
| Newfoundland (s) | Labrador | Lofoten (s) | 3,801.3 | 2,357.9 | 1,949.4 | 0.095 | 0.0000 |
| Newfoundland (s) | Labrador | Suffolk | 3,993.3 | 2,425.8 | 1,961.8 | 0.106 | 0.0000 |
| Newfoundland (s) | Labrador | Kiel Bight | 4,008.5 | 2,415.2 | 1,981.8 | 0.099 | 0.0000 |
| Newfoundland (s) | Labrador | Møre (s) | 3,857.1 | 2,357.5 | 1,926.3 | 0.101 | 0.0000 |
| Newfoundland (s) | Labrador | Bornholm Basin | 3,739.3 | 2,312.7 | 1,873.5 | 0.105 | 0.0000 |
| Bornholm Basin | Lofoten (s) | Iceland (s) | 2,931.2 | 1,839.6 | 1,739.2 | 0.028 | 0.0090 |
| Bornholm Basin | Suffolk | Iceland (s) | 2,877.7 | 1,835.4 | 1,749.3 | 0.024 | 0.0182 |
| Bornholm Basin | Møre (s) | Suffolk | 2,324.7 | 2,322.5 | 2,239.2 | 0.018 | 0.0311 |

**Supplementary Table 14:** $D_{\mathrm{BBAA}}$-statistics for *Gadus morhua* populations and outgroups within the supergene region on LG 2.
The $D_{\mathrm{BBAA}}$ version of the $D$-statistic[35] was calculated as for Supplementary Table 10, except that only sites from the supergene region on LG 2 were used. The highest $D$ values are underlined.

| P1 | P2 | P3 | $C_{\mathrm{BBAA}}$ | $C_{\mathrm{ABBA}}$ | $C_{\mathrm{BABA}}$ | $D_{\mathrm{BBAA}}$ | $p$ |
|---|---|---|---|---|---|---|---|
| Labrador | *G. chalcogrammus* | *G. macrocephalus* | 4,523.6 | 1,220.8 | 941.6 | 0.129 | 0.0000 |
| Iceland (m) | Møre (s) | *G. macrocephalus* | 8,549.2 | 451.4 | 329.6 | 0.156 | 0.0002 |
| Bornholm Basin | Suffolk | *G. macrocephalus* | 8,449.9 | 428.6 | 326.2 | 0.136 | 0.0002 |
| Bornholm Basin | Kiel Bight | *G. macrocephalus* | 8,583.7 | 444.1 | 324.8 | 0.155 | 0.0000 |
| Bornholm Basin | Lofoten (s) | *G. macrocephalus* | 7,876.1 | 415.2 | 320.4 | 0.129 | 0.0003 |
| Iceland (m) | Newfoundland (m) | *G. macrocephalus* | 12,739.7 | 127.8 | 108.2 | 0.083 | 0.0325 |
| Lofoten (m) | Labrador | *G. macrocephalus* | 10,727.8 | 113.8 | 101.1 | 0.059 | 0.0910 |
| Lofoten (m) | Bornholm Basin | *G. macrocephalus* | 10,515.5 | 110.4 | 96.6 | 0.066 | 0.0752 |
| Lofoten (m) | Møre (m) | *G. macrocephalus* | 11,448.6 | 115.5 | 105.5 | 0.045 | 0.0714 |
| Labrador | *G. chalcogrammus* | *G. ogac* | 4,161.8 | 1,162.4 | 926.9 | 0.113 | 0.0000 |
| *G. macrocephalus* | *G. ogac* | Møre (s) | 11,666.4 | 157.8 | 84.2 | 0.304 | 0.0000 |
| *G. macrocephalus* | *G. ogac* | Suffolk | 12,617.5 | 169.2 | 94.1 | 0.285 | 0.0000 |
| *G. macrocephalus* | *G. ogac* | Kiel Bight | 12,868.9 | 169.2 | 95.3 | 0.279 | 0.0000 |

**Supplementary Table 14 (continued):** $D_{\mathrm{BBAA}}$-statistics for *Gadus morhua* populations and outgroups within the supergene region on LG 2.

| P1 | P2 | P3 | $C_{\mathrm{BBAA}}$ | $C_{\mathrm{ABBA}}$ | $C_{\mathrm{BABA}}$ | $D_{\mathrm{BBAA}}$ | $p$ |
|---|---|---|---|---|---|---|---|
| *G. macrocephalus* | *G. ogac* | Lofoten (s) | 11,722.3 | 156.2 | 86.2 | 0.289 | 0.0000 |
| *G. macrocephalus* | *G. ogac* | Newfoundland (s) | 13,040.0 | 179.1 | 96.6 | 0.299 | 0.0000 |
| *G. macrocephalus* | *G. ogac* | Newfoundland (m) | 12,934.5 | 177.2 | 95.8 | 0.299 | 0.0000 |
| *G. macrocephalus* | *G. ogac* | Labrador | 12,633.0 | 173.8 | 93.1 | 0.302 | 0.0000 |
| *G. macrocephalus* | *G. ogac* | Iceland (m) | 13,348.1 | 188.5 | 98.5 | 0.314 | 0.0000 |
| *G. macrocephalus* | *G. ogac* | Lofoten (m) | 11,184.0 | 160.1 | 84.8 | 0.308 | 0.0000 |
| *G. macrocephalus* | *G. ogac* | Bornholm Basin | 12,146.6 | 170.1 | 90.4 | 0.306 | 0.0000 |
| *G. macrocephalus* | *G. ogac* | Iceland (s) | 11,465.5 | 162.2 | 81.2 | 0.333 | 0.0000 |
| *G. macrocephalus* | *G. ogac* | Møre (m) | 13,275.2 | 183.5 | 98.0 | 0.304 | 0.0000 |
| Lofoten (m) | Møre (s) | *G. chalcogrammus* | 4,646.8 | 563.8 | 482.2 | 0.078 | 0.0275 |
| Iceland (s) | Suffolk | *G. chalcogrammus* | 5,020.7 | 586.2 | 503.1 | 0.076 | 0.0242 |
| Lofoten (m) | Kiel Bight | *G. chalcogrammus* | 5,025.2 | 615.6 | 516.2 | 0.088 | 0.0095 |
| Lofoten (m) | Lofoten (s) | *G. chalcogrammus* | 4,676.1 | 578.6 | 498.8 | 0.074 | 0.0216 |
| Iceland (s) | Newfoundland (m) | *G. chalcogrammus* | 8,021.9 | 129.8 | 118.1 | 0.047 | 0.0710 |
| Iceland (s) | Labrador | *G. chalcogrammus* | 7,841.8 | 132.3 | 112.9 | 0.079 | 0.0335 |
| Iceland (s) | Iceland (m) | *G. chalcogrammus* | 8,316.7 | 138.8 | 122.4 | 0.063 | 0.0526 |
| Lofoten (m) | Bornholm Basin | *G. chalcogrammus* | 7,701.9 | 152.3 | 109.8 | 0.162 | 0.0003 |
| Lofoten (m) | Møre (m) | *G. chalcogrammus* | 8,400.6 | 155.4 | 136.2 | 0.066 | 0.0159 |
| Suffolk | Møre (s) | Newfoundland (s) | 5,848.9 | 232.1 | 204.8 | 0.062 | 0.0179 |
| Suffolk | Møre (s) | Newfoundland (m) | 5,806.1 | 225.4 | 196.7 | 0.068 | 0.0125 |
| Suffolk | Møre (s) | Labrador | 5,678.6 | 227.7 | 202.2 | 0.059 | 0.0101 |
| Newfoundland (s) | Iceland (m) | Møre (s) | 3,884.3 | 380.7 | 296.8 | 0.124 | 0.0068 |
| Newfoundland (s) | Lofoten (m) | Møre (s) | 3,425.5 | 334.6 | 288.7 | 0.074 | 0.0423 |
| Newfoundland (s) | Bornholm Basin | Møre (s) | 3,612.4 | 372.1 | 260.2 | 0.177 | 0.0001 |
| Newfoundland (s) | Iceland (s) | Møre (s) | 3,433.4 | 323.3 | 254.2 | 0.120 | 0.0034 |
| Newfoundland (s) | Møre (m) | Møre (s) | 3,853.4 | 378.1 | 296.7 | 0.121 | 0.0058 |
| Newfoundland (s) | Newfoundland (m) | Suffolk | 4,180.7 | 298.6 | 268.8 | 0.053 | 0.0214 |
| Newfoundland (s) | Labrador | Suffolk | 4,082.7 | 296.8 | 275.3 | 0.038 | 0.0891 |
| Newfoundland (s) | Iceland (m) | Suffolk | 4,101.5 | 404.2 | 303.7 | 0.142 | 0.0032 |
| Newfoundland (s) | Lofoten (m) | Suffolk | 3,596.5 | 355.6 | 301.9 | 0.082 | 0.0245 |
| Newfoundland (s) | Bornholm Basin | Suffolk | 3,801.8 | 389.0 | 267.9 | 0.184 | 0.0001 |
| Newfoundland (s) | Iceland (s) | Suffolk | 3,576.3 | 342.8 | 247.8 | 0.161 | 0.0001 |
| Newfoundland (s) | Møre (m) | Suffolk | 4,072.1 | 399.8 | 307.8 | 0.130 | 0.0042 |
| Newfoundland (s) | Newfoundland (m) | Kiel Bight | 4,228.3 | 308.4 | 278.4 | 0.051 | 0.0353 |
| Newfoundland (s) | Labrador | Kiel Bight | 4,130.4 | 313.1 | 284.5 | 0.048 | 0.0419 |
| Newfoundland (s) | Iceland (m) | Kiel Bight | 4,146.2 | 416.1 | 314.0 | 0.140 | 0.0043 |
| Newfoundland (s) | Lofoten (m) | Kiel Bight | 3,645.5 | 357.0 | 310.0 | 0.071 | 0.0542 |
| Newfoundland (s) | Bornholm Basin | Kiel Bight | 3,833.9 | 406.4 | 279.6 | 0.185 | 0.0001 |
| Newfoundland (s) | Iceland (s) | Kiel Bight | 3,624.7 | 348.2 | 256.6 | 0.151 | 0.0005 |
| Newfoundland (s) | Møre (m) | Kiel Bight | 4,115.3 | 412.1 | 317.6 | 0.130 | 0.0031 |
| Newfoundland (s) | Labrador | Lofoten (s) | 3,914.1 | 285.2 | 261.1 | 0.044 | 0.0461 |
| Newfoundland (s) | Iceland (m) | Lofoten (s) | 3,933.1 | 381.0 | 290.6 | 0.135 | 0.0049 |
| Newfoundland (s) | Lofoten (m) | Lofoten (s) | 3,471.0 | 339.4 | 291.5 | 0.076 | 0.0446 |
| Newfoundland (s) | Bornholm Basin | Lofoten (s) | 3,649.7 | 383.3 | 258.6 | 0.194 | 0.0001 |
| Newfoundland (s) | Iceland (s) | Lofoten (s) | 3,477.8 | 324.7 | 238.8 | 0.152 | 0.0002 |
| Newfoundland (s) | Møre (m) | Lofoten (s) | 3,909.0 | 379.8 | 294.3 | 0.127 | 0.0045 |
| Møre (m) | Iceland (m) | Newfoundland (s) | 891.3 | 625.1 | 588.6 | 0.030 | 0.0532 |

**Supplementary Table 14 (continued):** $D_{\mathrm{BBAA}}$-statistics for *Gadus morhua* populations and outgroups within the supergene region on LG 2.

| P1 | P2 | P3 | $C_{\mathrm{BBAA}}$ | $C_{\mathrm{ABBA}}$ | $C_{\mathrm{BABA}}$ | $D_{\mathrm{BBAA}}$ | $p$ |
|---|---|---|---|---|---|---|---|
| Møre (m) | Bornholm Basin | Newfoundland (s) | 856.1 | 574.5 | 520.8 | 0.049 | 0.0155 |
| Møre (m) | Iceland (s) | Newfoundland (s) | 776.0 | 546.4 | 471.5 | 0.074 | 0.0051 |
| Møre (m) | Iceland (m) | Newfoundland (m) | 882.2 | 636.2 | 588.7 | 0.039 | 0.0303 |
| Møre (m) | Bornholm Basin | Newfoundland (m) | 846.9 | 577.2 | 520.6 | 0.052 | 0.0203 |
| Møre (m) | Iceland (s) | Newfoundland (m) | 767.2 | 560.9 | 467.3 | 0.091 | 0.0000 |
| Newfoundland (s) | Labrador | Iceland (m) | 793.8 | 661.6 | 600.1 | 0.049 | 0.0070 |
| Newfoundland (s) | Labrador | Lofoten (m) | 761.2 | 602.8 | 559.8 | 0.037 | 0.0037 |
| Newfoundland (s) | Labrador | Bornholm Basin | 737.1 | 627.0 | 572.6 | 0.045 | 0.0353 |
| Newfoundland (s) | Labrador | Iceland (s) | 648.7 | 575.6 | 521.9 | 0.049 | 0.0119 |
| Newfoundland (s) | Labrador | Møre (m) | 819.9 | 652.1 | 588.0 | 0.052 | 0.0029 |
| Lofoten (m) | Bornholm Basin | Iceland (m) | 659.5 | 637.8 | 579.9 | 0.048 | 0.0194 |
| Møre (m) | Bornholm Basin | Lofoten (m) | 639.9 | 630.7 | 592.8 | 0.031 | 0.0870 |

**Supplementary Table 15:** $D_{\mathrm{fix}}$-statistics for *Gadus morhua* populations and outgroups within the supergene region on LG 2.

The $D_{\mathrm{fix}}$ version of the $D$-statistic[35] was calculated as for Supplementary Table 10, except that only sites from the supergene region on LG 2 were used and the input tree was the one inferred for the supergene on LG 2 under the multi-species coalescent model (Fig. 4e). The highest $D$ values are underlined.

| P1 | P2 | P3 | $C_{\mathrm{BBAA}}$ | $C_{\mathrm{ABBA}}$ | $C_{\mathrm{BABA}}$ | $D_{\mathrm{BBAA}}$ | $p$ |
|---|---|---|---|---|---|---|---|
| Labrador | *G. chalcogrammus* | *G. macrocephalus* | 4,523.6 | 1,220.8 | 941.6 | 0.129 | 0.0000 |
| Iceland (m) | Møre (s) | *G. macrocephalus* | 8,549.2 | 451.4 | 329.6 | 0.156 | 0.0002 |
| Bornholm Basin | Suffolk | *G. macrocephalus* | 8,449.9 | 428.6 | 326.2 | 0.136 | 0.0002 |
| Bornholm Basin | Kiel Bight | *G. macrocephalus* | 8,583.7 | 444.1 | 324.8 | 0.155 | 0.0000 |
| Bornholm Basin | Lofoten (s) | *G. macrocephalus* | 7,876.1 | 415.2 | 320.4 | 0.129 | 0.0003 |
| Iceland (m) | Newfoundland (m) | *G. macrocephalus* | 12,739.7 | 127.8 | 108.2 | 0.083 | 0.0325 |
| Lofoten (m) | Labrador | *G. macrocephalus* | 10,727.8 | 113.8 | 101.1 | 0.059 | 0.0910 |
| Lofoten (m) | Bornholm Basin | *G. macrocephalus* | 10,515.5 | 110.4 | 96.6 | 0.066 | 0.0752 |
| Lofoten (m) | Møre (m) | *G. macrocephalus* | 11,448.6 | 115.5 | 105.5 | 0.045 | 0.0714 |
| Labrador | *G. chalcogrammus* | *G. ogac* | 4,161.8 | 1,162.4 | 926.9 | 0.113 | 0.0000 |
| *G. macrocephalus* | *G. ogac* | Møre (s) | 11,666.4 | 157.8 | 84.2 | 0.304 | 0.0000 |
| *G. macrocephalus* | *G. ogac* | Suffolk | 12,617.5 | 169.2 | 94.1 | 0.285 | 0.0000 |
| *G. macrocephalus* | *G. ogac* | Kiel Bight | 12,868.9 | 169.2 | 95.3 | 0.279 | 0.0000 |
| *G. macrocephalus* | *G. ogac* | Lofoten (s) | 11,722.3 | 156.2 | 86.2 | 0.289 | 0.0000 |
| *G. macrocephalus* | *G. ogac* | Newfoundland (s) | 13,040.0 | 179.1 | 96.6 | 0.299 | 0.0000 |
| *G. macrocephalus* | *G. ogac* | Newfoundland (m) | 12,934.5 | 177.2 | 95.8 | 0.299 | 0.0000 |
| *G. macrocephalus* | *G. ogac* | Labrador | 12,633.0 | 173.8 | 93.1 | 0.302 | 0.0000 |
| *G. macrocephalus* | *G. ogac* | Iceland (m) | 13,348.1 | 188.5 | 98.5 | 0.314 | 0.0000 |
| *G. macrocephalus* | *G. ogac* | Lofoten (m) | 11,184.0 | 160.1 | 84.8 | 0.308 | 0.0000 |
| *G. macrocephalus* | *G. ogac* | Bornholm Basin | 12,146.6 | 170.1 | 90.4 | 0.306 | 0.0000 |
| *G. macrocephalus* | *G. ogac* | Iceland (s) | 11,465.5 | 162.2 | 81.2 | $\underline{0.333}$ | 0.0000 |
| *G. macrocephalus* | *G. ogac* | Møre (m) | 13,275.2 | 183.5 | 98.0 | 0.304 | 0.0000 |

**Supplementary Table 15 (continued):** $D_{\text{fix}}$-statistics for *Gadus morhua* populations and outgroups within the supergene region on LG 2.

| P1 | P2 | P3 | $C_{\text{BBAA}}$ | $C_{\text{ABBA}}$ | $C_{\text{BABA}}$ | $D_{\text{BBAA}}$ | $p$ |
|---|---|---|---|---|---|---|---|
| Lofoten (m) | Møre (s) | *G. chalcogrammus* | 4,646.8 | 563.8 | 482.2 | 0.078 | 0.0275 |
| Iceland (s) | Suffolk | *G. chalcogrammus* | 5,020.7 | 586.2 | 503.1 | 0.076 | 0.0242 |
| Lofoten (m) | Kiel Bight | *G. chalcogrammus* | 5,025.2 | 615.6 | 516.2 | 0.088 | 0.0095 |
| Lofoten (m) | Lofoten (s) | *G. chalcogrammus* | 4,676.1 | 578.6 | 498.8 | 0.074 | 0.0216 |
| Iceland (s) | Newfoundland (m) | *G. chalcogrammus* | 8,021.9 | 129.8 | 118.1 | 0.047 | 0.0710 |
| Iceland (s) | Labrador | *G. chalcogrammus* | 7,841.8 | 132.3 | 112.9 | 0.079 | 0.0335 |
| Iceland (s) | Iceland (m) | *G. chalcogrammus* | 8,316.7 | 138.8 | 122.4 | 0.063 | 0.0526 |
| Lofoten (m) | Bornholm Basin | *G. chalcogrammus* | 7,701.9 | 152.3 | 109.8 | 0.162 | 0.0003 |
| Lofoten (m) | Møre (m) | *G. chalcogrammus* | 8,400.6 | 155.4 | 136.2 | 0.066 | 0.0159 |
| Suffolk | Møre (s) | Newfoundland (s) | 5,848.9 | 232.1 | 204.8 | 0.062 | 0.0179 |
| Suffolk | Møre (s) | Newfoundland (m) | 5,806.1 | 225.4 | 196.7 | 0.068 | 0.0125 |
| Suffolk | Møre (s) | Labrador | 5,678.6 | 227.7 | 202.2 | 0.059 | 0.0101 |
| Newfoundland (s) | Iceland (m) | Møre (s) | 3,884.3 | 380.7 | 296.8 | 0.124 | 0.0068 |
| Newfoundland (s) | Lofoten (m) | Møre (s) | 3,425.5 | 334.6 | 288.7 | 0.074 | 0.0423 |
| Newfoundland (s) | Bornholm Basin | Møre (s) | 3,612.4 | 372.1 | 260.2 | 0.177 | 0.0001 |
| Newfoundland (s) | Iceland (s) | Møre (s) | 3,433.4 | 323.3 | 254.2 | 0.120 | 0.0034 |
| Newfoundland (s) | Møre (m) | Møre (s) | 3,853.4 | 378.1 | 296.7 | 0.121 | 0.0058 |
| Newfoundland (s) | Newfoundland (m) | Suffolk | 4,180.7 | 298.6 | 268.8 | 0.053 | 0.0214 |
| Newfoundland (s) | Labrador | Suffolk | 4,082.7 | 296.8 | 275.3 | 0.038 | 0.0891 |
| Newfoundland (s) | Iceland (m) | Suffolk | 4,101.5 | 404.2 | 303.7 | 0.142 | 0.0032 |
| Newfoundland (s) | Lofoten (m) | Suffolk | 3,596.5 | 355.6 | 301.9 | 0.082 | 0.0245 |
| Newfoundland (s) | Bornholm Basin | Suffolk | 3,801.8 | 389.0 | 267.9 | 0.184 | 0.0001 |
| Newfoundland (s) | Iceland (s) | Suffolk | 3,576.3 | 342.8 | 247.8 | 0.161 | 0.0001 |
| Newfoundland (s) | Møre (m) | Suffolk | 4,072.1 | 399.8 | 307.8 | 0.130 | 0.0042 |
| Newfoundland (s) | Newfoundland (m) | Kiel Bight | 4,228.3 | 308.4 | 278.4 | 0.051 | 0.0353 |
| Newfoundland (s) | Labrador | Kiel Bight | 4,130.4 | 313.1 | 284.5 | 0.048 | 0.0419 |
| Newfoundland (s) | Iceland (m) | Kiel Bight | 4,146.2 | 416.1 | 314.0 | 0.140 | 0.0043 |
| Newfoundland (s) | Lofoten (m) | Kiel Bight | 3,645.5 | 357.0 | 310.0 | 0.071 | 0.0542 |
| Newfoundland (s) | Bornholm Basin | Kiel Bight | 3,833.9 | 406.4 | 279.6 | 0.185 | 0.0001 |
| Newfoundland (s) | Iceland (s) | Kiel Bight | 3,624.7 | 348.2 | 256.6 | 0.151 | 0.0005 |
| Newfoundland (s) | Møre (m) | Kiel Bight | 4,115.3 | 412.1 | 317.6 | 0.130 | 0.0031 |
| Newfoundland (s) | Labrador | Lofoten (s) | 3,914.1 | 285.2 | 261.1 | 0.044 | 0.0461 |
| Newfoundland (s) | Iceland (m) | Lofoten (s) | 3,933.1 | 381.0 | 290.6 | 0.135 | 0.0049 |
| Newfoundland (s) | Lofoten (m) | Lofoten (s) | 3,471.0 | 339.4 | 291.5 | 0.076 | 0.0446 |
| Newfoundland (s) | Bornholm Basin | Lofoten (s) | 3,649.7 | 383.3 | 258.6 | 0.194 | 0.0001 |
| Newfoundland (s) | Iceland (s) | Lofoten (s) | 3,477.8 | 324.7 | 238.8 | 0.152 | 0.0002 |
| Newfoundland (s) | Møre (m) | Lofoten (s) | 3,909.0 | 379.8 | 294.3 | 0.127 | 0.0045 |
| Møre (m) | Iceland (m) | Newfoundland (s) | 891.3 | 625.1 | 588.6 | 0.030 | 0.0532 |
| Møre (m) | Bornholm Basin | Newfoundland (s) | 856.1 | 574.5 | 520.8 | 0.049 | 0.0155 |
| Møre (m) | Iceland (s) | Newfoundland (s) | 776.0 | 546.4 | 471.5 | 0.074 | 0.0051 |
| Møre (m) | Iceland (m) | Newfoundland (m) | 882.2 | 636.2 | 588.7 | 0.039 | 0.0303 |
| Møre (m) | Bornholm Basin | Newfoundland (m) | 846.9 | 577.2 | 520.6 | 0.052 | 0.0203 |
| Møre (m) | Iceland (s) | Newfoundland (m) | 767.2 | 560.9 | 467.3 | 0.091 | 0.0000 |
| Newfoundland (s) | Labrador | Iceland (m) | 793.8 | 661.6 | 600.1 | 0.049 | 0.0070 |
| Newfoundland (s) | Labrador | Lofoten (m) | 761.2 | 602.8 | 559.8 | 0.037 | 0.0037 |
| Newfoundland (s) | Labrador | Bornholm Basin | 737.1 | 627.0 | 572.6 | 0.045 | 0.0353 |
| Newfoundland (s) | Labrador | Iceland (s) | 648.7 | 575.6 | 521.9 | 0.049 | 0.0119 |

**Supplementary Table 15 (continued):** $D_{\text{fix}}$-statistics for *Gadus morhua* populations and outgroups within the supergene region on LG 2.

| P1 | P2 | P3 | $C_{\text{BBAA}}$ | $C_{\text{ABBA}}$ | $C_{\text{BABA}}$ | $D_{\text{BBAA}}$ | $p$ |
|---|---|---|---|---|---|---|---|
| Newfoundland (s) | Labrador | Møre (m) | 819.9 | 652.1 | 588.0 | 0.052 | 0.0029 |
| Møre (m) | Bornholm Basin | Lofoten (m) | 639.9 | 630.7 | 592.8 | 0.031 | 0.0870 |
| Iceland (m) | Lofoten (m) | Iceland (s) | 547.9 | 600.6 | 559.8 | 0.035 | 0.0693 |

**Supplementary Table 16:** $D_{\text{BBAA}}$-statistics for *Gadus morhua* populations and outgroups within the supergene region on LG 7.

The $D_{\text{BBAA}}$ version of the $D$-statistic[35] was calculated as for Supplementary Table 10, except that only sites from the supergene region on LG 7 were used. The highest $D$ values are underlined.

| P1 | P2 | P3 | $C_{\text{BBAA}}$ | $C_{\text{ABBA}}$ | $C_{\text{BABA}}$ | $D_{\text{BBAA}}$ | $p$ |
|---|---|---|---|---|---|---|---|
| Lofoten (m) | *G. chalcogrammus* | *G. macrocephalus* | 9,783.1 | 1,774.3 | 1,527.7 | 0.075 | 0.0000 |
| Labrador | Møre (s) | *G. macrocephalus* | 19,051.8 | 655.2 | 571.7 | 0.068 | 0.0139 |
| Lofoten (m) | Bornholm Basin | *G. macrocephalus* | 17,032.2 | 599.9 | 511.2 | 0.080 | 0.0067 |
| Labrador | Lofoten (s) | *G. macrocephalus* | 18,511.5 | 638.6 | 565.2 | 0.061 | 0.0098 |
| Labrador | Suffolk | *G. macrocephalus* | 19,593.7 | 664.2 | 582.2 | 0.066 | 0.0066 |
| Labrador | Kiel Bight | *G. macrocephalus* | 19,806.5 | 667.7 | 572.7 | 0.077 | 0.0017 |
| Labrador | Newfoundland (m) | *G. macrocephalus* | 30,404.5 | 130.4 | 118.6 | 0.048 | 0.0301 |
| Labrador | Newfoundland (s) | *G. macrocephalus* | 30,383.7 | 122.5 | 111.2 | 0.048 | 0.0655 |
| Lofoten (m) | Iceland (m) | *G. macrocephalus* | 28,753.7 | 131.1 | 116.6 | 0.058 | 0.0394 |
| Lofoten (m) | Møre (m) | *G. macrocephalus* | 28,871.1 | 128.8 | 114.2 | 0.060 | 0.0676 |
| *G. macrocephalus* | *G. ogac* | *G. chalcogrammus* | 26,488.1 | 215.9 | 149.9 | 0.180 | 0.0000 |
| *G. macrocephalus* | *G. ogac* | Møre (s) | 25,584.8 | 240.8 | 152.9 | 0.223 | 0.0000 |
| *G. macrocephalus* | *G. ogac* | Bornholm Basin | 24,421.1 | 241.8 | 145.5 | 0.249 | 0.0000 |
| *G. macrocephalus* | *G. ogac* | Lofoten (s) | 24,835.1 | 229.9 | 151.7 | 0.205 | 0.0000 |
| *G. macrocephalus* | *G. ogac* | Suffolk | 26,375.0 | 248.0 | 160.5 | 0.214 | 0.0000 |
| *G. macrocephalus* | *G. ogac* | Kiel Bight | 26,699.2 | 247.8 | 158.5 | 0.220 | 0.0000 |
| *G. macrocephalus* | *G. ogac* | Newfoundland (m) | 26,935.1 | 313.9 | 154.9 | 0.339 | 0.0000 |
| *G. macrocephalus* | *G. ogac* | Newfoundland (s) | 26,915.3 | 317.4 | 151.6 | 0.354 | 0.0000 |
| *G. macrocephalus* | *G. ogac* | Labrador | 26,105.4 | 304.8 | 147.0 | 0.349 | 0.0000 |
| *G. macrocephalus* | *G. ogac* | Iceland (m) | 27,210.5 | 317.7 | 154.9 | 0.344 | 0.0000 |
| *G. macrocephalus* | *G. ogac* | Iceland (s) | 23,784.4 | 286.6 | 130.1 | <u>0.376</u> | 0.0000 |
| *G. macrocephalus* | *G. ogac* | Møre (m) | 27,158.5 | 317.1 | 154.5 | 0.345 | 0.0000 |
| *G. macrocephalus* | *G. ogac* | Lofoten (m) | 24,258.4 | 285.7 | 136.6 | 0.353 | 0.0000 |
| Lofoten (m) | Møre (s) | *G. chalcogrammus* | 10,368.6 | 1,050.6 | 967.4 | 0.041 | 0.0414 |
| Lofoten (m) | Bornholm Basin | *G. chalcogrammus* | 9,899.0 | 999.6 | 920.9 | 0.041 | 0.0375 |
| Lofoten (m) | Lofoten (s) | *G. chalcogrammus* | 10,105.6 | 1,053.3 | 952.7 | 0.050 | 0.0151 |
| Lofoten (m) | Suffolk | *G. chalcogrammus* | 10,589.6 | 1,068.7 | 977.6 | 0.044 | 0.0259 |
| Lofoten (m) | Kiel Bight | *G. chalcogrammus* | 10,736.8 | 1,077.0 | 983.4 | 0.045 | 0.0339 |
| Lofoten (m) | Møre (m) | *G. chalcogrammus* | 20,737.2 | 173.1 | 150.0 | 0.072 | 0.0138 |
| Labrador | Newfoundland (m) | Møre (s) | 11,167.1 | 343.9 | 313.4 | 0.046 | 0.0207 |
| Labrador | Iceland (m) | Møre (s) | 10,976.7 | 379.2 | 343.2 | 0.050 | 0.0397 |
| Labrador | Møre (m) | Møre (s) | 10,921.5 | 390.2 | 353.8 | 0.049 | 0.0812 |

**Supplementary Table 16 (continued):** $D_{\mathrm{BBAA}}$-statistics for *Gadus morhua* populations and outgroups within the supergene region on LG 7.

| P1 | P2 | P3 | $C_{\mathrm{BBAA}}$ | $C_{\mathrm{ABBA}}$ | $C_{\mathrm{BABA}}$ | $D_{\mathrm{BBAA}}$ | $p$ |
|---|---|---|---|---|---|---|---|
| Møre (s) | Lofoten (s) | Bornholm Basin | 1,266.1 | 1,216.4 | 1,142.6 | 0.031 | 0.0089 |
| Labrador | Newfoundland (m) | Bornholm Basin | 10,769.9 | 328.2 | 302.0 | 0.042 | 0.0593 |
| Lofoten (s) | Bornholm Basin | Newfoundland (s) | 15,742.8 | 227.9 | 210.3 | 0.040 | 0.0900 |
| Lofoten (s) | Bornholm Basin | Labrador | 15,430.6 | 224.1 | 207.1 | 0.039 | 0.0866 |
| Newfoundland (s) | Iceland (m) | Bornholm Basin | 10,824.1 | 354.0 | 320.3 | 0.050 | 0.0452 |
| Suffolk | Bornholm Basin | Iceland (s) | 14,357.4 | 197.8 | 180.0 | 0.047 | 0.0580 |
| Suffolk | Bornholm Basin | Møre (m) | 16,305.9 | 225.9 | 209.4 | 0.038 | 0.0960 |
| Suffolk | Bornholm Basin | Lofoten (m) | 14,674.8 | 200.7 | 184.9 | 0.041 | 0.0828 |
| Labrador | Newfoundland (m) | Lofoten (s) | 11,054.0 | 333.8 | 309.9 | 0.037 | 0.0482 |
| Newfoundland (s) | Iceland (m) | Lofoten (s) | 11,086.1 | 364.3 | 330.7 | 0.048 | 0.0601 |
| Newfoundland (s) | Iceland (m) | Suffolk | 11,448.1 | 374.7 | 344.3 | 0.042 | 0.0826 |
| Labrador | Newfoundland (m) | Kiel Bight | 11,467.6 | 350.8 | 322.7 | 0.042 | 0.0466 |
| Labrador | Iceland (m) | Kiel Bight | 11,283.4 | 385.2 | 355.0 | 0.041 | 0.0632 |
| Iceland (s) | Iceland (m) | Newfoundland (m) | 1,486.2 | 1,023.8 | 945.8 | 0.040 | 0.0070 |
| Lofoten (m) | Iceland (m) | Newfoundland (s) | 1,524.5 | 1,144.1 | 1,044.4 | 0.046 | 0.0066 |
| Newfoundland (m) | Newfoundland (s) | Møre (m) | 1,564.5 | 1,221.2 | 1,119.0 | 0.044 | 0.0665 |
| Lofoten (m) | Iceland (m) | Labrador | 1,557.8 | 1,172.0 | 1,056.5 | 0.052 | 0.0026 |
| Newfoundland (m) | Labrador | Iceland (s) | 1,424.0 | 1,082.1 | 997.6 | 0.041 | 0.0202 |
| Newfoundland (m) | Labrador | Møre (m) | 1,604.8 | 1,206.8 | 1,127.5 | 0.034 | 0.0502 |
| Lofoten (m) | Møre (m) | Iceland (m) | 1,313.3 | 1,285.3 | 1,150.1 | 0.056 | 0.0017 |

**Supplementary Table 17:** $D_{\mathrm{fix}}$-statistics for *Gadus morhua* populations and outgroups within the supergene region on LG 7.
The $D_{\mathrm{fix}}$ version of the $D$-statistic[35] was calculated as for Supplementary Table 10, except that only sites from the supergene region on LG 7 were used and the input tree was the one inferred for the supergene on LG 7 under the multi-species coalescent model (Fig. 5a). The highest $D$ values are underlined.

| P1 | P2 | P3 | $C_{\mathrm{BBAA}}$ | $C_{\mathrm{ABBA}}$ | $C_{\mathrm{BABA}}$ | $D_{\mathrm{BBAA}}$ | $p$ |
|---|---|---|---|---|---|---|---|
| Lofoten (m) | *G. chalcogrammus* | *G. macrocephalus* | 9,783.1 | 1,774.3 | 1,527.7 | 0.075 | 0.0000 |
| Labrador | Møre (s) | *G. macrocephalus* | 19,051.8 | 655.2 | 571.7 | 0.068 | 0.0139 |
| Lofoten (m) | Bornholm Basin | *G. macrocephalus* | 17,032.2 | 599.9 | 511.2 | 0.080 | 0.0067 |
| Labrador | Lofoten (s) | *G. macrocephalus* | 18,511.5 | 638.6 | 565.2 | 0.061 | 0.0098 |
| Labrador | Suffolk | *G. macrocephalus* | 19,593.7 | 664.2 | 582.2 | 0.066 | 0.0066 |
| Labrador | Kiel Bight | *G. macrocephalus* | 19,806.5 | 667.7 | 572.7 | 0.077 | 0.0017 |
| Labrador | Newfoundland (m) | *G. macrocephalus* | 30,404.5 | 130.4 | 118.6 | 0.048 | 0.0301 |
| Labrador | Newfoundland (s) | *G. macrocephalus* | 30,383.7 | 122.5 | 111.2 | 0.048 | 0.0655 |
| Lofoten (m) | Iceland (m) | *G. macrocephalus* | 28,753.7 | 131.1 | 116.6 | 0.058 | 0.0394 |
| Lofoten (m) | Møre (m) | *G. macrocephalus* | 28,871.1 | 128.8 | 114.2 | 0.060 | 0.0676 |
| *G. macrocephalus* | *G. ogac* | *G. chalcogrammus* | 26,488.1 | 215.9 | 149.9 | 0.180 | 0.0000 |
| *G. macrocephalus* | *G. ogac* | Møre (s) | 25,584.8 | 240.8 | 152.9 | 0.223 | 0.0000 |
| *G. macrocephalus* | *G. ogac* | Bornholm Basin | 24,421.1 | 241.8 | 145.5 | 0.249 | 0.0000 |
| *G. macrocephalus* | *G. ogac* | Lofoten (s) | 24,835.1 | 229.9 | 151.7 | 0.205 | 0.0000 |

**Supplementary Table 17 (continued):** $D_{\mathrm{fix}}$-statistics for *Gadus morhua* populations and outgroups within the supergene region on LG 7.

| P1 | P2 | P3 | $C_{\mathrm{BBAA}}$ | $C_{\mathrm{ABBA}}$ | $C_{\mathrm{BABA}}$ | $D_{\mathrm{BBAA}}$ | $p$ |
|---|---|---|---|---|---|---|---|
| *G. macrocephalus* | *G. ogac* | Suffolk | 26,375.0 | 248.0 | 160.5 | 0.214 | 0.0000 |
| *G. macrocephalus* | *G. ogac* | Kiel Bight | 26,699.2 | 247.8 | 158.5 | 0.220 | 0.0000 |
| *G. macrocephalus* | *G. ogac* | Newfoundland (m) | 26,935.1 | 313.9 | 154.9 | 0.339 | 0.0000 |
| *G. macrocephalus* | *G. ogac* | Newfoundland (s) | 26,915.3 | 317.4 | 151.6 | 0.354 | 0.0000 |
| *G. macrocephalus* | *G. ogac* | Labrador | 26,105.4 | 304.8 | 147.0 | 0.349 | 0.0000 |
| *G. macrocephalus* | *G. ogac* | Iceland (m) | 27,210.5 | 317.7 | 154.9 | 0.344 | 0.0000 |
| *G. macrocephalus* | *G. ogac* | Iceland (s) | 23,784.4 | 286.6 | 130.1 | 0.376 | 0.0000 |
| *G. macrocephalus* | *G. ogac* | Møre (m) | 27,158.5 | 317.1 | 154.5 | 0.345 | 0.0000 |
| *G. macrocephalus* | *G. ogac* | Lofoten (m) | 24,258.4 | 285.7 | 136.6 | 0.353 | 0.0000 |
| Lofoten (m) | Møre (s) | *G. chalcogrammus* | 10,368.6 | 1,050.6 | 967.4 | 0.041 | 0.0414 |
| Lofoten (m) | Bornholm Basin | *G. chalcogrammus* | 9,899.0 | 999.6 | 920.9 | 0.041 | 0.0375 |
| Lofoten (m) | Lofoten (s) | *G. chalcogrammus* | 10,105.6 | 1,053.3 | 952.7 | 0.050 | 0.0151 |
| Lofoten (m) | Suffolk | *G. chalcogrammus* | 10,589.6 | 1,068.7 | 977.6 | 0.044 | 0.0259 |
| Lofoten (m) | Kiel Bight | *G. chalcogrammus* | 10,736.8 | 1,077.0 | 983.4 | 0.045 | 0.0339 |
| Lofoten (m) | Møre (m) | *G. chalcogrammus* | 20,737.2 | 173.1 | 150.0 | 0.072 | 0.0138 |
| Kiel Bight | Lofoten (s) | Møre (s) | 1,217.9 | 1,376.1 | 1,174.9 | 0.079 | 0.0001 |
| Kiel Bight | Suffolk | Møre (s) | 1,234.6 | 1,307.1 | 1,193.7 | 0.045 | 0.0015 |
| Labrador | Newfoundland (m) | Møre (s) | 11,167.1 | 343.9 | 313.4 | 0.046 | 0.0207 |
| Labrador | Iceland (m) | Møre (s) | 10,976.7 | 379.2 | 343.2 | 0.050 | 0.0397 |
| Labrador | Møre (m) | Møre (s) | 10,921.5 | 390.2 | 353.8 | 0.049 | 0.0812 |
| Møre (s) | Lofoten (s) | Bornholm Basin | 1,266.1 | 1,216.4 | 1,142.6 | 0.031 | 0.0089 |
| Labrador | Newfoundland (m) | Bornholm Basin | 10,769.9 | 328.2 | 302.0 | 0.042 | 0.0593 |
| Lofoten (s) | Bornholm Basin | Newfoundland (s) | 15,742.8 | 227.9 | 210.3 | 0.040 | 0.0900 |
| Lofoten (s) | Bornholm Basin | Labrador | 15,430.6 | 224.1 | 207.1 | 0.039 | 0.0866 |
| Newfoundland (s) | Iceland (m) | Bornholm Basin | 10,824.1 | 354.0 | 320.3 | 0.050 | 0.0452 |
| Suffolk | Bornholm Basin | Iceland (s) | 14,357.4 | 197.8 | 180.0 | 0.047 | 0.0580 |
| Suffolk | Bornholm Basin | Møre (m) | 16,305.9 | 225.9 | 209.4 | 0.038 | 0.0960 |
| Suffolk | Bornholm Basin | Lofoten (m) | 14,674.8 | 200.7 | 184.9 | 0.041 | 0.0828 |
| Kiel Bight | Suffolk | Lofoten (s) | 1,230.0 | 1,310.7 | 1,232.1 | 0.031 | 0.0259 |
| Labrador | Newfoundland (m) | Lofoten (s) | 11,054.0 | 333.8 | 309.9 | 0.037 | 0.0482 |
| Newfoundland (s) | Iceland (m) | Lofoten (s) | 11,086.1 | 364.3 | 330.7 | 0.048 | 0.0601 |
| Newfoundland (s) | Iceland (m) | Suffolk | 11,448.1 | 374.7 | 344.3 | 0.042 | 0.0826 |
| Labrador | Newfoundland (m) | Kiel Bight | 11,467.6 | 350.8 | 322.7 | 0.042 | 0.0466 |
| Labrador | Iceland (m) | Kiel Bight | 11,283.4 | 385.2 | 355.0 | 0.041 | 0.0632 |
| Iceland (s) | Iceland (m) | Newfoundland (m) | 1,486.2 | 1,023.8 | 945.8 | 0.040 | 0.0070 |
| Lofoten (m) | Iceland (m) | Newfoundland (s) | 1,524.5 | 1,144.1 | 1,044.4 | 0.046 | 0.0066 |
| Newfoundland (m) | Newfoundland (s) | Møre (m) | 1,564.5 | 1,221.2 | 1,119.0 | 0.044 | 0.0665 |
| Lofoten (m) | Iceland (m) | Labrador | 1,557.8 | 1,172.0 | 1,056.5 | 0.052 | 0.0026 |
| Newfoundland (m) | Labrador | Iceland (s) | 1,424.0 | 1,082.1 | 997.6 | 0.041 | 0.0202 |
| Newfoundland (m) | Labrador | Møre (m) | 1,604.8 | 1,206.8 | 1,127.5 | 0.034 | 0.0502 |
| Lofoten (m) | Møre (m) | Iceland (m) | 1,313.3 | 1,285.3 | 1,150.1 | 0.056 | 0.0017 |
| Iceland (m) | Iceland (s) | Lofoten (m) | 1,077.1 | 1,165.1 | 1,035.7 | 0.059 | 0.0002 |

**Supplementary Table 18:** $D_{\mathrm{BBAA}}$-statistics for *Gadus morhua* populations and outgroups within the supergene region on LG 12.
The $D_{\mathrm{BBAA}}$ version of the $D$-statistic[35] was calculated as for Supplementary Table 10, except that only sites from the supergene region on LG 12 were used. The highest $D$ values are underlined.

| P1 | P2 | P3 | $C_{\mathrm{BBAA}}$ | $C_{\mathrm{ABBA}}$ | $C_{\mathrm{BABA}}$ | $D_{\mathrm{BBAA}}$ | $p$ |
|---|---|---|---:|---:|---:|---:|---:|
| Bornholm Basin | *G. chalcogrammus* | *G. macrocephalus* | 13,979.6 | 2,751.8 | 2,566.6 | 0.035 | 0.0179 |
| Bornholm Basin | Lofoten (s) | *G. macrocephalus* | 23,094.0 | 363.2 | 293.2 | 0.107 | 0.0094 |
| Bornholm Basin | Suffolk | *G. macrocephalus* | 33,148.2 | 491.0 | 391.6 | 0.113 | 0.0005 |
| Bornholm Basin | Kiel Bight | *G. macrocephalus* | 33,505.7 | 485.5 | 408.0 | 0.087 | 0.0026 |
| *G. macrocephalus* | *G. ogac* | *G. chalcogrammus* | 34,681.2 | 296.2 | 226.2 | 0.134 | 0.0009 |
| *G. macrocephalus* | *G. ogac* | Lofoten (s) | 23,711.0 | 297.2 | 165.5 | 0.285 | 0.0000 |
| *G. macrocephalus* | *G. ogac* | Suffolk | 34,836.4 | 414.9 | 232.1 | 0.283 | 0.0000 |
| *G. macrocephalus* | *G. ogac* | Kiel Bight | 35,330.2 | 415.8 | 237.6 | 0.273 | 0.0000 |
| *G. macrocephalus* | *G. ogac* | Bornholm Basin | 32,867.1 | 423.3 | 210.6 | 0.336 | 0.0000 |
| *G. macrocephalus* | *G. ogac* | Iceland (s) | 31,577.5 | 395.5 | 197.0 | 0.335 | 0.0000 |
| *G. macrocephalus* | *G. ogac* | Møre (s) | 33,823.1 | 438.9 | 213.1 | 0.346 | 0.0000 |
| *G. macrocephalus* | *G. ogac* | Lofoten (m) | 31,503.6 | 410.2 | 188.9 | <u>0.369</u> | 0.0000 |
| *G. macrocephalus* | *G. ogac* | Newfoundland (s) | 35,366.2 | 449.3 | 220.4 | 0.342 | 0.0000 |
| *G. macrocephalus* | *G. ogac* | Iceland (m) | 35,865.7 | 458.8 | 226.7 | 0.339 | 0.0000 |
| *G. macrocephalus* | *G. ogac* | Møre (m) | 35,777.5 | 454.6 | 222.2 | 0.343 | 0.0000 |
| *G. macrocephalus* | *G. ogac* | Newfoundland (m) | 35,303.2 | 448.4 | 221.6 | 0.338 | 0.0000 |
| *G. macrocephalus* | *G. ogac* | Labrador | 34,386.4 | 432.6 | 215.0 | 0.336 | 0.0000 |
| Iceland (m) | Lofoten (s) | *G. chalcogrammus* | 16,586.6 | 518.0 | 464.1 | 0.055 | 0.0767 |
| Iceland (m) | Suffolk | *G. chalcogrammus* | 23,884.5 | 730.8 | 596.9 | 0.101 | 0.0016 |
| Iceland (m) | Kiel Bight | *G. chalcogrammus* | 24,186.9 | 711.4 | 638.1 | 0.054 | 0.0294 |
| Iceland (m) | Iceland (s) | *G. chalcogrammus* | 25,400.4 | 252.7 | 231.1 | 0.045 | 0.0345 |
| Iceland (m) | Møre (s) | *G. chalcogrammus* | 27,557.4 | 291.4 | 257.2 | 0.062 | 0.0308 |
| Iceland (m) | Newfoundland (s) | *G. chalcogrammus* | 28,062.6 | 319.1 | 287.4 | 0.052 | 0.0677 |
| Iceland (m) | Møre (m) | *G. chalcogrammus* | 28,871.7 | 307.4 | 277.0 | 0.052 | 0.0124 |
| Iceland (m) | Newfoundland (m) | *G. chalcogrammus* | 28,164.5 | 311.6 | 273.2 | 0.066 | 0.0050 |
| Iceland (m) | Labrador | *G. chalcogrammus* | 27,617.1 | 329.3 | 282.9 | 0.076 | 0.0073 |
| Lofoten (m) | Bornholm Basin | Lofoten (s) | 3,652.4 | 704.3 | 604.1 | 0.077 | 0.0066 |
| Lofoten (m) | Iceland (s) | Lofoten (s) | 3,520.5 | 631.9 | 533.6 | 0.084 | 0.0142 |
| Lofoten (m) | Møre (s) | Lofoten (s) | 3,810.3 | 656.3 | 605.3 | 0.040 | 0.0724 |
| Labrador | Newfoundland (s) | Lofoten (s) | 3,973.7 | 730.0 | 680.4 | 0.035 | 0.0978 |
| Lofoten (m) | Iceland (m) | Lofoten (s) | 3,886.5 | 678.8 | 629.7 | 0.038 | 0.0764 |
| Lofoten (m) | Møre (m) | Lofoten (s) | 3,890.5 | 683.1 | 633.7 | 0.037 | 0.0395 |
| Labrador | Newfoundland (m) | Lofoten (s) | 4,015.7 | 733.7 | 661.9 | 0.051 | 0.0681 |
| Lofoten (m) | Bornholm Basin | Suffolk | 4,839.8 | 962.0 | 811.7 | 0.085 | 0.0087 |
| Lofoten (m) | Iceland (s) | Suffolk | 4,586.9 | 824.6 | 742.6 | 0.052 | 0.0313 |
| Labrador | Newfoundland (s) | Suffolk | 5,323.9 | 999.3 | 940.4 | 0.030 | 0.0808 |
| Lofoten (m) | Møre (m) | Suffolk | 5,194.6 | 919.9 | 871.4 | 0.027 | 0.0930 |
| Labrador | Newfoundland (m) | Suffolk | 5,359.6 | 987.9 | 908.7 | 0.042 | 0.0809 |
| Lofoten (m) | Bornholm Basin | Kiel Bight | 4,878.6 | 974.6 | 826.1 | 0.083 | 0.0051 |
| Lofoten (m) | Iceland (s) | Kiel Bight | 4,620.7 | 840.4 | 744.6 | 0.060 | 0.0127 |
| Lofoten (m) | Møre (s) | Kiel Bight | 5,100.4 | 893.7 | 840.1 | 0.031 | 0.0948 |
| Labrador | Newfoundland (s) | Kiel Bight | 5,388.7 | 1,016.3 | 948.1 | 0.035 | 0.0776 |
| Lofoten (m) | Møre (m) | Kiel Bight | 5,237.2 | 930.3 | 881.9 | 0.027 | 0.0946 |

**Supplementary Table 18 (continued):** $D_{\mathrm{BBAA}}$-statistics for *Gadus morhua* populations and outgroups within the supergene region on LG 12.

| P1 | P2 | P3 | $C_{\mathrm{BBAA}}$ | $C_{\mathrm{ABBA}}$ | $C_{\mathrm{BABA}}$ | $D_{\mathrm{BBAA}}$ | $p$ |
|---|---|---|---|---|---|---|---|
| Labrador | Newfoundland (m) | Kiel Bight | 5,408.1 | 1,017.8 | 910.1 | 0.056 | 0.0313 |
| Møre (m) | Møre (s) | Bornholm Basin | 1,590.3 | 1,549.4 | 1,383.9 | 0.056 | 0.0231 |
| Newfoundland (s) | Newfoundland (m) | Bornholm Basin | 1,701.7 | 1,481.1 | 1,361.7 | 0.042 | 0.0328 |
| Lofoten (m) | Iceland (s) | Newfoundland (m) | 1,506.7 | 1,308.6 | 1,188.6 | 0.048 | 0.0028 |
| Iceland (m) | Møre (s) | Lofoten (m) | 1,531.2 | 1,520.1 | 1,408.7 | 0.038 | 0.0382 |
| Lofoten (m) | Møre (s) | Newfoundland (s) | 1,804.9 | 1,383.2 | 1,325.4 | 0.021 | 0.0511 |
| Iceland (m) | Møre (s) | Møre (m) | 1,564.0 | 1,557.7 | 1,474.1 | 0.028 | 0.0635 |
| Newfoundland (s) | Newfoundland (m) | Møre (s) | 1,671.3 | 1,571.4 | 1,405.3 | 0.056 | 0.0028 |
| Bornholm Basin | Lofoten (m) | Møre (m) | 1,530.5 | 1,521.2 | 1,414.7 | 0.036 | 0.0315 |
| Newfoundland (s) | Newfoundland (m) | Lofoten (m) | 1,697.5 | 1,469.7 | 1,383.0 | 0.030 | 0.0413 |
| Newfoundland (s) | Labrador | Lofoten (m) | 1,637.4 | 1,532.4 | 1,440.6 | 0.031 | 0.0591 |
| Lofoten (m) | Møre (m) | Newfoundland (s) | 1,767.5 | 1,458.1 | 1,375.1 | 0.029 | 0.0200 |
| Newfoundland (s) | Newfoundland (m) | Iceland (m) | 1,743.8 | 1,583.1 | 1,440.8 | 0.047 | 0.0052 |
| Lofoten (m) | Iceland (m) | Labrador | 1,719.6 | 1,505.2 | 1,426.9 | 0.027 | 0.0384 |

**Supplementary Table 19:** $D_{\mathrm{fix}}$-statistics for *Gadus morhua* populations and outgroups within the supergene region on LG 12.
The $D_{\mathrm{fix}}$ version of the $D$-statistic[35] was calculated as for Supplementary Table 10, except that only sites from the supergene region on LG 12 were used and the input tree was the one inferred for the supergene on LG 12 under the multi-species coalescent model (Fig. 5e). The highest $D$ values are underlined.

| P1 | P2 | P3 | $C_{\mathrm{BBAA}}$ | $C_{\mathrm{ABBA}}$ | $C_{\mathrm{BABA}}$ | $D_{\mathrm{BBAA}}$ | $p$ |
|---|---|---|---|---|---|---|---|
| Bornholm Basin | *G. chalcogrammus* | *G. macrocephalus* | 13,979.6 | 2,751.8 | 2,566.6 | 0.035 | 0.0179 |
| Bornholm Basin | Lofoten (s) | *G. macrocephalus* | 23,094.0 | 363.2 | 293.2 | 0.107 | 0.0094 |
| Bornholm Basin | Suffolk | *G. macrocephalus* | 33,148.2 | 491.0 | 391.6 | 0.113 | 0.0005 |
| Bornholm Basin | Kiel Bight | *G. macrocephalus* | 33,505.7 | 485.5 | 408.0 | 0.087 | 0.0026 |
| *G. macrocephalus* | *G. ogac* | *G. chalcogrammus* | 34,681.2 | 296.2 | 226.2 | 0.134 | 0.0009 |
| *G. macrocephalus* | *G. ogac* | Lofoten (s) | 23,711.0 | 297.2 | 165.5 | 0.285 | 0.0000 |
| *G. macrocephalus* | *G. ogac* | Suffolk | 34,836.4 | 414.9 | 232.1 | 0.283 | 0.0000 |
| *G. macrocephalus* | *G. ogac* | Kiel Bight | 35,330.2 | 415.8 | 237.6 | 0.273 | 0.0000 |
| *G. macrocephalus* | *G. ogac* | Bornholm Basin | 32,867.1 | 423.3 | 210.6 | 0.336 | 0.0000 |
| *G. macrocephalus* | *G. ogac* | Iceland (s) | 31,577.5 | 395.5 | 197.0 | 0.335 | 0.0000 |
| *G. macrocephalus* | *G. ogac* | Møre (s) | 33,823.1 | 438.9 | 213.1 | 0.346 | 0.0000 |
| *G. macrocephalus* | *G. ogac* | Lofoten (m) | 31,503.6 | 410.2 | 188.9 | <u>0.369</u> | 0.0000 |
| *G. macrocephalus* | *G. ogac* | Newfoundland (s) | 35,366.2 | 449.3 | 220.4 | 0.342 | 0.0000 |
| *G. macrocephalus* | *G. ogac* | Iceland (m) | 35,865.7 | 458.8 | 226.7 | 0.339 | 0.0000 |
| *G. macrocephalus* | *G. ogac* | Møre (m) | 35,777.5 | 454.6 | 222.2 | 0.343 | 0.0000 |
| *G. macrocephalus* | *G. ogac* | Newfoundland (m) | 35,303.2 | 448.4 | 221.6 | 0.338 | 0.0000 |
| *G. macrocephalus* | *G. ogac* | Labrador | 34,386.4 | 432.6 | 215.0 | 0.336 | 0.0000 |
| Iceland (m) | Lofoten (s) | *G. chalcogrammus* | 16,586.6 | 518.0 | 464.1 | 0.055 | 0.0767 |
| Iceland (m) | Suffolk | *G. chalcogrammus* | 23,884.5 | 730.8 | 596.9 | 0.101 | 0.0016 |
| Iceland (m) | Kiel Bight | *G. chalcogrammus* | 24,186.9 | 711.4 | 638.1 | 0.054 | 0.0294 |

**Supplementary Table 19 (continued):** $D_{\text{fix}}$-statistics for *Gadus morhua* populations and outgroups within the supergene region on LG 12.

| P1 | P2 | P3 | $C_{\text{BBAA}}$ | $C_{\text{ABBA}}$ | $C_{\text{BABA}}$ | $D_{\text{BBAA}}$ | $p$ |
|---|---|---|---|---|---|---|---|
| Iceland (m) | Iceland (s) | *G. chalcogrammus* | 25,400.4 | 252.7 | 231.1 | 0.045 | 0.0345 |
| Iceland (m) | Møre (s) | *G. chalcogrammus* | 27,557.4 | 291.4 | 257.2 | 0.062 | 0.0308 |
| Iceland (m) | Newfoundland (s) | *G. chalcogrammus* | 28,062.6 | 319.1 | 287.4 | 0.052 | 0.0677 |
| Iceland (m) | Møre (m) | *G. chalcogrammus* | 28,871.7 | 307.4 | 277.0 | 0.052 | 0.0124 |
| Iceland (m) | Newfoundland (m) | *G. chalcogrammus* | 28,164.5 | 311.6 | 273.2 | 0.066 | 0.0050 |
| Iceland (m) | Labrador | *G. chalcogrammus* | 27,617.1 | 329.3 | 282.9 | 0.076 | 0.0073 |
| Lofoten (m) | Bornholm Basin | Lofoten (s) | 3,652.4 | 704.3 | 604.1 | 0.077 | 0.0066 |
| Lofoten (m) | Iceland (s) | Lofoten (s) | 3,520.5 | 631.9 | 533.6 | 0.084 | 0.0142 |
| Lofoten (m) | Møre (s) | Lofoten (s) | 3,810.3 | 656.3 | 605.3 | 0.040 | 0.0724 |
| Labrador | Newfoundland (s) | Lofoten (s) | 3,973.7 | 730.0 | 680.4 | 0.035 | 0.0978 |
| Lofoten (m) | Iceland (m) | Lofoten (s) | 3,886.5 | 678.8 | 629.7 | 0.038 | 0.0764 |
| Lofoten (m) | Møre (m) | Lofoten (s) | 3,890.5 | 683.1 | 633.7 | 0.037 | 0.0395 |
| Labrador | Newfoundland (m) | Lofoten (s) | 4,015.7 | 733.7 | 661.9 | 0.051 | 0.0681 |
| Lofoten (m) | Bornholm Basin | Suffolk | 4,839.8 | 962.0 | 811.7 | 0.085 | 0.0087 |
| Lofoten (m) | Iceland (s) | Suffolk | 4,586.9 | 824.6 | 742.6 | 0.052 | 0.0313 |
| Labrador | Newfoundland (s) | Suffolk | 5,323.9 | 999.3 | 940.4 | 0.030 | 0.0808 |
| Lofoten (m) | Møre (m) | Suffolk | 5,194.6 | 919.9 | 871.4 | 0.027 | 0.0930 |
| Labrador | Newfoundland (m) | Suffolk | 5,359.6 | 987.9 | 908.7 | 0.042 | 0.0809 |
| Lofoten (m) | Bornholm Basin | Kiel Bight | 4,878.6 | 974.6 | 826.1 | 0.083 | 0.0051 |
| Lofoten (m) | Iceland (s) | Kiel Bight | 4,620.7 | 840.4 | 744.6 | 0.060 | 0.0127 |
| Lofoten (m) | Møre (s) | Kiel Bight | 5,100.4 | 893.7 | 840.1 | 0.031 | 0.0948 |
| Labrador | Newfoundland (s) | Kiel Bight | 5,388.7 | 1,016.3 | 948.1 | 0.035 | 0.0776 |
| Lofoten (m) | Møre (m) | Kiel Bight | 5,237.2 | 930.3 | 881.9 | 0.027 | 0.0946 |
| Labrador | Newfoundland (m) | Kiel Bight | 5,408.1 | 1,017.8 | 910.1 | 0.056 | 0.0313 |
| Newfoundland (s) | Iceland (s) | Bornholm Basin | 1,300.2 | 1,525.4 | 1,216.0 | 0.113 | 0.0036 |
| Newfoundland (s) | Møre (s) | Bornholm Basin | 1,399.1 | 1,783.7 | 1,320.9 | 0.149 | 0.0000 |
| Newfoundland (s) | Lofoten (m) | Bornholm Basin | 1,342.9 | 1,735.7 | 1,334.2 | 0.131 | 0.0005 |
| Newfoundland (s) | Iceland (m) | Bornholm Basin | 1,439.2 | 1,772.6 | 1,388.4 | 0.122 | 0.0016 |
| Newfoundland (s) | Møre (m) | Bornholm Basin | 1,495.2 | 1,707.4 | 1,408.6 | 0.096 | 0.0103 |
| Newfoundland (s) | Newfoundland (m) | Bornholm Basin | 1,701.7 | 1,481.1 | 1,361.7 | 0.042 | 0.0328 |
| Newfoundland (s) | Møre (s) | Iceland (s) | 1,213.9 | 1,635.0 | 1,248.8 | 0.134 | 0.0000 |
| Newfoundland (s) | Lofoten (m) | Iceland (s) | 1,206.7 | 1,572.0 | 1,273.6 | 0.105 | 0.0010 |
| Newfoundland (s) | Iceland (m) | Iceland (s) | 1,248.5 | 1,647.5 | 1,298.5 | 0.118 | 0.0006 |
| Newfoundland (s) | Møre (m) | Iceland (s) | 1,275.6 | 1,620.3 | 1,283.5 | 0.116 | 0.0022 |
| Newfoundland (s) | Newfoundland (m) | Iceland (s) | 1,509.4 | 1,395.0 | 1,292.7 | 0.038 | 0.0173 |
| Lofoten (m) | Møre (s) | Newfoundland (s) | 1,804.9 | 1,383.2 | 1,325.4 | 0.021 | 0.0511 |
| Lofoten (m) | Møre (s) | Iceland (m) | 1,520.1 | 1,531.2 | 1,408.7 | 0.042 | 0.0311 |
| Newfoundland (s) | Møre (m) | Møre (s) | 1,417.3 | 1,844.9 | 1,398.8 | 0.138 | 0.0001 |
| Newfoundland (s) | Newfoundland (m) | Møre (s) | 1,671.3 | 1,571.4 | 1,405.3 | 0.056 | 0.0028 |
| Newfoundland (s) | Møre (m) | Lofoten (m) | 1,458.1 | 1,767.5 | 1,375.1 | 0.125 | 0.0001 |
| Newfoundland (s) | Newfoundland (m) | Lofoten (m) | 1,697.5 | 1,469.7 | 1,383.0 | 0.030 | 0.0413 |
| Newfoundland (s) | Labrador | Lofoten (m) | 1,637.4 | 1,532.4 | 1,440.6 | 0.031 | 0.0591 |
| Labrador | Newfoundland (s) | Newfoundland (m) | 1,520.4 | 1,646.6 | 1,572.4 | 0.023 | 0.0919 |
| Newfoundland (s) | Møre (m) | Iceland (m) | 1,495.7 | 1,854.2 | 1,461.6 | 0.118 | 0.0006 |
| Newfoundland (s) | Newfoundland (m) | Iceland (m) | 1,743.8 | 1,583.1 | 1,440.8 | 0.047 | 0.0052 |
| Lofoten (m) | Iceland (m) | Labrador | 1,719.6 | 1,505.2 | 1,426.9 | 0.027 | 0.0384 |

**Supplementary Table 20:** Predicted genes within the region of the supergene on LG 12 with evidence for double crossover (positions 7,478,537 bp to 7,752,994 bp).
Gene IDs refer to the gadMor2 assembly annotation[10]. BLASTN searches suggest that gene ID 00027785 corresponds to vitellogenin type A, while both IDs 00027782 and 00027783 correspond to vitellogenin type B. As vitellogenin B

| Beginning | End | Strand | Gene ID | Predicted gene | GO terms |
|---|---|---|---|---|---|
| 7,468,870 | 7,485,996 | − | 00027772 | TTLL7 (*Homo sapiens*) | GO:0006464 |
| 7,493,441 | 7,504,363 | + | 00027774 | PRKACA (*Sus scrofa*) | GO:0004672, GO:0004674, GO:0005524, GO:0006468, GO:0016772 |
| 7,504,544 | 7,507,926 | + | 00027775 | SAMD13 (*Homo sapiens*) | GO:0005515 |
| 7,507,198 | 7,513,354 | − | 00027776 | UOX (*Oryctolagus cuniculus*) | — |
| 7,512,454 | 7,516,738 | + | 00027777 | DNASE2B (*Homo sapiens*) | GO:0004531, GO:0006259 |
| 7,517,560 | 7,520,918 | + | 00027779 | rpf1 (*Danio rerio*) | — |
| 7,526,726 | 7,530,804 | − | 00027781 | Ctbs (*Mus musculus*) | GO:0004553, GO:0004568, GO:0005975, GO:0006032 |
| 7,534,022 | 7,547,615 | + | 00027782 | vtg1 (*Oncorhynchus mykiss*) | GO:0005319, GO:0006869 |
| 7,551,466 | 7,559,692 | − | 00027783 | Vitellogenin (*Fundulus heteroclitus*) | GO:0005319, GO:0006869 |
| 7,565,225 | 7,575,883 | + | 00027785 | vtg1 (*Oncorhynchus mykiss*) | GO:0005319, GO:0006869 |
| 7,575,985 | 7,583,560 | − | 00027786 | ssx2ip (*Oryzias latipes*) | — |
| 7,604,691 | 7,610,866 | − | 00027788 | Lpar3 (*Mus musculus*) | GO:0004930, GO:0007186, GO:0016021, GO:0070915 |
| 7,613,633 | 7,625,764 | − | 00027790 | MCOLN2 (*Homo sapiens*) | — |
| 7,627,573 | 7,633,411 | − | 00027791 | CLCC1 (*Bos taurus*) | — |
| 7,633,462 | 7,641,637 | − | 00027793 | FTSJ3 (*Gallus gallus*) | GO:0001510, GO:0005634, GO:0006364, GO:0008168, GO:0008649, GO:0031167, GO:0032259 |
| 7,641,843 | 7,644,988 | − | 00027795 | Hrasls (*Mus musculus*) | — |
| 7,645,411 | 7,660,397 | + | 00027796 | Mfn1 (*Rattus norvegicus*) | GO:0003924, GO:0005525, GO:0005741, GO:0008053, GO:0016021 |
| 7,660,315 | 7,678,716 | − | 00027799 | GNB4 (*Homo sapiens*) | GO:0005515, GO:0016491, GO:0051537, GO:0055114 |
| 7,684,579 | 7,693,849 | + | 00027802 | ACTL6A (*Homo sapiens*) | — |
| 7,692,544 | 7,696,524 | − | 00027803 | Mrpl47 (*Mus musculus*) | GO:0003735, GO:0005761, GO:0006412 |
| 7,696,509 | 7,699,595 | + | 00027804 | NDUFB5 (*Bos taurus*) | — |
| 7,700,430 | 7,730,778 | + | 00027805 | usp13 (*Danio rerio*) | GO:0005515, GO:0006511, GO:0008270, GO:0036459 |
| 7,734,260 | 7,769,915 | − | 00027810 | PEX5L (*Homo sapiens*) | GO:0005515 |
| 7,823,234 | 7,825,557 | − | 00027816 | Cpn2 (*Mus musculus*) | GO:0005515 |

**Supplementary Table 21:** Genomic signatures of extreme bottlenecks.
To assess how long signatures of brief but extreme bottlenecks can remain in the genomes of the affected population, we simulated genetic data with msprime, under a model of a single panmictic population that experienced an extreme bottleneck in the past during which the population size was reduced to a single sequence for a duration of 20 generations. At the end of the simulation, one diploid genome was sampled from the population and the proportion of the diploid genome coalescing in (or before) the bottleneck, $p_{\text{bottleneck}}$, was quantified. We further quantified the genetic diversity of the simulated genome, $\pi_{\text{bottleneck}}$, as well as the genetic diversity of a second genome simulated with the same settings except the bottleneck, $\pi_{\text{no bottleneck}}$. The simulated linkage group had a length of 20 Mb and the generation time was assumed to be 10 years. We repeated the simulation with five different values for the diploid population size, $N_{\text{e}}$, (outside the bottleneck) between 1,000 and 100,000 and with five different values for the age of the bottleneck, $t_{\text{bottleneck}}$, between 3,000 years and 3,000,000 years. Reported are the mean values across 20 simulations followed by standard deviations in parentheses.

| $N_{\text{e}}$ | $t_{\text{bottleneck}}$ | $p_{\text{bottleneck}}$ | $\pi_{\text{bottleneck}}$ ($\times 10^3$) | $\pi_{\text{no bottleneck}}$ ($\times 10^3$) |
|---:|---:|---:|---:|---:|
| 1,000 | 3,000 | 0.89 (0.06) | 0.004 (0.001) | 0.030 (0.004) |
| 1,000 | 10,000 | 0.61 (0.07) | 0.012 (0.002) | 0.031 (0.003) |
| 1,000 | 30,000 | 0.22 (0.04) | 0.025 (0.003) | 0.032 (0.004) |
| 1,000 | 100,000 | 0.01 (0.00) | 0.031 (0.003) | 0.031 (0.003) |
| 1,000 | 300,000 | 0.00 (0.00) | 0.032 (0.003) | 0.030 (0.004) |
| 1,000 | 1,000,000 | 0.00 (0.00) | 0.032 (0.003) | 0.031 (0.004) |
| 1,000 | 3,000,000 | 0.00 (0.00) | 0.031 (0.004) | 0.030 (0.005) |
| 3,000 | 3,000 | 0.96 (0.05) | 0.005 (0.001) | 0.094 (0.009) |
| 3,000 | 10,000 | 0.85 (0.05) | 0.015 (0.001) | 0.093 (0.008) |
| 3,000 | 30,000 | 0.59 (0.05) | 0.038 (0.003) | 0.090 (0.011) |
| 3,000 | 100,000 | 0.19 (0.01) | 0.078 (0.005) | 0.095 (0.011) |
| 3,000 | 300,000 | 0.01 (0.00) | 0.095 (0.005) | 0.093 (0.010) |
| 3,000 | 1,000,000 | 0.00 (0.00) | 0.095 (0.005) | 0.096 (0.007) |
| 3,000 | 3,000,000 | 0.00 (0.00) | 0.097 (0.005) | 0.092 (0.009) |
| 10,000 | 3,000 | 0.98 (0.03) | 0.005 (0.000) | 0.318 (0.011) |
| 10,000 | 10,000 | 0.95 (0.02) | 0.016 (0.001) | 0.307 (0.011) |
| 10,000 | 30,000 | 0.86 (0.02) | 0.044 (0.002) | 0.314 (0.013) |
| 10,000 | 100,000 | 0.61 (0.02) | 0.123 (0.005) | 0.312 (0.018) |
| 10,000 | 300,000 | 0.22 (0.01) | 0.242 (0.010) | 0.304 (0.035) |
| 10,000 | 1,000,000 | 0.01 (0.00) | 0.315 (0.011) | 0.314 (0.013) |
| 10,000 | 3,000,000 | 0.00 (0.00) | 0.316 (0.016) | 0.315 (0.015) |
| 30,000 | 3,000 | 0.99 (0.03) | 0.005 (0.001) | 0.946 (0.024) |
| 30,000 | 10,000 | 0.99 (0.01) | 0.016 (0.001) | 0.953 (0.018) |
| 30,000 | 30,000 | 0.96 (0.01) | 0.047 (0.002) | 0.939 (0.038) |
| 30,000 | 100,000 | 0.84 (0.04) | 0.144 (0.005) | 0.940 (0.019) |
| 30,000 | 300,000 | 0.61 (0.02) | 0.369 (0.007) | 0.943 (0.015) |
| 30,000 | 1,000,000 | 0.19 (0.01) | 0.762 (0.017) | 0.944 (0.018) |
| 30,000 | 3,000,000 | 0.01 (0.00) | 0.942 (0.023) | 0.939 (0.027) |
| 100,000 | 3,000 | 1.00 (0.00) | 0.005 (0.000) | 3.122 (0.044) |
| 100,000 | 10,000 | 1.00 (0.01) | 0.016 (0.001) | 3.132 (0.043) |
| 100,000 | 30,000 | 0.98 (0.01) | 0.047 (0.002) | 3.139 (0.041) |
| 100,000 | 100,000 | 0.95 (0.01) | 0.152 (0.003) | 3.124 (0.037) |

**Supplementary Table 21 (continued):** Consequences of extreme bottlenecks.

| $N_e$ | $t_{bottleneck}$ | $p_{bottleneck}$ | $\pi_{bottleneck}$ ($\times 10^3$) | $\pi_{no\ bottleneck}$ ($\times 10^3$) |
|---|---|---|---|---|
| 100,000 | 300,000 | 0.86 (0.02) | 0.437 (0.008) | 3.138 (0.051) |
| 100,000 | 1,000,000 | 0.61 (0.01) | 1.236 (0.013) | 3.122 (0.035) |
| 100,000 | 3,000,000 | 0.22 (0.00) | 2.444 (0.035) | 3.131 (0.053) |

**Supplementary Table 22:** *Gadus morhua* specimens used in analyses of linkage disequilibrium. Genomes of 100 *Gadus morhua* specimens were used exclusively in analyses of linkage disequilibrium and selected so that at each of the four LGs with supergenes, both the ancestral and the derived orientation of the supergene were found in exactly 50 specimens.

| Specimen ID | ENA accession(s) | Population | LG 1 | LG 2 | LG 7 | LG 12 |
|---|---|---|---|---|---|---|
| AVE1403Z09 | NA | Møre | derived | ancestral | derived | ancestral |
| AVE1403Z11 | ERR5321906 | Møre | derived | ancestral | derived | ancestral |
| AVE1403Z17 | NA | Møre | derived | ancestral | derived | ancestral |
| AVE1403Z20 | NA | Møre | derived | ancestral | derived | ancestral |
| AVE1403Z27 | NA | Møre | derived | ancestral | derived | ancestral |
| AVE1403Z29 | NA | Møre | derived | ancestral | derived | ancestral |
| AVE1403Z34 | NA | Møre | derived | ancestral | derived | ancestral |
| AVE1403Z35 | NA | Møre | derived | ancestral | derived | ancestral |
| CEL1002Z01 | NA | Celtic Sea | ancestral | derived | ancestral | derived |
| CEL1002Z02 | NA | Celtic Sea | ancestral | derived | ancestral | derived |
| CEL1002Z08 | NA | Celtic Sea | ancestral | derived | ancestral | derived |
| CEL1002Z10 | NA | Celtic Sea | ancestral | derived | ancestral | derived |
| CEL1002Z12 | NA | Celtic Sea | ancestral | derived | ancestral | derived |
| CEL1002Z14 | NA | Celtic Sea | ancestral | derived | ancestral | derived |
| CEL1002Z15 | NA | Celtic Sea | ancestral | derived | ancestral | derived |
| CEL1002Z16 | NA | Celtic Sea | ancestral | derived | ancestral | derived |
| CEL1002Z17 | NA | Celtic Sea | ancestral | derived | ancestral | derived |
| CEL1002Z24 | NA | Celtic Sea | ancestral | derived | ancestral | derived |
| ENG0701Z01 | NA | English Channel | ancestral | derived | ancestral | derived |
| ENG0701Z02 | NA | English Channel | ancestral | derived | ancestral | derived |
| ENG0701Z03 | NA | English Channel | ancestral | derived | ancestral | derived |
| ENG0701Z05 | NA | English Channel | ancestral | derived | ancestral | derived |
| ENG0701Z07 | NA | English Channel | ancestral | derived | ancestral | derived |
| ENG0701Z09 | NA | English Channel | ancestral | derived | ancestral | derived |
| ENG0701Z16 | NA | English Channel | ancestral | derived | ancestral | derived |
| ENG0701Z20 | NA | English Channel | ancestral | derived | ancestral | derived |
| ENG0701Z22 | NA | English Channel | ancestral | derived | ancestral | derived |
| ENG0902Z01 | NA | English Channel | ancestral | derived | ancestral | derived |
| ENG0902Z03 | NA | English Channel | ancestral | derived | ancestral | derived |
| ENG0902Z04 | NA | English Channel | ancestral | derived | ancestral | derived |
| ICC0304Z14 | NA | Iceland | derived | ancestral | derived | ancestral |
| ICC0304Z15 | NA | Iceland | derived | ancestral | derived | ancestral |
| ICC0304Z16 | NA | Iceland | derived | ancestral | derived | ancestral |
| ICO0304Z01 | NA | Iceland | derived | ancestral | derived | ancestral |
| ICO0304Z10 | NA | Iceland | derived | ancestral | derived | ancestral |

**Supplementary Table 22 (continued):** *Gadus morhua* specimens used in analyses of linkage disequilibrium.

| Specimen ID | ENA accession(s) | Population | LG 1 | LG 2 | LG 7 | LG 12 |
|---|---|---|---|---|---|---|
| ICO0304Z11 | NA | Iceland | derived | ancestral | derived | ancestral |
| ICO0304Z14 | NA | Iceland | derived | ancestral | derived | ancestral |
| ICO0304Z15 | NA | Iceland | derived | ancestral | derived | ancestral |
| ICO0304Z16 | NA | Iceland | derived | ancestral | derived | ancestral |
| ICO0304Z19 | NA | Iceland | derived | ancestral | derived | ancestral |
| ICO0304Z20 | ERR5321899 | Iceland | derived | ancestral | derived | ancestral |
| ICO0304Z21 | NA | Iceland | derived | ancestral | derived | ancestral |
| IRI0802Z01 | NA | Irish Sea | ancestral | derived | ancestral | derived |
| IRI0902Z02 | NA | Irish Sea | ancestral | derived | ancestral | derived |
| IRI0902Z06 | NA | Irish Sea | ancestral | derived | ancestral | derived |
| IRI0903Z01 | NA | Irish Sea | ancestral | derived | ancestral | derived |
| IRI0903Z02 | NA | Irish Sea | ancestral | derived | ancestral | derived |
| IRI0903Z03 | NA | Irish Sea | ancestral | derived | ancestral | derived |
| IRI0903Z04 | NA | Irish Sea | ancestral | derived | ancestral | derived |
| IRI1002Z03 | NA | Irish Sea | ancestral | derived | ancestral | derived |
| IRI1003Z05 | NA | Irish Sea | ancestral | derived | ancestral | derived |
| IRI1003Z07 | NA | Irish Sea | ancestral | derived | ancestral | derived |
| IRI1003Z12 | NA | Irish Sea | ancestral | derived | ancestral | derived |
| KIE1103Z20 | ERR2850485 | Kiel Bight | ancestral | derived | ancestral | derived |
| LOF1103Z04 | ERR4851787,88 | Lofoten | derived | ancestral | derived | ancestral |
| LOF1103Z06 | ERR4851791,92 | Lofoten | derived | ancestral | derived | ancestral |
| LOF1103Z08 | ERR4851795,96 | Lofoten | derived | ancestral | derived | ancestral |
| LOF1103Z10 | ERR4851799,800 | Lofoten | derived | ancestral | derived | ancestral |
| LOF1103Z11 | ERR4851801,02 | Lofoten | derived | ancestral | derived | ancestral |
| LOF1103Z17 | ERR4851813,14 | Lofoten | derived | ancestral | derived | ancestral |
| LOF1103Z20 | ERR4851819,20 | Lofoten | derived | ancestral | derived | ancestral |
| LOF1103Z24 | ERR4851827,28 | Lofoten | derived | ancestral | derived | ancestral |
| LOF1403Z02 | ERR4851830 | Lofoten | derived | ancestral | derived | ancestral |
| LOF1403Z04 | ERR4851832 | Lofoten | derived | ancestral | derived | ancestral |
| LOF1403Z06 | ERR4851834 | Lofoten | derived | ancestral | derived | ancestral |
| LOF1403Z07 | ERR4851835 | Lofoten | derived | ancestral | derived | ancestral |
| LOF1403Z08 | ERR4851836 | Lofoten | derived | ancestral | derived | ancestral |
| LOF1403Z09 | ERR4851837 | Lofoten | derived | ancestral | derived | ancestral |
| LOF1403Z11 | ERR4851839 | Lofoten | derived | ancestral | derived | ancestral |
| LOF1403Z12 | ERR4851840 | Lofoten | derived | ancestral | derived | ancestral |
| LOF1403Z14 | ERR4851842 | Lofoten | derived | ancestral | derived | ancestral |
| LOF1403Z17 | ERR4851845 | Lofoten | derived | ancestral | derived | ancestral |
| LOF1403Z18 | ERR4851846 | Lofoten | derived | ancestral | derived | ancestral |
| LOF1403Z24 | ERR4851852 | Lofoten | derived | ancestral | derived | ancestral |
| LOW1503Z01 | ERR2850495-98 | Suffolk | ancestral | derived | ancestral | derived |
| LOW1503Z04 | ERR2850507-10 | Suffolk | ancestral | derived | ancestral | derived |
| LOW1503Z07 | ERR2850519-22 | Suffolk | ancestral | derived | ancestral | derived |
| LOW1503Z08 | ERR2850523-26 | Suffolk | ancestral | derived | ancestral | derived |
| LOW1504Z02 | ERR2850559-62 | Suffolk | ancestral | derived | ancestral | derived |
| LOW1504Z05 | ERR2850571-74 | Suffolk | ancestral | derived | ancestral | derived |
| LOW1504Z06 | ERR2850575-78 | Suffolk | ancestral | derived | ancestral | derived |
| LOW1504Z07 | ERR2850579-82 | Suffolk | ancestral | derived | ancestral | derived |

**Supplementary Table 22 (continued):** *Gadus morhua* specimens used in analyses of linkage disequilibrium.

| Specimen ID | ENA accession(s) | Population | LG 1 | LG 2 | LG 7 | LG 12 |
|---|---|---|---|---|---|---|
| LOW1504Z08 | ERR2850583-86 | Suffolk | ancestral | derived | ancestral | derived |
| SCA0601Z04 | NA | North Yorkshire | ancestral | derived | ancestral | derived |
| SCA0601Z12 | NA | North Yorkshire | ancestral | derived | ancestral | derived |
| SOR1403Z01 | NA | Finnmark | derived | ancestral | derived | ancestral |
| SOR1403Z02 | NA | Finnmark | derived | ancestral | derived | ancestral |
| SOR1403Z07 | NA | Finnmark | derived | ancestral | derived | ancestral |
| SOR1404Z02 | NA | Finnmark | derived | ancestral | derived | ancestral |
| SOR1404Z03 | NA | Finnmark | derived | ancestral | derived | ancestral |
| SOR1404Z05 | NA | Finnmark | derived | ancestral | derived | ancestral |
| SOR1404Z06 | NA | Finnmark | derived | ancestral | derived | ancestral |
| SOR1404Z09 | NA | Finnmark | derived | ancestral | derived | ancestral |
| SOR1404Z10 | NA | Finnmark | derived | ancestral | derived | ancestral |
| SOR1404Z12 | NA | Finnmark | derived | ancestral | derived | ancestral |
| TVE1105Z09 | ERR2850674 | Agder | ancestral | derived | ancestral | derived |
| TVE1105Z19 | ERR2850683 | Agder | ancestral | derived | ancestral | derived |
| TVE1205Z01 | ERR2850689 | Agder | ancestral | derived | ancestral | derived |
| TVE1205Z10 | ERR2850698 | Agder | ancestral | derived | ancestral | derived |
| TVE1205Z13 | ERR2850701 | Agder | ancestral | derived | ancestral | derived |

### Supplementary References


1. Árnason, E. & Halldórsdóttir, K. Codweb: Whole-genome sequencing uncovers extensive reticulations fueling adaptation among Atlantic, Arctic, and Pacific gadids. *Sci. Adv.* **5,** eaat8788 (2019).
2. Solís-Lemus, C. & Ané, C. Inferring phylogenetic networks with maximum pseudolikelihood under incomplete lineage sorting. *PLOS Genet.* **12,** e1005896 (2016).
3. Saitou, N. & Nei, M. The neighbor-joining method: a new method for reconstructing phylogenetic trees. *Mol. Biol. Evol.* **4,** 406–425 (1987).
4. Paradis, E., Claude, J. & Strimmer, K. APE: Analyses of Phylogenetics and Evolution in R language. *Bioinformatics* **20,** 289–290 (2004).
5. Coulson, M. W., Marshall, H. D., Pepin, P. & Carr, S. M. Mitochondrial genomics of gadine fishes: implications for taxonomy and biogeographic origins from whole-genome data sets. *Genome* **49,** 1115–1130 (2006).
6. Teletchea, F., Laudet, V. & Hänni, C. Phylogeny of the Gadidae (sensu Svetovidov, 1948) based on their morphology and two mitochondrial genes. *Mol. Phylogenet. Evol.* **38,** 189–199 (2006).
7. Owens, H. L. Evolution of codfishes (Teleostei: Gadinae) in geographical and ecological space: evidence that physiological limits drove diversification of subarctic fishes. *J. Biogeogr.* **42,** 1091–1102 (2015).
8. Malmstrøm, M. *et al.* Evolution of the immune system influences speciation rates in teleost fishes. *Nat. Genet.* **48,** 1204–1210 (2016).
9. Hughes, L. C. *et al.* Comprehensive phylogeny of ray-finned fishes (Actinopterygii) based on transcriptomic and genomic data. *Proc. Natl. Acad. Sci. U.S.A.* **5,** 201719358 (2018).
10. Tørresen, O. K. *et al.* An improved genome assembly uncovers prolific tandem repeats in Atlantic cod. *BMC Genomics* **18,** 95 (2017).
11. Günther, T. & Nettelblad, C. The presence and impact of reference bias on population genomic studies of prehistoric human populations. *PLOS Genet.* **15,** e1008302 (2019).
12. Sheng, G.-L. *et al.* Paleogenome reveals genetic contribution of extinct giant panda to extant populations. *Curr. Biol.* **29,** 1695–1700.e6 (2019).
13. Kirubakaran, T. G. *et al.* Two adjacent inversions maintain genomic differentiation between migratory and stationary ecotypes of Atlantic cod. *Mol. Ecol.* **25,** 2130–2143 (2016).
14. Kucuk, E. *et al.* Kollector: transcript-informed, targeted de novo assembly of gene loci. *Bioinformatics* **33,** 1782–1788 (2017).
15. Roth, O. *et al.* Evolution of male pregnancy associated with remodeling of canonical vertebrate immunity in seahorses and pipefishes. *Proc. Natl. Acad. Sci. U.S.A.* **117,** 9431–9439 (2020).

16. Nguyen, L.-T., Schmidt, H. A., Von Haeseler, A. & Minh, B. Q. IQ-TREE: A fast and effective stochastic algorithm for estimating maximum-likelihood phylogenies. *Mol. Biol. Evol.* **32,** 268–274 (2015).

17. Roa-Varon, A. & Orti, G. Phylogenetic relationships among families of Gadiformes (Teleostei, Paracanthopterygii) based on nuclear and mitochondrial data. *Mol. Phylogenet. Evol.* **52,** 688–704 (2009).

18. Green, R. E. *et al.* A draft sequence of the Neandertal genome. *Science* **328,** 710–722 (2010).

19. Durand, E. Y., Patterson, N., Reich, D. & Slatkin, M. Testing for ancient admixture between closely related populations. *Mol. Biol. Evol.* **28,** 2239–2252 (2011).

20. Korneliussen, T. S., Albrechtsen, A. & Nielsen, R. ANGSD: Analysis of next generation sequencing data. *BMC Bioinformatics* **15,** 356 (2014).

21. Kelleher, J., Etheridge, A. M. & McVean, G. Efficient coalescent simulation and genealogical analysis for large sample sizes. *PLOS Comput. Biol.* **12,** e1004842 (2016).

22. Juric, I., Aeschbacher, S. & Coop, G. The strength of selection against Neanderthal introgression. *PLOS Genet.* **12,** e1006340 (2016).

23. Schumer, M. *et al.* Natural selection interacts with recombination to shape the evolution of hybrid genomes. *Science* **360,** 656–660 (2018).

24. Matute, D. R. *et al.* Rapid and predictable evolution of admixed populations between two *Drosophila* species pairs. *Genetics* **214,** 211–230 (2020).

25. Tørresen, O. K. *et al.* Genomic architecture of haddock (*Melanogrammus aeglefinus*) shows expansions of innate immune genes and short tandem repeats. *BMC Genomics* **19,** 240 (2018).

26. Kirubakaran, T. G. *et al.* A nanopore based chromosome-level assembly representing Atlantic cod from the Celtic Sea. *G3,* g3.401423.2020 (2020).

27. Ronco, F. *et al.* Drivers and dynamics of a massive adaptive radiation in cichlid fishes. *Nature* **589,** 76–81 (2021).

28. Barth, J. M. I. *et al.* Disentangling structural genomic and behavioural barriers in a sea of connectivity. *Mol. Ecol.* **87,** 449 (2019).

29. Musilova, Z. *et al.* Vision using multiple distinct rod opsins in deep-sea fishes. *Science* **364,** 588–592 (2019).

30. Speidel, L., Forest, M., Shi, S. & Myers, S. R. A method for genome-wide genealogy estimation for thousands of samples. *Nat. Genet.* 1321–1329 (2019).

31. The International HapMap Consortium. The International HapMap Project. *Nature* **426,** 789–796 (2003).

32. Bouckaert, R. R. *et al.* BEAST 2.5: An advanced software platform for Bayesian evolutionary analysis. *PLOS Comput. Biol.* **15,** e1006650 (2019).

33. Ogilvie, H. A., Bouckaert, R. R. & Drummond, A. J. StarBEAST2 brings faster species tree inference and accurate estimates of substitution rates. *Mol. Biol. Evol.* **34,** 2101–2114 (2017).

34. Müller, N. F., Ogilvie, H. A., Zhang, C., Drummond, A. & Stadler, T. Inference of species histories in the presence of gene flow. *bioRxiv.* doi:10.1101/348391 (2018).

35. Malinsky, M., Matschiner, M. & Svardal, H. Dsuite - Fast D-statistics and related admixture evidence from VCF files. *Mol. Ecol. Resour.* **19,** 1655 (2020).

36. Ruegg, K., Anderson, E. C., Boone, J., Pouls, J. & Smith, T. B. A role for migration-linked genes and genomic islands in divergence of a songbird. *Mol. Ecol.* **23,** 4757–4769 (2014).
